# Tracking tau and cellular responses in human iPSC-microglia from uptake to seedable secretion in extracellular vesicles

**DOI:** 10.1101/2025.07.17.664991

**Authors:** Maria Kreger Karabova, Anna del Ser-Badia, Anne Hedegaard, Sam J. Washer, Zeynep Baykam, Darragh P. O’Brien, Iolanda Vendrell, Svenja S. Hester, Roman Fischer, Errin Johnson, Charlotte E. Melia, Teige R.S. Matthews-Palmer, Rishi Matadeen, Alessia Santambrogio, Michael A. Metrick, Michele Vendruscolo, Sophie Keeling, Kimberly Ai Xian Cheam, William A. McEwan, Kenneth S. Kosik, Theresa A. Day, William S. James, Sally A. Cowley

## Abstract

The templated spread of tau aggregates in tauopathies has been attributed to neuron-to-neuron spread, but microglia have also been implicated through mouse studies. Here we examine in detail the uptake, processing, release and seeding of tau using human iPS-derived microglia (iMGL). We show that tau is taken up by iMGL via LRP1 and heparan sulfate proteoglycans, with a role for LRRK2 in LRP1 trafficking, and that phagocytosed fibrils can escape into the cytoplasm. Monomeric tau has minimal effects on iMGL, but recombinant or brain-derived tau fibrils induce a shift towards chemokine and interferon response subtypes, alongside downregulation of homeostatic and MHC genes. Endogenous tau protein is undetectable in iMGL, and monomeric internalised tau is digested to completion, but fibrillar tau is more resistant to degradation and becomes phosphorylated on two specific residues. Finally, fibrillar tau is released by iMGL, visualized within extracellular vesicles by cryo-EM, and can seed tau aggregation in downstream neurons.

## Introduction

In neurodegenerative diseases collectively known as tauopathies, the microtubule-associated protein tau (MAPT) accumulates and propagates across neuroanatomically- connected brain regions in a spatiotemporal manner that correlates with disease progression[1]. This stereotypical propagation of tau was first observed by Braak and Braak in individuals with Alzheimer’s disease (AD)[2], but is known to happen in other tauopathies, including progressive supranuclear palsy (PSP)[3, 4], argyrophilic grain disease (AGD)[5] and Pick’s disease (PiD)[6]. Aside from cell-autonomous mechanisms, a growing body of research implies that intercellular spread of tau occurs in a prion-like manner, whereby misfolded tau templates the misfolding of physiological forms of tau. This cell-to-cell tau propagation was first demonstrated by injecting brain extracts from P301S tau transgenic mice into wild-type human tau-expressing mice[7]. Subsequently, numerous studies have replicated this finding using different forms of tau seeds and experimental approaches, including *in vivo*[8–11] and cell-based assays[12–15].

Key characteristics of prion-like proteins include cellular uptake, templated seeding, and subsequent transfer between cells. These mechanisms have primarily been explored in the context of tau transmission between neurons. For instance, fibrillar tau endocytosis[16] and subsequent seeded tau aggregation in cultured neurons are mediated by the low-density lipoprotein receptor LRP1[17] and heparan sulfate proteoglycans (HSPGs)[18, 19] and are highly dependent on the levels of membrane cholesterol[20]. Tau aggregates can then be trafficked inside exosomes to synaptically-connected neurons[21], a process that is facilitated by neuronal activity, triggering seeded aggregation in neighbouring neurons[22]. However, pathological tau propagation may also involve other brain cell types besides neurons.

Microglia are the major brain-parenchyma resident phagocytes, responsible for maintaining brain homeostasis by constant surveillance and removal of dead cells, redundant synapses and protein aggregates. Therefore, dysregulation of microglial function has been consistently implicated in tauopathy risk and progression[23]. Post-mortem tissue analyses[24], and positron emission tomography (PET) studies of living subjects[25], have correlated proinflammatory microglial phenotypes with pathological tau deposition and cognitive deficits in tauopathies. Importantly, genome-wide association studies (GWAS) have identified genetic risk variants for AD, including TREM2 and CD33, which are exclusively or predominantly expressed in microglia[26], further underscoring the necessity of deciphering the role of microglia in disease progression. However, most functional studies investigating tau uptake, degradation, and spread by microglia, utilise mouse models or immortalized cell lines, which poorly recapitulate human physiology. Mouse microglia can take up tau through shared pathways with neurons (via LRP1 and HSPGs[27]), and through alternative routes by binding to CX3CR1[28, 29], or via Fc receptors when antibody-bound[30, 31]. Following internalisation, tau is targeted for degradation through the endolysosomal pathway[31, 32]. But incomplete tau degradation may lead to the secretion of seeding-competent tau to the extracellular space[33] and into extracellular vesicles (EVs)[34–37]. Indeed, high-resolution cryo-Electron Microscopy (EM) and proteomic analyses have recently shown that straight and paired helical filaments composed of truncated hyperphosphorylated tau are tethered inside EVs isolated from brains of individuals with AD[38]. The exact brain cell types that these EVs come from remains to be elucidated.

Here, we capitalise on the development of human induced Pluripotent Stem Cell-derived microglia models (iPS-microglia-like cells, iMGL, and related iPS-primitive-macrophages, iMac) to explore the molecular mechanisms underlying the processing and spread of tau by human microglia. As with all myeloid cells, microglia are known to be exquisitely sensitive to bacterial endotoxin (lipopolysaccharide, LPS), which binds TLR4 and the myeloid-cell-specific co-receptor CD14. Therefore, reagents for microglial studies, especially *E.coli*-derived recombinant proteins, need to have negligible endotoxin levels. We have therefore optimised the production of endotoxin-free recombinant tau to avoid inadvertent TLR-mediated inflammatory signalling by residual LPS. We use this, alongside human AD brain-derived tau, to challenge iPS-microglia (Fig. 1A). We show that tau internalisation by human microglia is mediated primarily by HSPGs for fibrils, and by LRP1 for monomeric species (with LRRK2 influencing LRP1 trafficking). Human microglia cannot fully degrade fibrillar tau, and their transcriptome shifts towards chemokine and interferon response subtypes. Furthermore, we show that microglia undergo extensive changes to their phosphoproteome upon tau fibril uptake, and can post-translationally modify internalised recombinant tau fibrils at two specific residues.

**Fig. 1:**
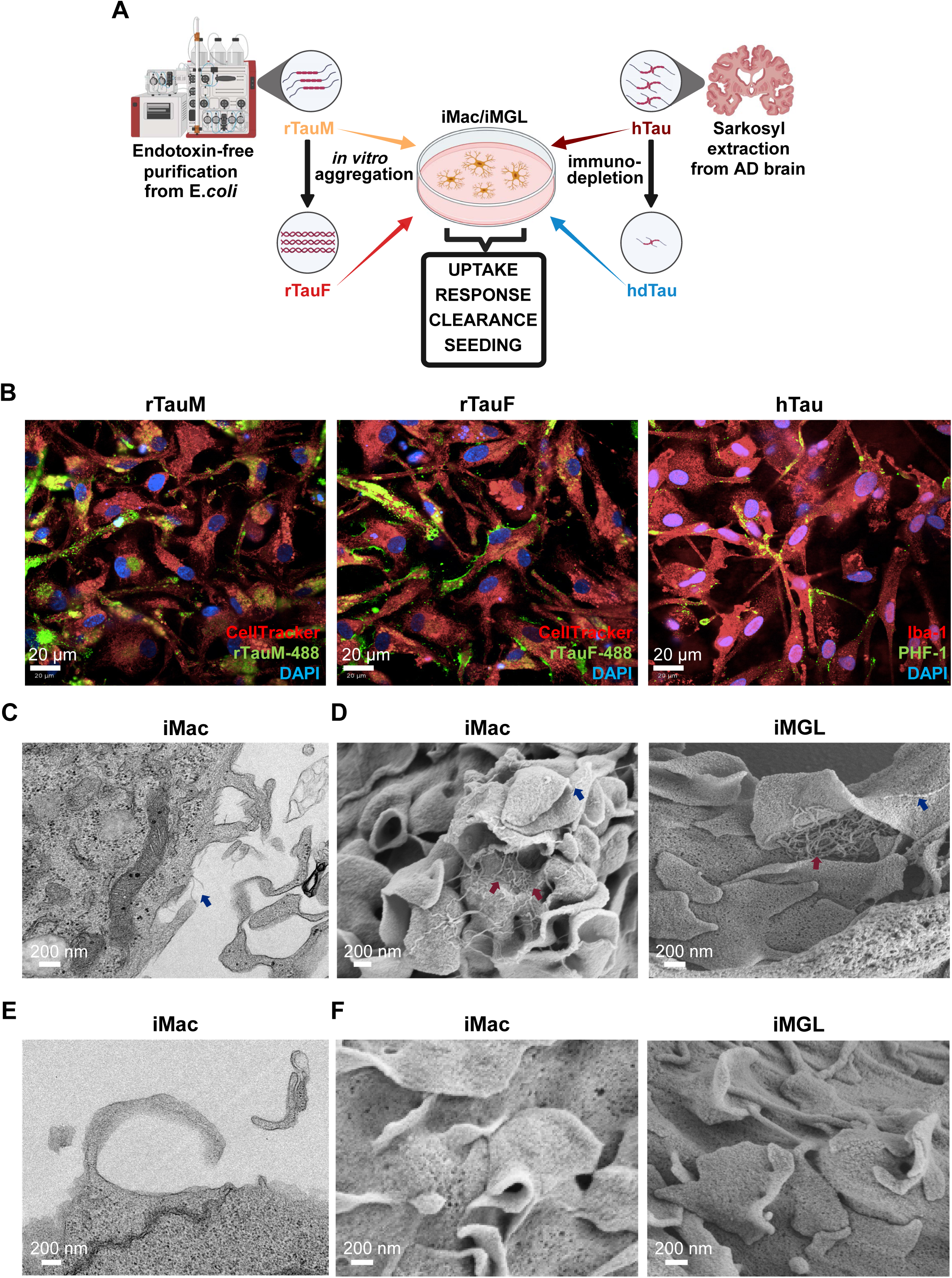
Uptake of monomeric and fibrillar tau by iMGL. **A** Schematic overview of experimental setup. **B** Representative confocal microscopy images of healthy control-derived iMGL following overnight incubation with DyLight 488- conjugated rTauF, rTauM, or hTau (PHF-1). **C** Transmission and **D** Scanning electron micrographs depicting rTauF tethered to the iMac/iMGL plasma membrane (blue arrows) and during uptake (red arrows). **E** Transmission and **F** Scanning (right) electron micrographs of vehicle treated cells, magnification = 69.75 K (iMac) and 40.48 K (iMGL).

Microglia secrete undegraded seeding-competent tau to the conditioned medium, including in EVs, and tau fibrils are clearly visible by cryo-EM as packaged inside these EVs, supporting the role of human microglia in tau spread in tauopathies.

## Results

### iMGL take up tau in monomeric and aggregated form

Tau uptake was initially explored using healthy control-derived iPS-Microglia-Like cells (iMGL) by confocal microscopy (Fig. 1A). Recombinant, full-length 2N4R tau monomer (rTauM) was purified following an initial Triton X-114 on-column wash to remove endotoxin from the lysate before it could strongly associate with tau. Characterisation showed it to have endotoxin levels of <0.01 EU/mL (at a tau concentration of 1 mg/mL) and to be seed- competent when fibrilised with heparin (Supp. Fig. 1A-G). When fed to iMGL, monomer, fibrils (rTauF) and human brain-derived tau (hTau, Supp. Fig. 2A-E) all showed signal overlap with CellTracker-labelled iMGL cytoplasm (Fig. 1B) by immunocytochemistry, with tau fibrils also visible lining the cell periphery. Tau fibrils were also visualised by scanning and transmission electron microscopy on the surface of tau-fibril-fed iPS-Macrophages (iMac) and iMGL (Fig. 1C and D, blue arrows, versus vehicle-treated cells Fig. 1E and F), along with examples of rTauF surrounded by plasma membrane protrusions (Fig. 1D, red arrow).

### LRP1 and HSPGs mediate tau uptake by iMGL

Tau monomer fed to iMac and iMGL colocalised with LRP1 by confocal microscopy, while fibrils colocalised less strongly (Fig. 2A, B). Tau internalisation was significantly reduced by the LRP1 competitive binder sRAP (with less inhibition of fibrils than monomer), and also by LRP1 knockdown using LentiCRISPR (versus an Intergenic region-targeting control gRNA, Fig. 2C and Suppl. Fig. 3A-C). In control cells (vehicle/INTG gRNA), internalised tau was detected in the surface LRP1-negative cell population as a result of receptor internalisation upon tau binding. This effect was significantly lower in cells treated with sRAP or LRP1 gRNA (Fig. 2C righthand graph). Tau binding to iMGL was also significantly blocked by heparin, which binds HSPGs, with tau monomer being inhibited less strongly than fibrils (Fig. 2D). Fractalkine (CX3CL1, which binds to CX3CR1 on myeloid cells) had no significant effect on tau uptake (Supp. Fig. 3D).

**Fig. 2:**
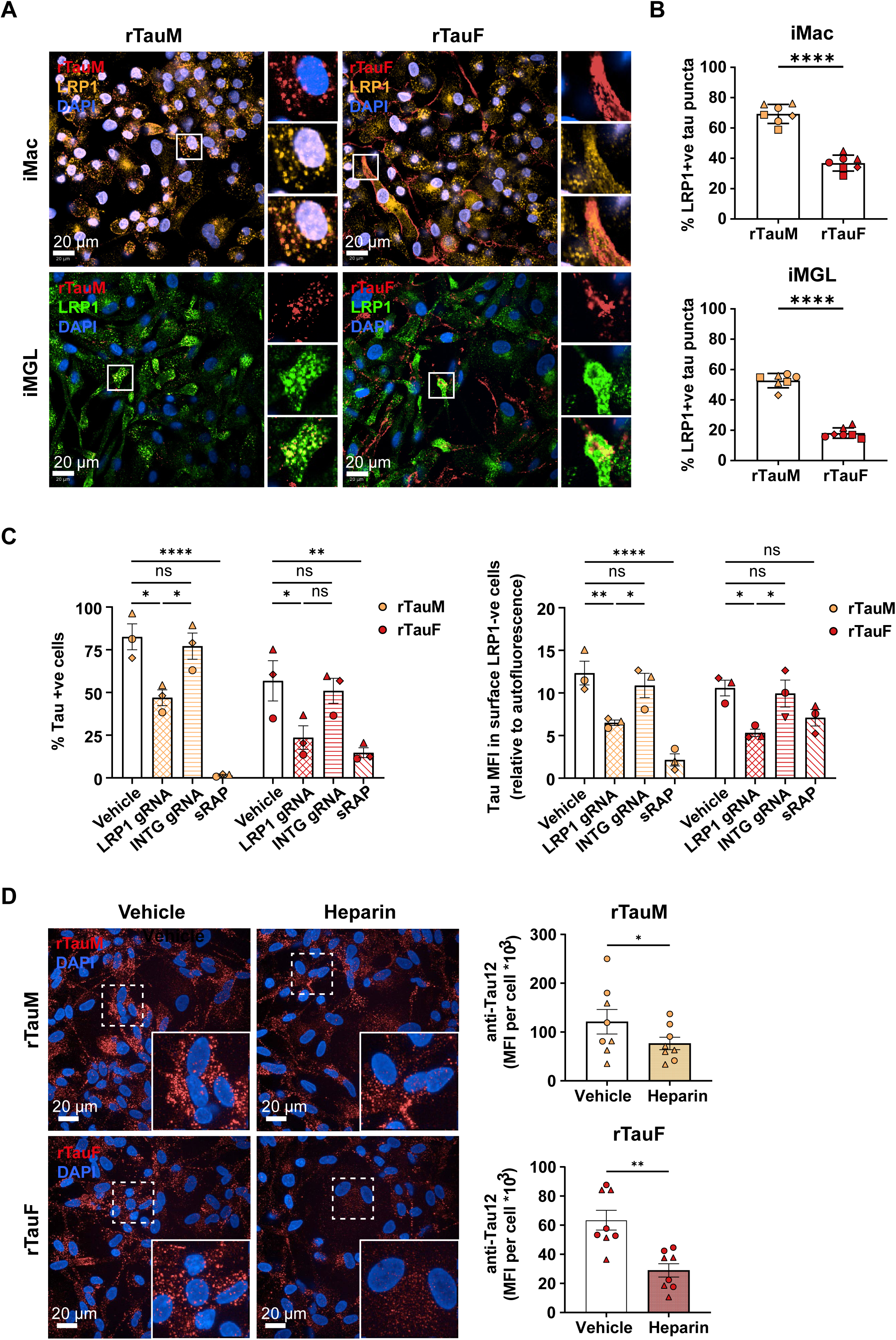
Monomeric and fibrillar tau internalisation by iMGL is mediated by LRP1 and HSPGs. **A** Representative confocal microscopy images of LRP1 colocalising with DyLight 488- conjugated (iMac) or anti-Tau12 (iMGL) immunostained tau monomer and fibrils after 2 h incubation. **B** Quantification of (A), two-way ANOVA with Šídák multiple comparison test, n = 1-2 in 4 control cell lines. **C** Cas9-LRP1 gRNA lenti-transduction of iMac (versus Intergenic region-targeting gRNA INTG), or blockade with sRAP are associated with a decrease in % of tau+ve cells (left graph) and internalised tau levels in surface LRP1-ve cells (right graph, Mean Fluorescence Intensity relative to autofluorescence). Two-way ANOVA with Tukey’s multiple comparisons test, n = 1 in in 3 control cell lines. **D** Representative confocal microscopy images and quantification of Tau12 mean integrated density in iMGL incubated with 10 µg/mL heparin for 1 h, and challenged with 1 µg/mL rTauM or rTauF for 2 h. Data is represented as mean ± SEM of n = 4 in 2 control cell lines. One-way ANOVA with Dunn’s multiple comparison test. \**P* < 0.05, \*\**P* < 0.01.

iMac harbouring the Parkinson’s-associated mutation G2019S in LRRK2 (which increases LRRK2 kinase activity) took up significantly more tau monomer, while LRRK2 knockout had the opposite effect (Supp. Fig. 4A). LRRK2 knockout significantly reduced surface LRP1 levels (Supp. Fig. 4B, C). Upon addition of tau, surface LRP1 decreased significantly in G2019S cells, but accumulated significantly in LRRK2 KO cells, suggesting that LRRK2 KO cells have a reduced ability to traffic LRP1-tau complexes from the surface to the cell interior (Supp. Fig. 4C).

### Tau fibril challenge promotes downregulation of homeostatic and MHC gene expression and upregulation of chemokine gene expression in iMGL

To explore the response of iMGL to tau exposure, we initially looked for secretion of the inflammatory cytokines IL-1β and IL-6 by ELISA. No significant response to 24 h rTauM, rTauF or hTau was observed, although iMGL did respond, as expected, to the positive control stimulation LPS (Supp. Fig 5A,B).

We therefore went on to use a global approach (RNA-seq) to determine what, if any, iMGL responses were induced by tau exposure (Schematic Fig. 3A). iMGL exposed to monomeric tau (rTauM) resulted in very few changes to gene expression versus vehicle control (Fig. 3B), with only five genes downregulated (*HSPG2, IFITM10, COL1A1, COL5A1,* and *APBA3,* log2FC < -0.5, FDR < 0.05) and no genes reaching the threshold for upregulation. A much greater change was observed when iMGL were exposed to fibrillar tau (rTauF, Fig. 3C), with 87 genes downregulated and 96 genes upregulated, including notable downregulation of homeostatic markers *CX3CR1* and *P2RY12.* In contrast, the top upregulated genes were linked to cytokine and chemokine responses, notably *CCL7, CCL2, CCL1, CXCL5.* Gene Ontology analysis of vehicle vs rTauF differentially expressed genes, showed an overrepresentation of chemokine and chemotaxis pathways in the upregulated gene set, and MHC-associated genes in their downregulated counterparts (Supp. Fig. 5B,C, Supp. Data 1,2,3).

**Fig. 3:**
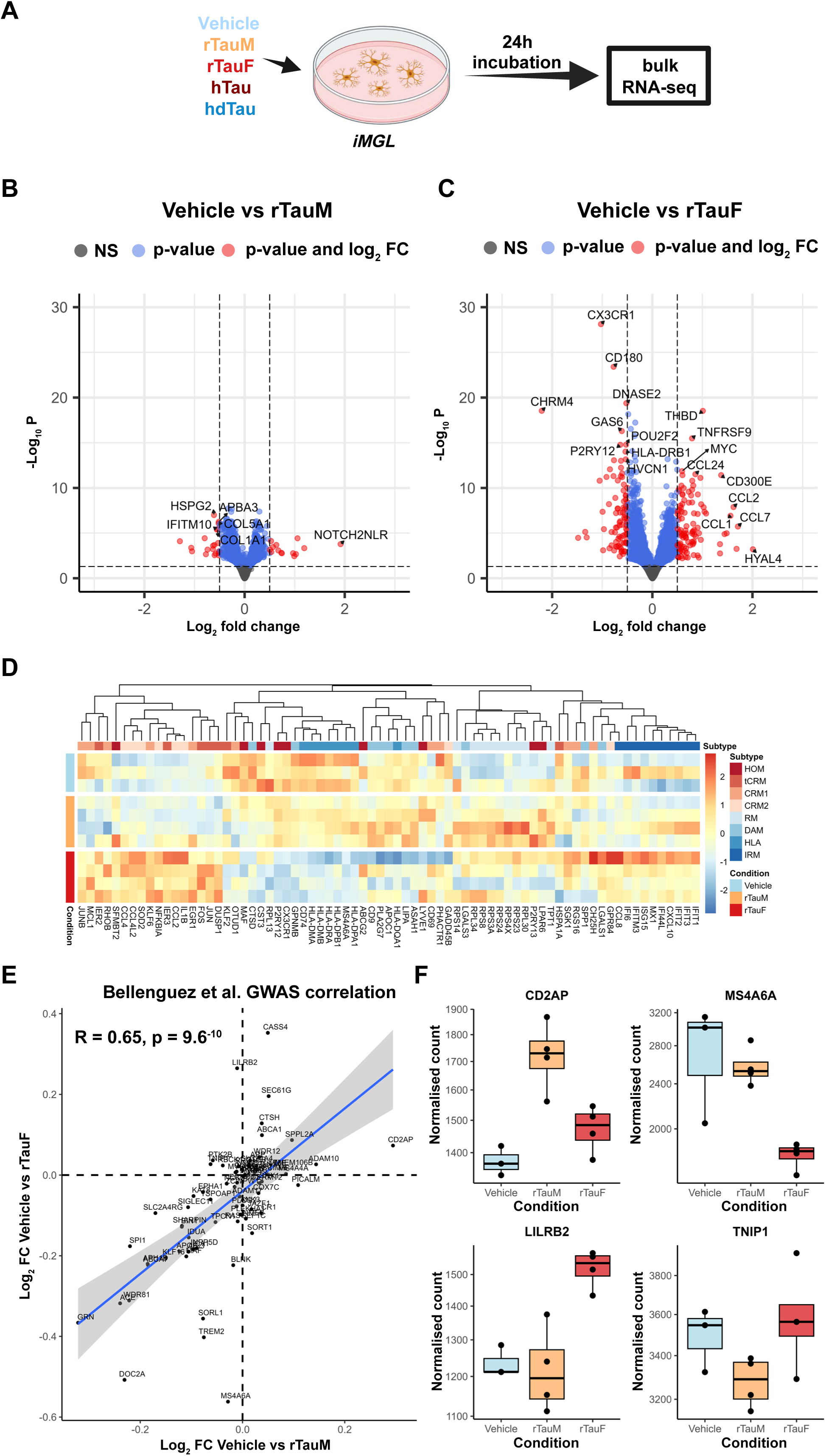
Transcriptomic analysis of iMGL exposed to rTau. **A** Experimental overview of rTau and hTau iMGL exposure for bulk RNA-seq. **B** DESeq2 volcano plot of Vehicle vs rTauM hits. iMGL treated with rTauM show little expression changes when compared to Vehicle control. Thresholds are Log2FC +0.5 or -0.5, and p-value cutoff at 0.05. Note these are non-FDR corrected p-values. **C** DESeq2 volcano plot of Vehicle vs rTauF hits. iMGL treated with rTauF show larger expression changes than that of rTauM. Thresholds are Log2FC +0.5 or -0.5, and p-value cutoff at 0.05. Note these are non-FDR corrected p-values. **D** Heatmap of the marker genes of iMGL subtypes described by Mancuso et al. Normalised expression scaled by gene is shown. **E** Correlation of the Log2FC of AD GWAS genes from Bellenguez et al. across the Vehicle vs rTauM and Vehicle vs rTauF. **F** Example AD GWAS genes showing unique expression changes in only rTauM or rTauF in both directions. *CD2AP,* upregulated in rTauM (Log2FC 0.29, FDR<0.01). *LILRB2,* upregulated in rTauF (Log2FC 0.26, FDR <0.01) *MS4A6A,* downregulated in rTauF (Log2FC -0.56, FDR<0.01), *TNIP1,* trending towards downregulation in rTauM (Log2FC -.0.063, FDR>0.05). Normalised Count data shown, points indicate individual samples.

Next, we explored whether rTau exposure resulted in microglial subtype switching by examining the expression of a panel of known subtype marker genes[39] from xenotransplanted iMGL (Fig. 3D). Expression of genes associated with cytokine response microglia (CRM) and interferon response microglia (IRM) increased following exposure to rTauF, with downregulation of HLA and homeostatic (HOM) markers (mapping to the gene ontology analysis). In contrast, microglia exposed to rTauM had a similar expression profile to vehicle, apart from slight upregulation of ribosomal microglial genes (RM).

Examining expression of the most recently published AD GWAS loci[26] (Fig. 3E), showed a strong positive correlation between vehicle vs rTauF and vehicle vs rTauM log2FC (R=0.65, p = 6.9x10^-6^). Several genes had a strong effect only in rTauM exposure (*CD2AP*, log2FC 0.29, FDR<0.01, *TNIP1*, log2FC -0.063, FDR>0.05), or only in rTauF exposure (*MS4A6A*, log2FC -0.56, FDR<0.01, *LILRB2*, log2FC 0.26, FDR <0.01) (Fig. 3F).

We then examined the effect of exposure of iMGL to tau purified from human AD brain (hTau) to assess shared transcriptional responses between hTau and rTauF. To control for all material present in the AD brain-derived preparation, we also exposed iMGL to the same preparation which had been immuno-depleted to reduce the level of tau (hdTau) (Supp. Fig. 6A-C). Comparing vehicle-exposed iMGL to either hTau or hdTau exposure identified substantial gene changes regardless of tau depletion state (Supp. Fig. 6D). To correct for the non-tau-specific background, we then compared hTau to hdTau to uncover tau-specific iMGL responses. This resulted in 29 upregulated genes (log2FC > 0.5, FDR < 0.05), and 23 downregulated genes (log2FC < -0.5, FDR < 0.05) (Fig. 4A). Genes involved in microglial homeostatic signalling were downregulated (*CX3CR1, P2RY12, OLFML3)* and chemotaxis genes were upregulated (*CCL2,* involved in tissue recruitment of blood monocytes, and *CXCL5*). There is a modest correlation in directionality in log2FC between vehicle vs rTauF and hdTau vs hTau (R=0.34, p<2.2x10^-16^), indicating shared signal regardless of tau origin (Fig. 4B). Notably, the top correlated upregulated genes are all involved in chemotaxis (*CXCL5, CCL2, CD300E, CCL7, CD226),* and the two most downregulated genes (*CHRM4, KIF26B)* in intracellular signalling.

**Fig. 4:**
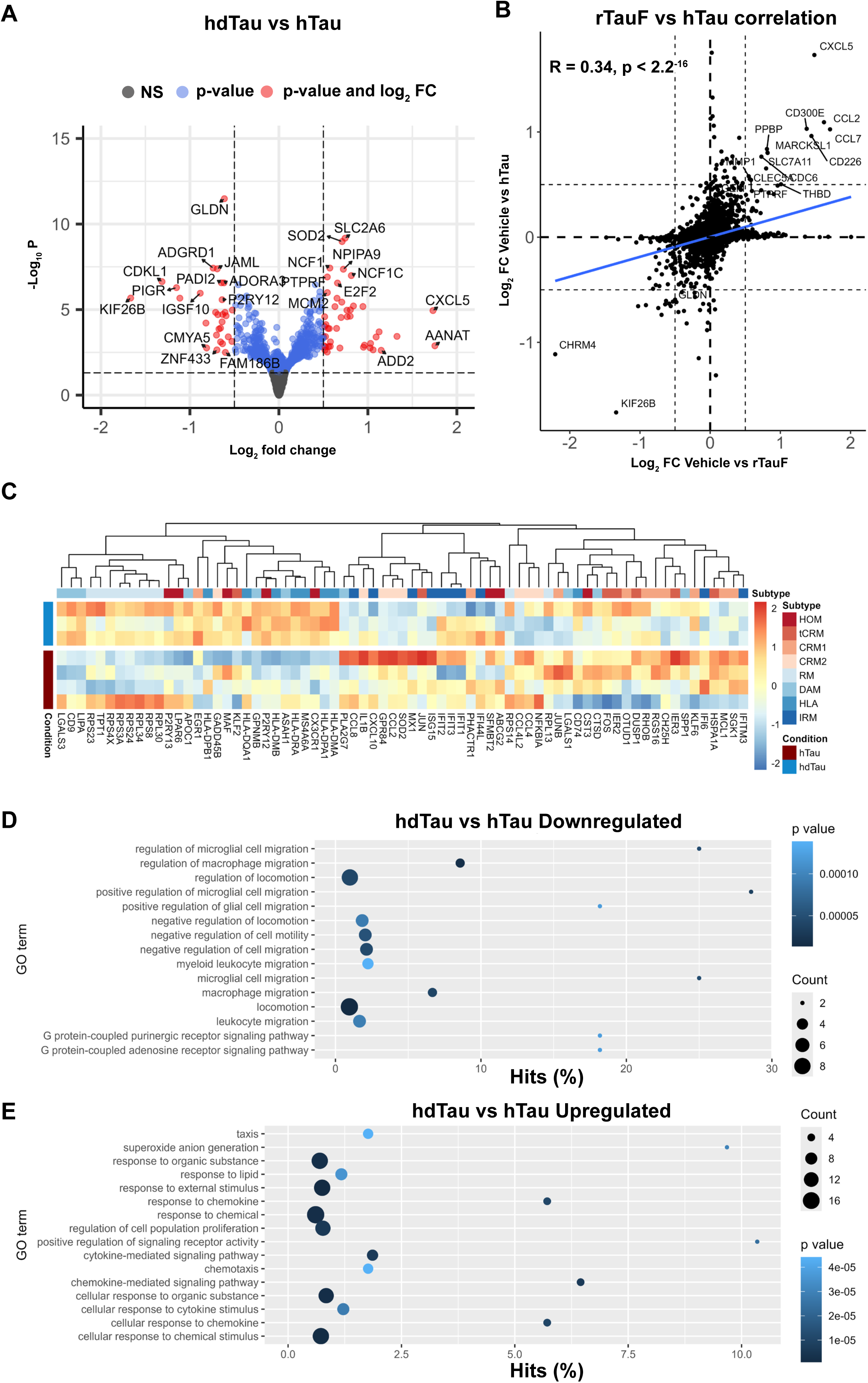
Transcriptomic analysis of iMGL exposed to hTau. **A** DESeq2 volcano plot of hdTau vs hTau hits. iMGL treated with hTau show similar level of expression changes to Vehicle vs rTauF. Thresholds are Log2FC +0.5 or -0.5, and p-value cut off at 0.05. Note these are non-FDR corrected p-values. **B** The Log2FC between Vehicle vs rTauF and hdTau vs hTau show some correlation of gene expression changes regardless of Tau origin. **C** Heatmap of the marker genes of iMGL subtypes described by Mancuso et al. Normalised expression scaled by gene is shown. **D** GOSeq of the downregulated genes identifies enrichment in pathways for negative regulation of chemotaxis, locomotion, and G-coupled protein receptor pathways. **E** GOSeq of the upregulated genes identified enrichment in pathways for chemotaxis, DNA replication/proliferation, and various chemical response pathways.

Examining hTau-challenged iMGL subtypes, hTau showed an upregulation of CRM and IRM subtype genes and downregulation of HOM, DAM, and RM subtypes (Fig. 4C), similar to the subtypes observed for rTauF. Finally, GO analysis for hTau identified similar pathways to rTauF, notably upregulation of chemotaxis-positive genes, and downregulation of chemotaxis inhibitor pathways (Fig. 4D,E).

Together, this shows that while monomeric tau challenge has negligible effects on iMGL gene expression, tau fibrils, whether recombinant or brain-derived, induce downregulation of homeostatic genes, upregulation of chemokine genes and a shift towards chemokine and interferon response subtypes.

### Tau fibrils are degraded poorly by iMGL and can escape into the cytoplasm

To better understand the ability of iMGL to degrade tau, we challenged iMGL with low or high doses of tau (pulse), then removed exogenous tau (by a mild TryplE wash) and assessed the remaining tau content in the cells, in addition to released tau, by ICC and ELISA after defined periods (chase) (Fig. 5A). The majority of rTauM was degraded within 6 h of chase, and relatively little was released into the supernatant. Conversely, rTauF was poorly degraded, with the majority of internalised tau still present in the cells, even at the lower dose and longer chase time, with the amount released into the supernatant increasing significantly with dose, pulse and chase time (Fig. 5B). Protease inhibition significantly increased the amount of rTauF detected in the cells, implying at least some capacity to degrade tau fibrils, likely by lysosomal enzymes (Supp. Fig. 7A,B).

**Fig. 5:**
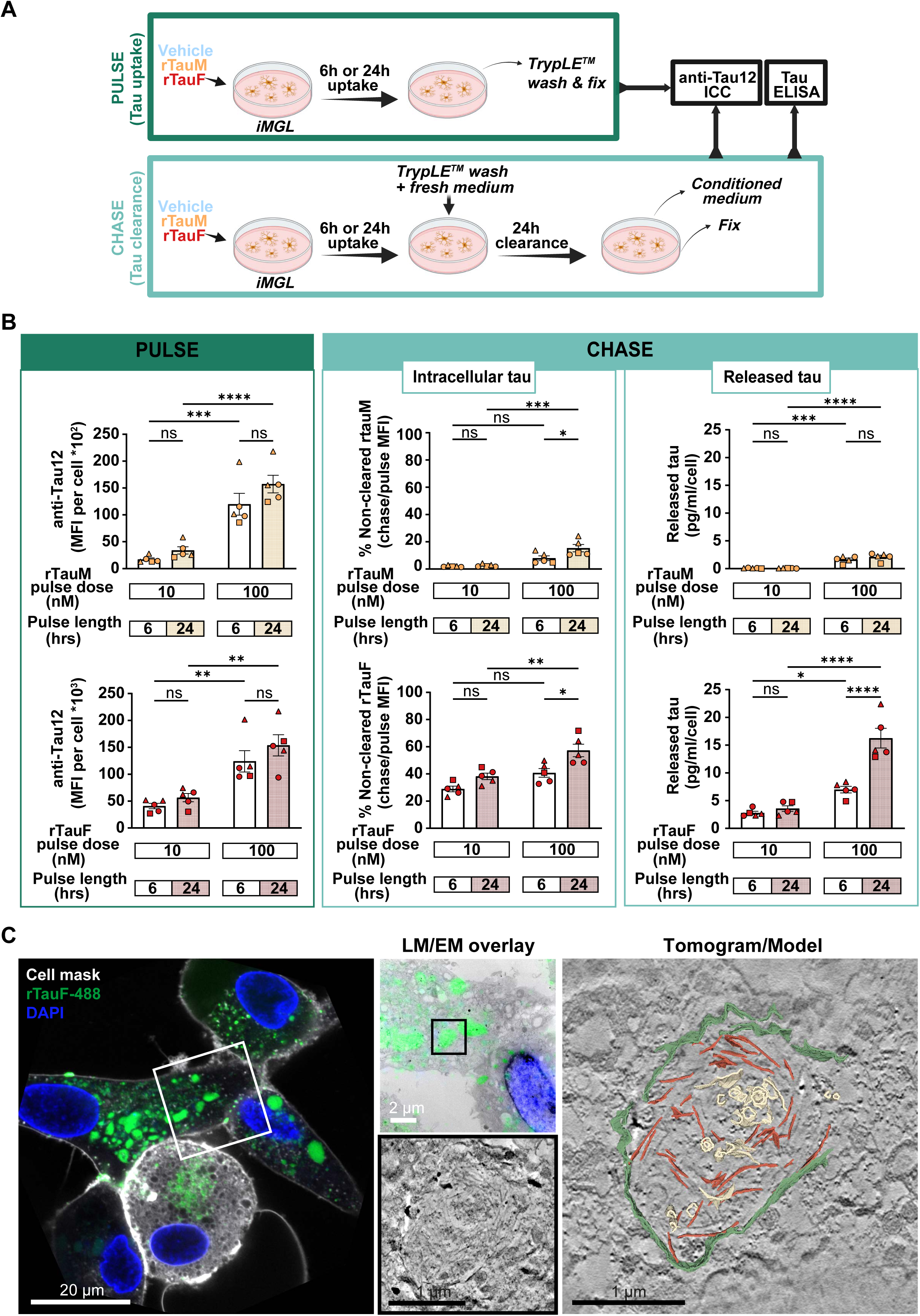
Fibrillar tau escapes degradation and it is freely present in the cytosol. **A** Schematic of tau pulse-chase experiment. **B** rTauM/rTauF internalisation (pulse) and clearance (chase) by iMGL. Intracellular tau levels in pulse and chase were quantified by anti-Tau 12 immunostaining (MFI), relative to the number of cells. The amount of released tau in the conditioned media was determined with total tau ELISA relative to the number of cells. N = 1-2 in 3 control cell lines, two-way ANOVA with Tukey’s multiple comparisons test, mean ± SEM, \**P* <0.05, \*\**P* <0.01, \*\*\**P* <0.001. **C** Correlative light and electron microscopy (CLEM) overlay (left) and region of interest (EM data boxed region) of iMGL incubated overnight (16 h) with Dylight-488-labelled rTauF. Tomogram slice with 3D model (right) of the same region indicates rTauF (red) and membranous structures (yellow) within a partial enclosing membrane (green).

To understand whether internalised tau fibrils remained in the endophagolysosomal system of iMGL, we performed Correlative Light and Electron Microscopy (CLEM), using 488-labelled tau fibrils. Electron tomography of 488-positive regions revealed accumulations of tau fibrils within partial or damaged membrane-bound compartments, suggesting that fibrils can compromise the integrity of their transport vesicles and escape into the cytosol (Fig. 5C, Supp. Fig. 7C, Supp. Videos 1 and 2).

### Tau-fibril-challenged iMGL phosphorylate tau and undergo extensive phosphoproteome remodelling

AD brains have been previously characterized by proteomic and phosphoproteomic analysis to identify biochemical and cellular pathways altered during disease progression[40–43]. However, it is still unclear how the proteome of each brain cell type is modified in AD. Here, using mass spectrometry, we analysed the proteome and phosphoproteome of iMGL treated with rTauM, rTauF, and hTau (Fig. 6A). Samples clustered primarily by cell line rather than cell treatment by principal component analysis (PCA), indicating that the main source of variation in our data arises from the genetic background of each donor (Supp. Fig. 8A). A total of 8,101 proteins were identified in the cell lysates (Supp. Data 4), but no major differences were found when comparing vehicle versus rTauM, rTauF, or hTau conditions, with 17, 12, and 58 differentially expressed proteins (DEPs) found in each, respectively (Fig. 6B, Supp. Data 4). Notably, the absence of tau (MAPT) in lysates from iMGL treated with rTauM indicates the ability of microglia to fully degrade monomeric, but not fibrillar, tau (Fig. 6B).

**Fig. 6:**
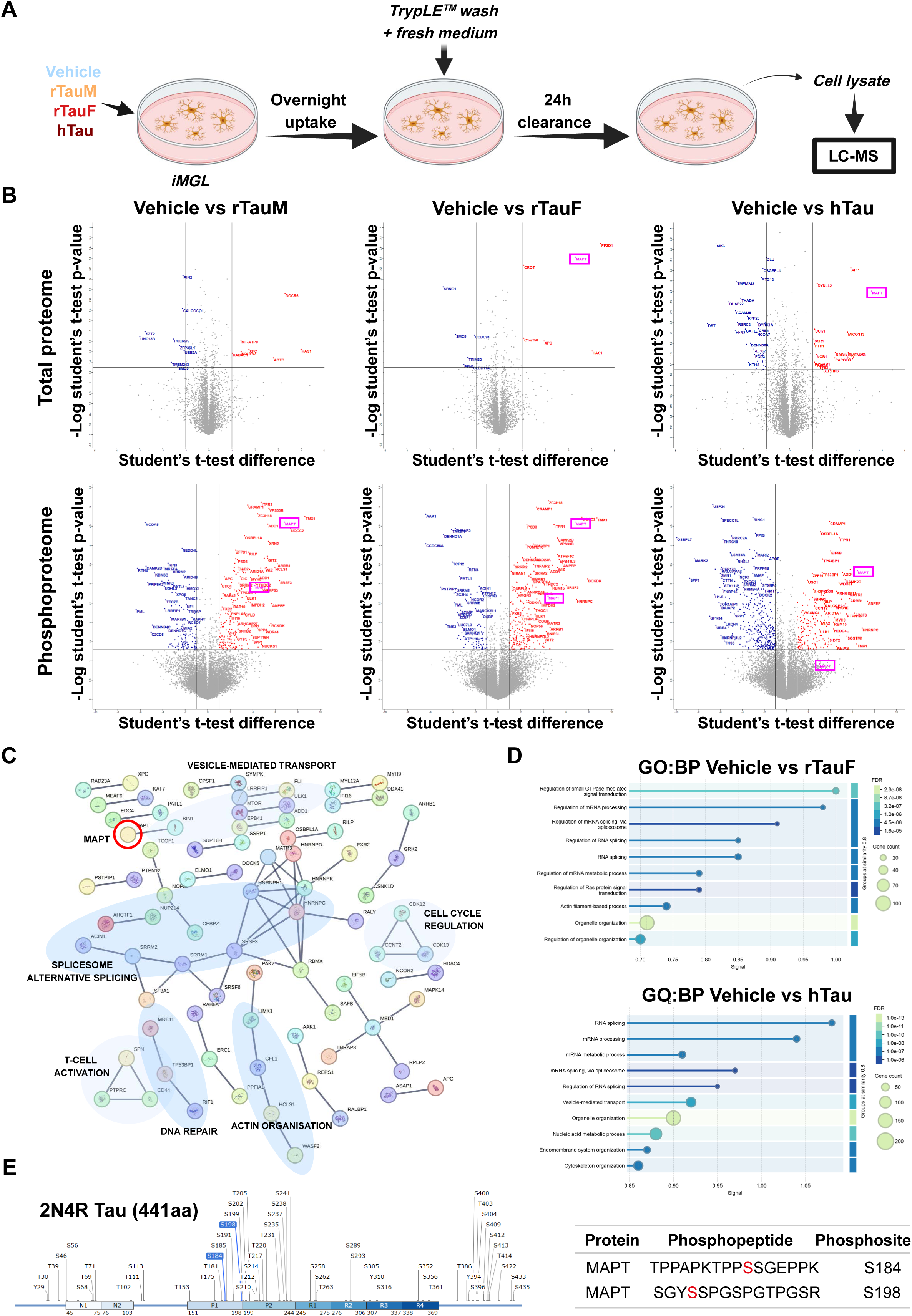
Proteomics and phosphoproteomics of iMGL lysates. **A** Experimental workflow. Cells were treated overnight with rTauM, rTauF or hTau, and allowed to process internalised tau for 24 h. Cell lysates, conditioned medium (CM) and extracellular vesicles (EVs) were collected for proteomic analysis. **B** Differentially expressed proteins (upper panel) and phosphopeptides (lower panel) in iMGL treated with rTauM, rTauF or hTau compared to vehicle-treated cells. Vertical line: fold change above 2; Horizontal line: significant proteins (-logp >1.301 or p<0.05). MAPT highlighted in pink boxes. **C** STRING analysis of differentially expressed phosphopeptides in iMGL treated with rTauF. Only high-confidence interactions (interaction score > 0.9) are displayed. MAPT highlighted with red circle. **D** Classification of differentially expressed phosphopeptides in rTauF and hTau iMGL based on GO biological processes. **E** Map of all 2N4R tau phosphosites, with the upregulated phosphoresidues (S184 and S194) found in iMGL treated with rTauF highlighted in blue (image created with SnapGene). MAPT phosphopeptides sequences are indicated in the table.

Phosphoproteomic analysis of microglia cell lysates identified 43,456 phosphopeptides across all samples (Supp. Data 5), with 329, 308 and 547 differentially expressed phosphopeptides identified in rTauM, rTauF, and hTau samples, respectively (Fig. 6B, Supp. Fig. 8B-D, Supp. Data 5). STRING network and Gene Ontology (GO) biological process (BP) analysis identified functional categories common to all tau-treated conditions studied, including vesicle-mediated transport, actin reorganization, and alternative splicing (Fig. 6C,D, Supp. Fig. 8B,C). Notably, phosphosites associated with actin cytoskeleton remodelling (HCLS1, WASF2), autophagy (MTOR, ULK1), vesicle sorting (BIN1), and mRNA processing (SRSF3, HNRNPC) were upregulated in microglia treated with rTauF and hTau (Fig. 6C).

Tau phosphorylation is highly correlated with tau pathology in AD and other tauopathies[42]. Whether microglia could be contributing to tau phosphorylation remains unknown. Since iMGL do not endogenously express tau, and *E.coli*-derived recombinant tau (rTauF) is not post-translationally modified (Supp. Data 4 and 6), we could observe in our phosphoproteomics dataset that tau residues S184 and S198 were phosphorylated in iMGL treated with rTauF (Fig. 6B,E, Supp. Data 5). These two phosphorylated sites were also found in the human AD tau pooled sample (Supp. Data 3), and in hTau-treated microglia (Fig. 6B, Supp. Data 5).

Together this shows that endogenous tau protein is undetectable in iMGL, and that iMGL can degrade exogenous monomeric tau to completion, but internalised fibrillar tau is not efficiently degraded and is actually phosphorylated in iMGL on two specific residues. It also shows that although the total cellular proteome does not change substantively over the relatively short tau-challenge timeframe studied here, the phosphoproteome shows extensive engagement of cytoskeletal remodelling and metabolic pathways.

### Tau-challenged iMGL secrete tau, including associated with extracellular vesicles

We next followed up on our earlier observation (Fig. 5) that showed secretion of internalised tau species by iMGL. Following 16 h tau pulse and 24 h chase (Fig. 7A), rTauF-challenged iMGL released significantly more tau to the CM than rTauM-challenged iMGL, as assayed by total tau ELISA (Fig. 7B), and MAPT peptides were detected in the supernatant by mass spectroscopy (Fig. 7C, Supp. Data 7). Additional DEPs were identified in the CM of iMGL treated with hTau, notably the upregulation of IL-6 and IL1RN (Fig. 7C, Supp. Data 7).

**Fig. 7:**
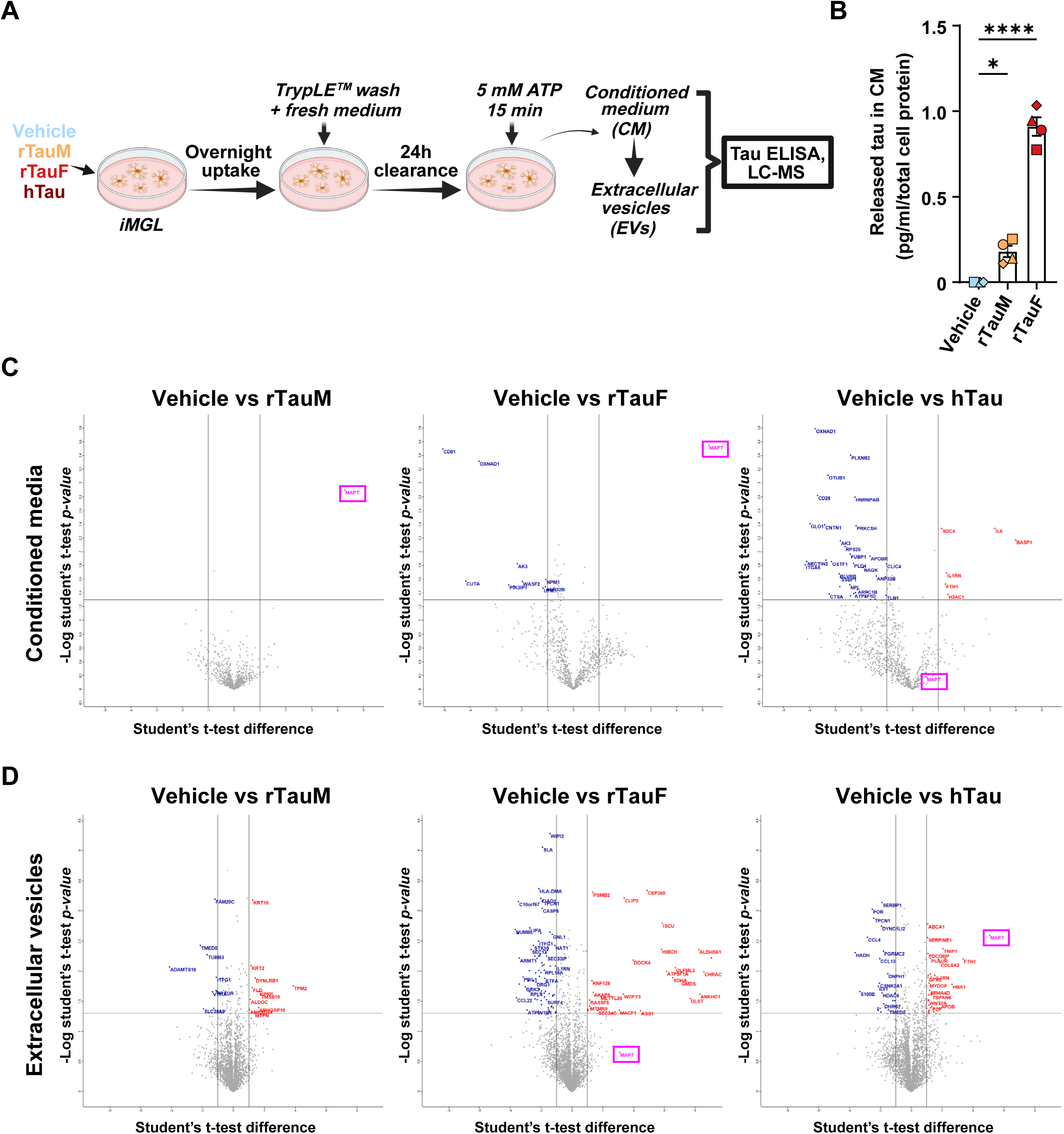
iMGL secrete tau to the conditioned media and inside extracellular vesicles. **A** Experimental workflow. Cells were treated overnight with rTauM, rTauF or hTau, and allowed to process internalised tau for 24 h. iMGL were stimulated with 5 mM ATP (15 min) to enhance the release of EVs to the CM. **B** Tau concentration in the CM of iMGL cells treated with rTauM or rTauF compared to vehicle, measured by ELISA and normalised to total cell protein. Data is represented as mean ± SEM of n = 1 from 4 control cell lines. One-way ANOVA comparing both rTauM and rTauF to vehicle, with Dunnett’s multiple comparisons test. \**P* < 0.0, *\*\*\*\*P* <0.0001 **C** Differentially expressed proteins in the CM of iMGL cells challenged with rTauM, rTauF or hTau. MAPT marked with pink boxes. Vertical line: fold change above 2; Horizontal line: significant proteins (-logp >1.301 or p<0.05). **D** Differentially expressed proteins in isolated EVs from iMGL challenged with rTauM, rTauF or hTau.

We went on to examine the protein content of EVs isolated from the CM of tau-treated iMGL (EV characterisation, Supp. Fig. 9A-D). A total of 3,740 proteins were identified, including canonical EVs markers important for EV formation, targeting and uptake (CD9, CD63, CD81, TSPAN14, MFGE8), cargo sorting (ANXA1/2/4/5/11, HSP90), membrane and vesicle trafficking (RAP1A/B, TSG101), and MHC molecules (HLA-A/B/C, HLA- DRB3) (Supp. Data 5). Markers of cellular organelles not associated with EVs, including calnexin (endoplasmic reticulum), GM130 (Golgi) and cytochrome C (mitochondria) were low in abundance or not detected. rTauM induced only 21 DEPs versus vehicle-treated iMGL, whereas rTauF and hTau induced 109 and 55 DEPs respectively (mostly downregulations). MAPT was not detected in rTauM EVs vs vehicle, was detectable upon rTauF treatment, but only reached significance following the hTau challenge. (Fig. 7D, Supp. Data 8).

Together this shows that EVs expressing typical markers were successfully isolated from iMGL conditioned media and that tau secreted by fibrillar tau-challenged iMGL can be detected in supernatants and associated with EVs.

### Tau fibrils can be visualised packaged inside EVs secreted by tau-challenged iMGL

Having demonstrated that fibrillar tau associates with EVs, we considered whether it may be detectable inside EVs released by iMGL. We performed cryo-electron tomography on purified and concentrated EV samples from iMGL treated with rTauF, comparing them to EVs from vehicle-treated cells. The majority of isolated EVs were within the range expected for exosomes (30-150 nm diameter)[44], with a significant increase in size distribution of EVs from rTauF-treated iMGL vs vehicle (Fig. 8A). EVs isolated from vehicle-treated iMGL did not visually contain filaments (95% of tomograms from the vehicle or TauF-treated datasets were correctly assigned to each group by a blinded independent assessor) (Fig. 8B). rTauF fibrils fed to iMGL are at least 10-fold longer (as conservatively measured from negative stain EM) than within fibril-containing EVs (cryo-ET measurements) (Fig. 8C), suggesting that the packaged fibrils have undergone partial cleavage inside the iMGL. The quantity of fibrils inside EVs varied widely, from singular filaments to numerous and densely packed (Fig. 8D). Fibrils were frequently aligned in bundles, but also occasionally oriented in opposing directions (Fig. 8E). Plotting individual fibril length against the diameter of the EV which contained them, revealed a positive correlation (Fig. 8F). A subset of EVs contained fibrils with visible helical twists, with crossover distances which align well with published cryo-tomography data from recombinantly produced 2N4R tau[45] (Fig. 8G). These results demonstrate that EVs can be a route through which iMGL package and dispose of rTauF. Notably, iMGL seemingly cleave the long rTauF fibrils and form EVs to match this length.

**Fig. 8:**
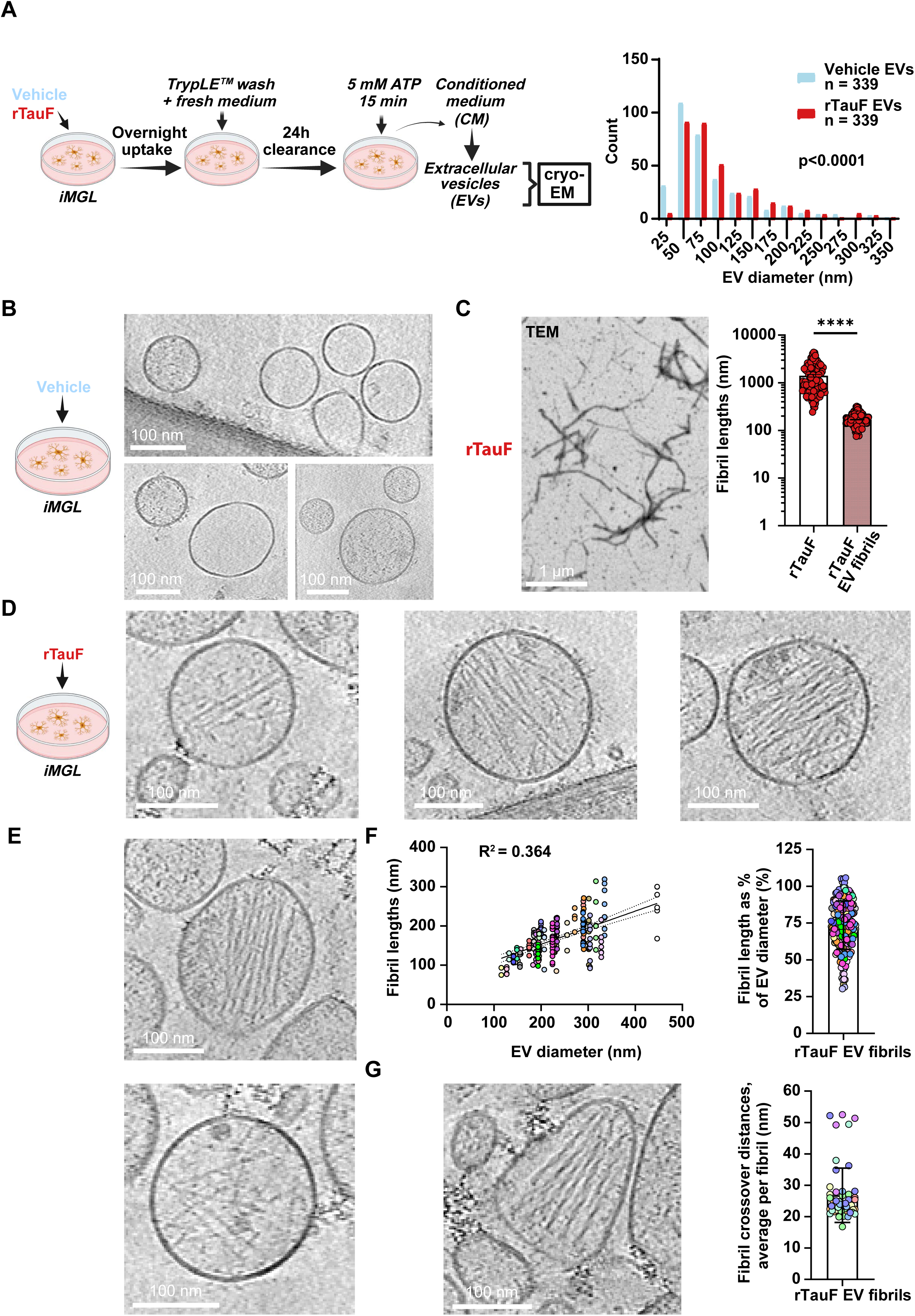
Cryo-ET reveals tau fibrils packaged inside secreted EVs. **A** Experimental overview (left). Diameter distribution of EVs secreted from vehicle or rTauF treated iMGL cells(right). Data is represented as frequency distributions of n = 339 EVs measured from 42 (Vehicle EVs) or 22 (rTauF EVs) tomograms. Kolmogorov- Smirnov test for distribution similarity, *\*\*\*\*P* <0.0001. **B** EVs from vehicle-treated iMGLs. **C** Appearance and length of rTauF used for treatments, compared to fibril lengths measured inside EVs. Data is represented as mean ± SEM of n = 113 fibrils (rTauF) measured across 7 TEM images and n = 223 fibrils (rTauF EV fibrils) measured from 35 EV tomograms. Two-tailed Mann-Whitney test. \*\*\*\**P* < 0.0001. **D** Example images of low, medium and high amounts of fibrils contained in EVs from rTauF-treated iMGLs. **E** Example images of fibrils either aligning inside the EVs to a high degree (top) or having more random orientations (bottom). **F** Fibril length as a function of EV diameter (left), or expressed as a percentage of EV diameter (right), same rTauF EV fibril data as displayed in C. Each dot represents an individual fibril, colour-coded to show which EV they belong to. Linear regression with R^2^ displayed. Data is represented as mean ± SD of n = 223 fibrils measured from 35 EV tomograms. **G** Representative image of the subset of EVs that contain visibly twisted fibrils. Quantification of the helical cross-over distances, averaged per fibril. Each dot-colour represents an individual EV from which several fibrils were often measured. Data is represented as mean ± SD of n = 54 fibrils measured from 8 EV tomograms.

### Fibrillar tau secreted in EVs from iMGL is seed-competent

Finally, we assessed the seeding competency of undigested tau within iMGL cell lysates (soluble and insoluble fractions), as well as in the CM and EVs (Fig. 9A). First, a K11 Real-Time Quaking-Induced Conversion (RT-QuIC) assay was used to determine the presence of seeding-competent tau aggregates in the cell lysates and CM (Supp. Fig. 10A-D). The K11 tau substrate used in the assay is a 4-repeat (4R) tau construct spanning aa residues 244-400 (Supp. Fig. 10B), previously shown to detect 4R and 3R/4R tauopathy-associated pathology and differentiate tau conformer strains[46]. Seeding of recombinant K11 tau substrate by undigested, insoluble rTauF and hTau in SDS iMGL lysates was evident. In comparison, soluble rTauF and hTau in Triton cell lysates as well as CM from iMGLtreated with rTauF and hTau exhibited more residual seeding, with larger ThT amplitude variability. Of note, iMGL samples treated with rTauF vs hTau showed reliably divergent ThT amplitudes consistent with structural differences previously reported by cryo-EM [45] and indicative of faithful propagation of two unique tau amyloid strains through cellular uptake and processing.

**Fig. 9:**
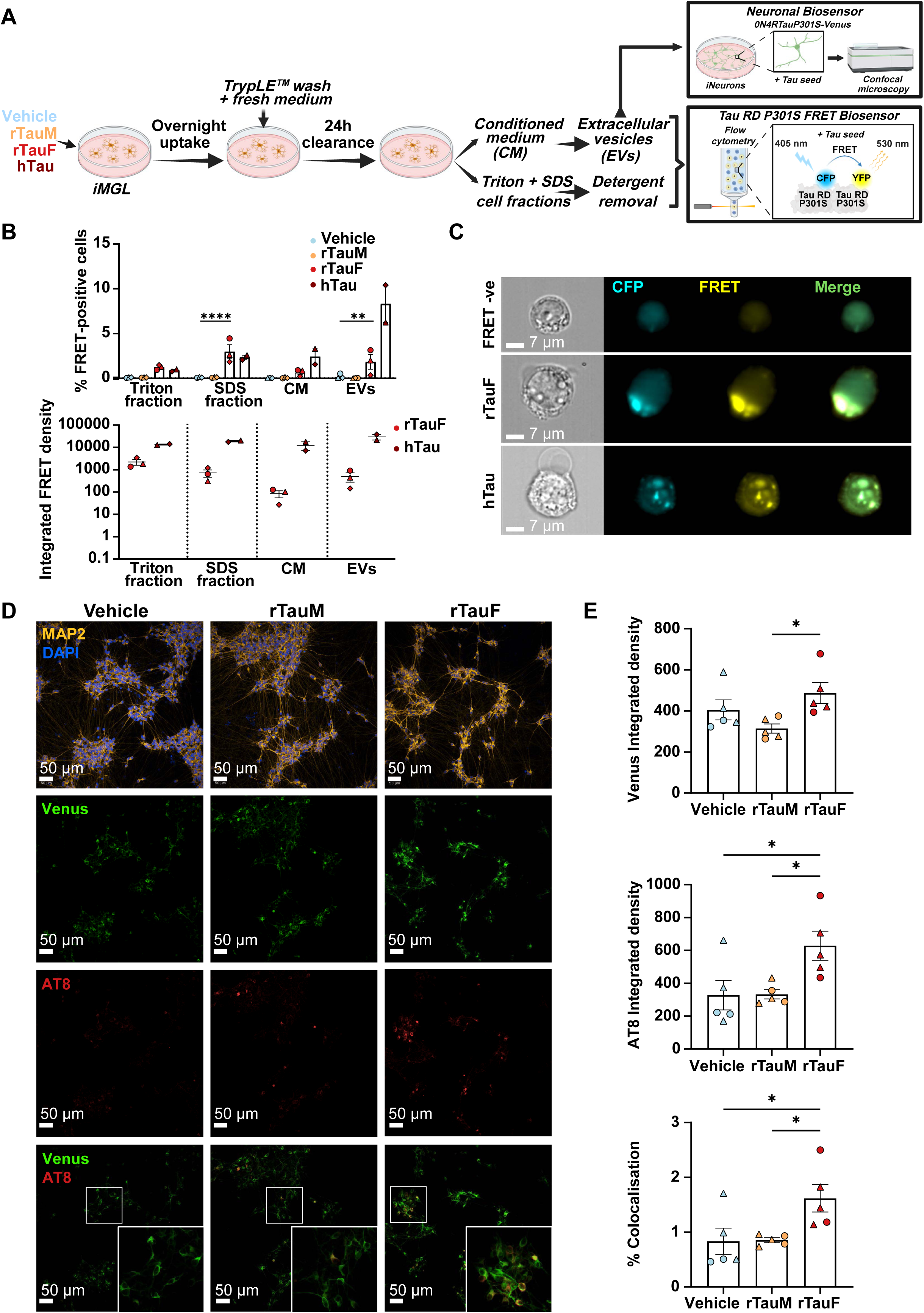
Undigested tau fibrils are present in iMGL cell lysates, CM and EVs and have differential seeding capacities. **A** Experimental workflow. Cells were treated overnight with rTauM, rTauF or hTau, and allowed to process internalised tau for 24 h. Soluble (triton) and insoluble (SDS) cell fractions, CM and EVs were collected and transfected into Tau RD P301S FRET Biosensor HEK cell line. iNeurons expressing 0N4RTauP301S-Venus were also transfected with EVs. **B** Percentage of FRET-positive cells and normalised integrated FRET density in Tau RD P301S FRET Biosensor HEK cells incubated for 48 h with Triton/SDS fractions (1 µg), CM (10 µL) or EVs (2 µg) from iMGL treated with rTauM, rTauF or hTau. Data is represented as mean ± SEM of n = 1 (run in triplicates) from 2-3 control cell lines. Two-way ANOVA with Tukey’s multiple comparison test was used. hTau samples were excluded from the analysis due to small sample size (n = 2). \*\**P* < 0.01, \*\*\*\**P* < 0.0001. **C** Representative images of Tau RD P301S FRET Biosensor HEK cells transfected with EVs isolated from rTauF or hTau-treated iMGL with or without (upper panel) intracellular tau inclusions. Cells were imaged in suspension by Amnis ImageStreamX MkII. **D** Representative images of iNeurons seeded with EVs from vehicle-, rTauM- or rTauF-treated iMGL. **E** Quantification of seeded tau aggregation measured by Venus and AT8 integrated density, and percentage of Venus/AT8 colocalization. Data is represented as mean ± SEM of n = 2-3 (run in duplicates) from 2 control cell lines. One-way ANOVA with Tukey’s multiple comparison test was used. \**P* < 0.05.

We next corroborated our findings using a well-established cell-based assay, the Tau RD P301S FRET Biosensor cell line[47] (Fig. 9A). Tau present in the soluble (Triton lysate) and insoluble (SDS lysate) fractions, CM and EVs from iMGL treated with rTauF and hTau, induced aggregation in the FRET Biosensor cell line, seen as an increase in the percentage of FRET-positive events (Fig. 9B). Since the samples included both total and enriched fractions (EVs), the integrated FRET density (calculated as the product of the percentage of FRET-positive cells and the FRET median fluorescence intensity) was normalized to the total tau levels (pg/mL) detected in each fraction by ELISA (Fig. 9B). ImageStream enabled visualization of intracellular tau aggregate formation in biosensor cells transfected with EVs isolated from rTauF- and hTau-treated iMGL (Fig. 9C).

We proceeded to evaluate the seeding-competency of EVs isolated from iMGL cultures treated with rTauM or rTauF using a more physiologically-relevant novel human neuronal biosensor, consisting of iNeurons expressing human 0N4R tau harbouring the P301S mutation fused to Venus (Fig. 9A). Treated iNeurons were fixed with methanol to remove soluble tau, and Venus and AT8 (pTau S202/T205) antibody signal intensity was quantified, in addition to Venus/AT8 colocalization. EVs from rTauF-treated iMGL induced significant tau aggregation in this neuronal biosensor (Fig. 9D,E).

These results indicate that tau fibrils observed inside iMGL EVs by cryo-EM are seed-competent, including the ability to seed aggregation in human neurons, and could play a role in the cell-to-cell transmission of tau pathology.

## Discussion

Here we examined the uptake, processing, release and seeding of tau in the context of human iPS-microglia, using endotoxin-free recombinant tau and AD-brain-derived tau. We have demonstrated uptake of monomeric and fibrillar tau via LRP1 and HSPGs and have visualized escape into the cytoplasm of internalised fibrils. We have shown that recombinant or brain-derived tau fibrils shift iMGL towards chemokine and interferon response subtypes and induce phosphoproteome remodelling. We have demonstrated that endogenous tau is not detectable in iMGL, that iMGL degrade fibrillar tau inefficiently and can phosphorylate undigested tau on specific residues. Finally, we have shown that iMGL release undegraded tau, including secretion as fibrils within EVs, and that these can seed tau aggregation in downstream cells, including neurons, as summarised in Fig. 10.

**Fig. 10:**
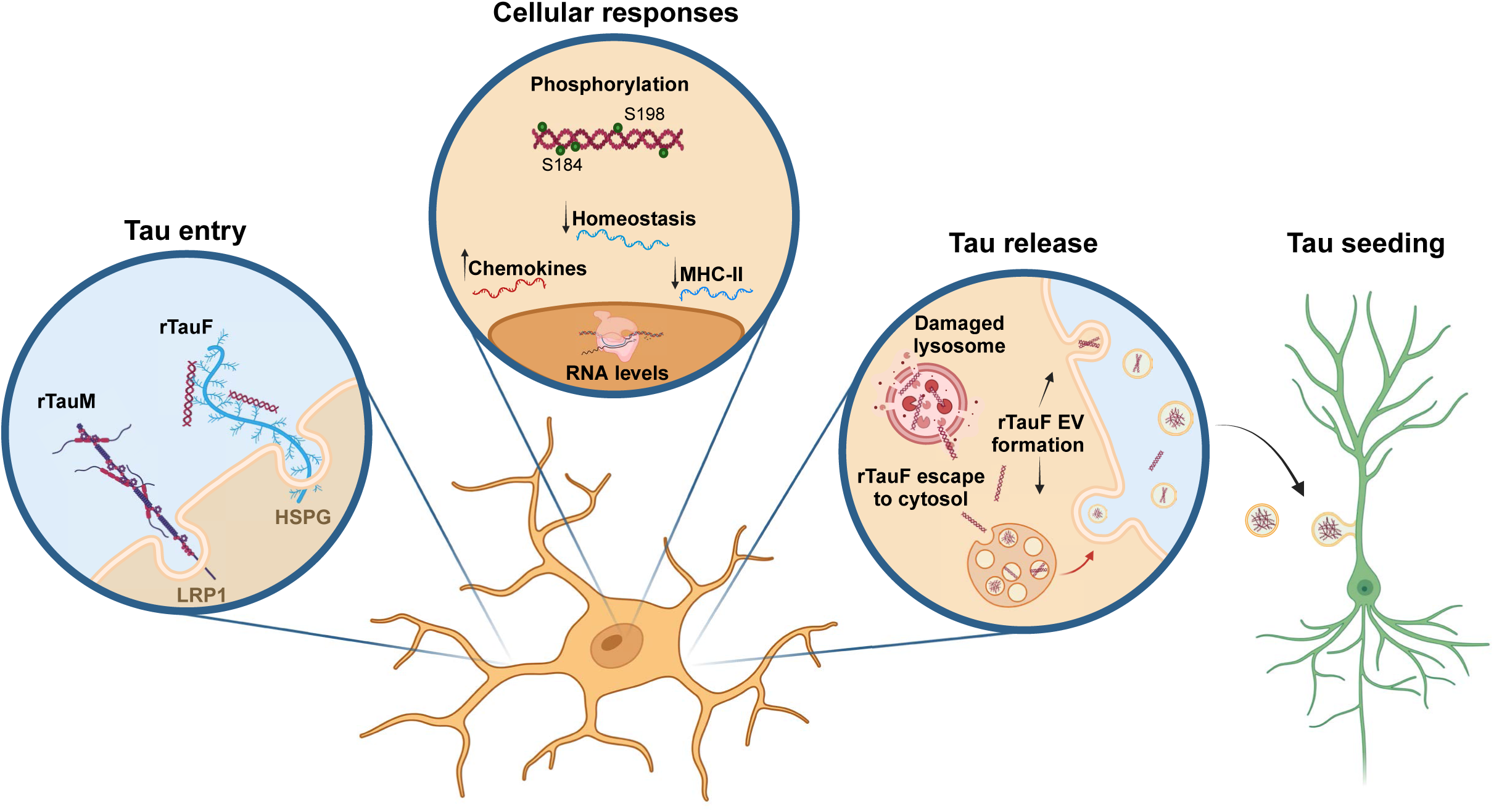
Graphic Summary A summary of tau processing by human iPS-microglia defined in this manuscript. Entry of monomeric tau occurs preferentially via LRP1, whereas fibrillar tau entry is in part facilitated by LRP1 and HSPGs. Cellular responses of microglia upon fibrillar tau internalisation (both recombinant and human brain-derived) include upregulation of chemokine and inferferon pathway genes, downregulation of homeostatic and MHC-II genes and proteomic changes mainly affecting phosphorylation states. In particular, phosphorylation of two serines on MAPT were detected. Evidence of tau release included its detection in conditioned media and in EVs both by proteomics and ELISA. Furthermore, cryo-electron tomography revealed the presence of fibril structures packed into individual EVs and the seeding capacity of this EV-associated tau was verified in two biosensor systems.

Although aspects of this complex process have been demonstrated previously in mouse *in vivo* and *in vitro* models[27–31, 33, 34, 36, 37], it has been unclear whether human microglia play a role in tau propagation. To address this, we have used human iPS-microglia. Many AD-relevant gene variants, particularly those expressed in microglia, are not well represented in rodent models[48], so studying pathophysiological mechanisms in a human microglia model is relevant for generating translationally-relevant information. iPS-derived microglia represent an accurate model for studying human microglia *in vitro,* being terminally differentiated and retaining the genetic background of the donors.

We found that uptake of monomeric tau into iMGL was strongly reliant on LRP1, and only modestly on HSPGs, while uptake of tau fibrils was less dependent on LRP1, and neither form was affected by Fractalkine (CX3CR1 ligand). Our LRP1 and HSPG results concur with recent mouse microglia data [27] and recent human iNeuron data[49], demonstrating that the different uptake pathways for tau monomer versus fibrils are conserved across cell types and species. Intriguingly, we found that the LRRK2 genotype influenced tau uptake via LRP1, with G2019S LRRK2 increasing uptake, and LRRK2 knockout reducing uptake and perturbing surface LRP1 levels. Since LRRK2 phosphorylates specific Rabs to influence vesicle trafficking[50, 51], it is conceivable that LRRK2 influences trafficking of LRP1 to/from the plasma membrane.

Endotoxin (LPS) is a strong inducer of TLR4-mediated signalling, especially in myeloid cells (including microglia), leading to the production of classic inflammatory cytokines. Endotoxin removal from recombinant tau post-purification is ineffective, presumably due to the establishment of strong interactions resulting from charge differences between tau and endotoxin[52, 53]. We therefore developed a tau purification protocol that removes the majority of endotoxin straight after bacterial lysis, through an initial on-column Triton X-114 wash step. This procedure reduces endotoxin in the final purified tau to negligible levels (Supp. Fig. 1). Reviewing the literature reveals that many microglia studies do not state the endotoxin levels in their recombinant tau preps (Supp. Table 9), so it is possible that some reported results are confounded by microglial responses to endotoxin. We detected minimal microglial response to our recombinant monomeric tau, and while recombinant fibrillar tau induced a transcriptomic shift to chemokine and interferon response subtypes, secretion of classic inflammatory cytokines IL-1β and IL-6 were not detected. The concentration of tau we used (1µg/mL, equating to ∼20nM) is likely within the range that microglia might encounter within the interstitial space. Our study therefore clarifies the response of microglia to tau only, without the confounding effect of endotoxin.

CLEM identified regions in tau-fibril-fed-iMGL that contained tau fibrils within partial or damaged enclosing vesicles, suggesting that internalised fibrils can escape into the cytoplasm of iMGL. The fact that we also see specific phosphorylation of internalised recombinant tau fibrils, and can visualise them packaged into EVs, confirms that tau can escape into the cytoplasm. It is in the cytoplasm that tau will most likely be phosphorylated, and also packaged into single-membrane-bound EVs.

Since endogenous tau protein was not detectable in iMGL by ELISA or proteomic analysis, and they have inherently low MAPT transcript levels, our results imply that iPS-microglia do not express readily detectable levels of tau, in contrast to a recent preprint[54]. Tau is expressed at high levels in neurons, where it binds to and stabilises microtubules, creating a relatively stable cellular architecture for maintaining a lifelong synaptic network. Microglia, meanwhile, constantly remodel their cytoskeleton, as their ramifications change over the scale of seconds/minutes, so endogenous tau expression is less likely to be required by microglia.

Recombinant tau was not phosphorylated by the *E.coli* producer cells, making it a simple system on which to observe subsequent phosphorylation events. The two specific phosphorylations (pS184 and pS198) added by microglia to internalised rTauF have been identified previously in AD (but not in healthy) brains [55]. The identity of the kinase(s) responsible merits further exploration. These phosphorylations could have significance, as they are in the proline-rich, microtubule-binding domain, and pS198 is increased at early Braak stages, correlating with formation of early misfolded tau oligomers[56].

Our analyses show that monomeric exogenous tau is effectively removed/degraded following uptake by iMGL, while fibrillar tau is degraded inefficiently. Since cells burdened with excess internalised tau fibrils may need to engage other mechanisms of disposal, we looked at routes of tau secretion following uptake. ELISA showed that iMGL release undegraded tau into the supernatant, and that within the supernatant, tau is associated with EVs. While mouse studies[34, 36, 37] have implicated microglial-derived EVs in the spread of tau, we have extended this to show definitively by cryo-EM that tau fibrils are found packaged within the cytosol of the EVs. We have also quantified several fibril parameters that can help us understand their seeding capacity. Tau fibrils in EVs were approximately 1/10 of the mean length of the input tau ‘meal’, implying partial digestion and/or size-sorting for inclusion within EVs by iMGL, and the potential for production of additional seeding-competent fibril ends than the input fibrils. Importantly, by using recombinant tau, we can be confident that the tau-laden EVs have been generated *de novo* by iMGL, as opposed to being residual EVs carried over from brain tau preps. Finally, a recent study[38] has explored the structure of tau fibrils within AD brain-derived EVs by cryo-EM and observed similar packaging of fibrils into EVs. In that study, it is not possible to attribute EVs to specific cell-type origins, but the fact that we see very similar structures from iMGL-EVs argues that in AD brains, microglia are likely contributing seeding-competent tau-fibril-laden EVs to the EV pool.

## Methods

See Supplementary Materials for extended methods and reagent tables.

### iPSC culture and ethics statement

hiPSC lines from four healthy donors (one also gene-edited to knockout LRRK2), and three Parkinson’s patients harbouring a LRRK2 G2019S allele, were used. See Supp. Table 1 for line details, and Supp.Table 2 for which lines were used in which experiments. Human induced pluripotent stem cell (hiPSC) lines used in this study were previously derived from dermal fibroblasts from donors recruited as part of the Oxford Parkinson’s Disease Centre / EU IMI programme StemBANCC, using Cytotune Sendai reprogramming viruses (A16517 ThermoFisher). SFC840-03-03, SFC841-03-01, SFC856-03-04, SFC832-03-06 and SFC855-03-06 have all been published previously[51, 57–60] and are deposited in EBiSC (STBCi026-A, STBCi044-A, STBCi063- A). SFC833-03-05 is deposited in EBiSC (STBCi005-B) and its characterisation data is available on hPSCReg. SFC840-03-03 LRRK2-/- D10 has been edited using CRISPR/Cas9 to knockout both alleles of LRRK2, has been published previously[51] and is also deposited in EBiSC (STBCi026-A-1). KOLF2.1S was also generated using Cytotune as part of HipSci and has been published previously[61]. iPSC lines were curated in the James & Lillian Martin Centre for Stem Cell Research (Sir William Dunn School of Pathology, University of Oxford), as QC’d frozen banked stocks and cultured in mTeSR1 (#85850, StemCell Technologies) or OXE8[62] on Geltrex (#A1413302, Invitrogen) at 37°C, 5% CO2 with minimal subsequent passaging (as clusters using 0.5 mM EDTA (#15575-020, Invitrogen)) to maintain karyotypic integrity, as previously described[59].

Ethical approval for the derivation and use of iPSC lines with the prefix SFC was previously obtained with written informed consent by donors that specifically stated that their skin biopsies would be used for the derivation of pluripotent stem cell lines (Ethics Committee: National Health Service, Health Research Authority, NRES Committee South Central, Berkshire, UK, REC 10/H0505/71). KOLF2.1S was obtained under material transfer agreement from the Wellcome Sanger Institute, Cambridge, UK (REC Reference 09/H0304/77, covered under HMDMC 14/013 and REC reference 14/LO/0345). All experiments were performed in accordance with UK guidelines and regulations and as set out in the REC.

### Differentiation of macrophage precursors and macrophages (iMac) from iPSCs

iPSCs were differentiated along a primitive myeloid route using our previously developed serum- and feeder-free protocol[63]. This differentiation pathway has been shown to be c-myb-independent[64], implying a primitive ontogeny that approximates to the embryonic/fetal origins of primitive macrophages, including microglia.

Briefly, iPSCs were lifted with TryplE (#12604013, Gibco), and seeded into Aggrewell 800 plates (#34815, STEMCELL Technologies) at 4x10^6 cells per Aggrewell in OXE8 medium (with 10 μM ROCK inhibitor Y-27632, #1201029, Abcam, on day of seeding) supplemented with 50 ng/mL BMP4 (#PHC9534, Peprotech), 50 ng/mL VEGF (#PHC9394, Peprotech), and 20 ng/mL SCF (#130-096-695, Miltenyi Biotec) to promote embryoid body formation directed to mesoderm and hemogenic endothelium. After daily feeding for 5-7 days, EBs were transferred to T175 flasks (‘factories’) and cultured with 25ng/mL IL-3 (# PHC0033, Invitrogen) and 100ng/mL M-CSF (# PHC9501, Invitrogen) to promote primitive myeloid differentiation. Non-adherent primitive macrophage precursors were harvested weekly between weeks 5 and 12 of the factory lifetime, for final differentiation.

For final differentiation to primitive generic macrophages, precursors were cultured for 7 days in macrophage medium[62] containing 100 ng/mL M-CSF (Supp. Table 3) at a density of 8x10^4^ / cm^2^ on standard tissue culture plates (Corning) or imaging plates (Perkin Elmer).

### Differentiation of iPSC-microglia-like (iMGL) cells from macrophage precursors

iPSC-microglia were differentiated from the above myeloid differentiation cultures as described by[65]. Briefly, harvested precursors were cultured for 14 days (incorporating 3 x 50% medium changes) in ITMG microglia medium (Supp. Table 3), containing 100 ng/mL IL-34, 25 ng/mL M-CSF and 10 ng/mL GM-CSF to aid microglia survival, and 50 ng/mL TGFβ1 to promote microglia identity and maturation.

### Recombinant tau production

Recombinant full length 2N4R tau (rTauM) expressed in *E.coli* was purified by sequential column chromatography that included an initial on-column Triton X-114 step[66] to deplete endotoxin from the lysed material (Supp. Fig.1A, B), to less than 0.01 EU/mL (at 1 mg/mL tau concentration). Recombinant tau fibrils (rTauF) were generated from rTauM by shaking in heparin. Fibrilization was confirmed by ThT assay and EM. Fibril toxicity was assessed (Supp. Fig. 2D) and seeding assays confirmed seedability (Supp. Fig. 1C-F). See Supp. Methods and Supplementary Tables 7 and 8.

### *In vitro* synthetic tau fibril assembly

After confirmation of purified human 2N4R tau monomers by intact mass spectrometry (Supp. Fig. 1B), aggregation was induced by heparin at a 4:1 tau:heparin ratio and 2 mM DTT for 11 days at 37°C with orbital shaking at 250 rpm. Tau fibril concentration was determined by Pierce BCA Protein Assay Kit - Reducing Agent Compatible due to the presence of β-mercaptoethanol and DTT in the sample. The formation of fibrils was evaluated by electron microscopy and Thioflavin T assay (Supp. Fig. 1C,D), and seedability was assessed by 4R tau RT-QuIC and cell-based seeding assays (Supp. Fig. 1E,F). Toxicity in iMGL was assessed with resazurin viability assay (Supp. Fig. 2D).

### Enrichment of tau PHFs from human AD brains and Immunodepletion of hTau

Brain-derived tau (hTau) was enriched from Alzheimer’s patient brains obtained from the London Neurodegenerative Diseases Brain Bank at King’s College London and QCed by EM and Western blot (Supp. Fig 2A-C), and toxicity assessed (Supp. Fig 2E). The treatment dose of 0.25 µg/mL was chosen as this conferred no toxicity to iMGL after 24 h incubation. See Supp. Methods

### Immunodepletion of hTau

Immunodepletion of tau from the hTau sample was performed using the Dynabeads ® Antibody Coupling Kit (#14311D, Life Technologies), following the manufacturer’s protocol.

In short, two preparations of 5 mg of beads were washed and then mixed with either 50 µg of mouse IgG1 antibody (control depletion, #SC-3877, Santa Cruz) or 50 µg of HT7 antibody (tau depletion, #MN1000, Invitrogen), achieving a coupling of 10 µg antibody / mg of beads. The mixtures were incubated on a roller overnight at 37°C. The next day, antibody-coupled beads were washed sequentially in 800 µL of HB and LB buffers, supplemented with 0.1% Tween 20 (#P7949, Sigma-Aldrich). Beads then underwent two quick washes in SB buff, followed by a 15 min wash while on a roller at RT and final storage in SB buffer at 25 mg/mL. For depletion, 5 mg of beads (control and tau-antibody coupled) were each mixed with 1 µg of hTau and left on a rotating wheel overnight at 4°C. Beads were removed from immuno-depleted solutions using a magnet and retained to run on a gel as a control.

### Tau, pharmacological and LentiCRISPR treatments of iMac/iMGL

Unless otherwise indicated, iMac/iMGL were incubated with 1 µg/mL human recombinant 2N4R tau monomer (rTauM) or heparin-aggregated fibrils (rTauF), or 0.25 µg/mL human AD brain-derived tau (hTau). A 2.5% TryplE^TM^ incubation was performed for 1 min[67] followed by 1x PBS wash was used to remove uninternalised tau. For tau uptake experiments, cells were preincubated with 500 nM sRAP (#153996, Ximbio), 10 µg/mL heparin (#H3393, Sigma-Aldrich) or 5 ng/mL FKN (#300-31, PeproTech) for 1h before tau treatment (Supp. Table 4). For tau degradation experiments, cells were incubated with a cocktail of protease inhibitors (50 µM Leupeptin, 50 µM Pepstatin A, and 50 µM E64d) or 25 µM MG-132 for 24 h. All incubation times are indicated in the figure legend.

Three LRP1 CRISPR/Cas9 lentiviral vectors were prepared to knockdown LRP1. The guide RNAs (gRNA) sequences targeted exon 12, 13, and 38 of LRP1 (Supp. Table 3). In addition, eight intergenic control (INTG) CRISPR/Cas9 lentivectors, predicted not to affect LRP1 or other protein-coding sequences, were prepared as a negative control of DNA damage response. Golden Gate Assembly via *BsmB*I was used to clone gRNA oligos into pLentiCRISPRv3 backbone, a novel version of pLentiCRISPRv2 (Addgene, #52961) with a modified gRNA scaffold (Washer SJ et al. 2025, in press). Plasmids were transformed into chemically competent Stbl3^TM^ *E*. *coli* (ThermoFisher Scientific, C737303) for amplification and plasmid purification. Lentiviral production was carried out using HEK293T, JetPRIME^®^ transfection reagent (Polyplus, 101000046), 5.5 μg of psPAX2 lentiviral packaging plasmid DNA, 4.4 μg of pMD2.G lentiviral envelope plasmid DNA, and 9 μg of pooled LRP1 gRNA plasmids or pooled INTG gRNA plasmids. Virus was collected at 48 and 72 h and concentrated by ultracentrifugation. Virus transduction was carried out during plating of myeloid precursors, in differentiation medium supplemented with 4 ug/mL of polybrene (Sigma, TR-1003-G), together with 1:400 dilution of Vpx packaged in Virus-Like-Particles (VLP) (pSIV3+_Vpx plasmid, Cosset lab, Inserm, France). The human immunodeficiency virus type 2 (HIV-2)- and simian immunodeficiency virus (SIV)- encoded accessory protein Vpx counteracts the cytoplasmic nucleotide depletion by SAM Domain and HD domain-containing protein 1 (SAMHD1), thus improving lentiviral transduction efficiency in SAMHD1-expressing myeloid cells such as macrophages and microglia[68].

### Immunocytochemistry and Flow Cytometry

Cells were fixed with 2% PFA for 10 min at RT, washed with PBS, and permeabilized with 0.1% Triton-X100 for 10 min at RT. Cells were incubated with blocking buffer (10% donkey serum, 5% BSA and 0.01% sodium azide in PBS) for 1h at RT and incubated overnight at 4°C with primary antibodies diluted in blocking buffer (Supp.Tables 5 and 6). After PBS washes, cells were incubated with secondary antibodies diluted in blocking buffer for 1h at RT, and with DAPI diluted in PBS for 10 min at RT (Supp. Table 5). Cells were washed with PBS and imaged using the automated spinning disk confocal Opera Phenix High Content Screening System (Perkin Elmer). Image analysis was performed with Columbus Image Data Storage and Analysis System (CambridgeSoft).

### Transmission electron microscopy (TEM)

The formation of in vitro aggregated recombinant tau fibrils was assessed by negative stain TEM. Tau monomer and fibril preparation were diluted 1:20 in 25 mM HEPES (# H0887, Sigma-Aldrich), pH 7.1. 10 μL of each sample was applied to freshly glow-discharged (Pelco EasiGlow) 300-mesh carbon-coated copper grids (#C267, TAAB Laboratories) for 2 min. After removing excess sample with filter paper, grids were washed once with ddH2O, negative-stained with 2% uranyl acetate (#R1260A, Agar Scientific) for 20 sec, and air-dried.

TEM was also used to assess the uptake of tau fibrils by iMac. Cells were differentiated on glass coverslips inserted into a 24-well tissue culture plate (#3524, Costar). Following overnight incubation with vehicle or 1 μg/mL of rTauF, cells were fixed with pre-warmed 2.5% glutaraldehyde (#R1020, Agar Scientific) and 2% PFA in 0.1 M PIPES buffer (#P6757, Sigma-Aldrich-Aldrich) pH 7.2, for 1 h at RT. After five washes with 0.1 M PIPES buffer, cells were incubated with 50 mM glycine (#G7126, Sigma-Aldrich) in 0.1 M PIPES buffer for 15 min at RT, then washed again with 0.1 M PIPES buffer. Secondary fixation was carried out with 1% osmium tetroxide (#O001/1, TAAB Laboratories) + 1.5% potassium ferrocyanide (#223111000, Acros Organics) in 0.1 M PIPES buffer at 4°C for 1 h. Samples were then washed 5 times with ddH2O, stained with 0.5% uranyl acetate (#R1260A, Agar Scientific) overnight at 4°C in the dark, then washed again twice with ddH2O, 10 min each time while being protected from light. Consecutive, 10 min long, ice-cold ethanol incubations (30%, 50%, 70%, 80%, 90% and 95%) on ice were used to dehydrate the samples. A final 20 min incubation in 100% dry ethanol was repeated twice. After epoxy resin infiltration (#AGR1140, Agar 100-Hard epoxy resin, Agar Scientific), coverslips were removed from the wells, inverted onto BEEM capsules (#AGG360-1, Type 00, Agar Scientific) filled with fresh 100% resin and blocks were polymerised for 24 h at 60°C. Ultrathin (90 nm) sections of the resin-embedded cells were obtained with a Diatome diamond knife on a Leica UC7 ultramicrotome. Individual sections were mounted onto 200-mesh carbon-coated copper grids (#C101, TAAB Laboratories), stained with Reynold’s lead citrate (#1963, Reynolds) for 5 min at RT, washed with five droplets of degassed ddH2O, and air-dried. All images were acquired using a Gatan OneView camera on a FEI Tecnai T12 transmission electron microscope operated at 120 kV.

### Scanning electron microscopy (SEM)

SEM was used to visualise iMac/iMGL following rTauF incubation. As with the TEM protocol, cells were differentiated on glass coverslips inserted into 24-well tissue culture plates (#3524, Costar), then incubated overnight with vehicle or 1 μg/mL of tau fibrils. Primary fixative of 2.5% glutaraldehyde in 0.1 M PIPES buffer, pH 7.2 was added for 1 h at RT for cross-linking. Cells were rinsed three times with 0.1 M PIPES buffer, then fixed with 1% osmium tetroxide in 0.1 M PIPES buffer for 1 h at 4°C. After three washes with ddH2O, samples were dehydrated in increasingly more concentrated ethanol (50% - 70% - 90% - 95%) for 5 min each, then three times in 100 % ethanol for 10 min each. Cells were then chemically dried using Hexamethyldisilazane (HMDS, #440191, Sigma-Aldrich) as follows: 1:1 100% ethanol:HMDS for 3 min, followed by two consecutive 2-min incubations with pure HMDS, after which the HMDS was removed, and the samples were left to dry overnight. Samples were sputter coated with ∼15 nm of gold using a Quorum Technologies Q150R ES coating unit. Images were captured with a Zeiss Sigma-Aldrich 300 Field Emission Gun Scanning Electron Microscope (FEG-SEM) operated at an accelerating voltage of 2kV.

### Correlative light and electron microscopy (CLEM)

iMGL, previously fed with 488-labelled rTauF, were fixed in 4% formaldehyde and 2.5% glutaraldehyde in 0.1 M PIPES buffer pH 7.2 for 1 h at room temperature. Cells were then stained and processed into resin as described in the TEM section above. The resulting sample block was trimmed to the grid co-ordinate noted during light microscopy imaging that contained cells of interest. Sections of 200 nm were cut from the resin blocks using a Leica UC7 Ultramicrotome and collected onto formvar-coated 3 mm copper slot grids (Agar). The sections were then post-stained with lead citrate and imaged using a JEOL JEM-1400Flash 120kV TEM equipped with a Gatan Rio camera or JEOL JEM-2100Plus 200kV TEM equipped with a Gatan OneView camera. Tilt series were collected at 200 kV over 116-120° around the regions of interest at 1° increments using SerialEM (4.0.26).

TEM images of the regions of interest were overlaid with the corresponding confocal z plane using linear transformations in Adobe Photoshop 2020. Tilt series alignment and tomogram reconstruction by weighted back-projection were carried out in IMOD (4.11.25), resulting tomograms were binned (bin2) and filtered (gaussian filter sigma 1.5) in ImageJ (Fiji 1.54f). Images stacks were segmented using Microscopy Image Browser (ver. 2.91) and visualised with 3D Slicer (5.8.0 r33216).

### Protein quantification

#### Protein sample collection

For total protein extraction from cells, cultures were washed once with ice-cold 1x PBS, then lysed directly with appropriate amounts of ice-cold Radioimmunoprecipitation (RIPA) buffer (#89901, ThermoFisher Scientific) supplemented with cOmplete^TM^ ULTRA Tablets, Mini, EDTA-free, EASYpack Protease Inhibitor Cocktail (#5892791001, Roche) and Pierce^TM^ Phosphatase Inhibitor Mini Tablets (# A32957, ThermoFisher Scientific). Cell lysates were scraped into LoBind^TM^ tubes (#15178344, Eppendorf) and centrifuged at 21,000 g for 30 min at 4°C. Cleared lysates were snap-frozen in liquid nitrogen and stored at -80°C for further use.

#### Cell-surface biotinylation

Cell surface proteins were extracted using Pierce^TM^ Cell Surface Biotinylation and Isolation Kit (# A44390, ThermoFisher Scientific). The manufacturer’s protocol was adapted as follows. Adherent cells were washed once with BupH^TM^ PBS at RT and biotinylated with 2 mL of 1x EZ-Link^TM^ Sulfo-NHS-SS-Biotin for 10 min at RT. After two washes with ice-cold BupH^TM^ Tris Buffered Saline (TBS), cells were scraped into 1 mL of TBS, transferred to Eppendorfs, and centrifuged at 500 g for 5 min at 4°C. Supernatants were discarded. Cell pellets were lysed on ice for 30 min with 200 μL of kit-provided lysis buffer, supplemented with Halt Protease and Phosphatase Inhibitor Cocktail, EDTA-free (100x) (#78441, ThermoFisher Scientific), then centrifuged at 21,000 g for 5 min at 4°C. Biotinylated proteins in the clarified supernatant were captured by 1 h RT incubation with NeutrAvidin^TM^ Agarose in 1:1 ratio. The captured protein-resin complex was washed four times before final protein elution with 50 μL of kit-provided elution buffer, supplemented with 10 mM Dithithreitol (DTT, #43816, Sigma-Aldrich). Eluted protein samples were snap-frozen in liquid nitrogen and stored at -80°C until required.

#### Sample quantification

Protein concentration in all cell lysate and EV samples was determined with Pierce^TM^ BCA Protein Assay Kit (#23227, ThermoFisher Scientific), diluting the sample in appropriate buffer where necessary to remain within the standard curve.

To assess total protein contents of purified EVs, samples were lysed using 10x RIPA buffer (#9806, Cell Signalling) due to the small volumes (<100 μL). 10x RIPA was diluted to 1x by adding appropriate amounts to quantified volumes of EV suspensions after which samples were briefly vortexed and incubated for 30 min on ice. EV samples were lysed immediately prior to experimentation.

#### Western blot

Levels of total and cell-surface proteins of interest in cell lysates, purified extracellular vesicles and tau samples were analysed with western blot. Samples were mixed with NuPAGE^TM^ 4x LDS sample buffer (#NP0007, Invitrogen) and NuPAGE^TM^ 10x Sample Reducing Agent (#NP0009, Invitrogen), unless eluted in buffer supplemented with DTT (#43816, Sigma-Aldrich). Samples were then heated at 70°C for 10 min. A total of 25 μL of sample containing a minimum of 10-50 μg of protein was loaded onto Novex^TM^ WedgeWell^TM^ 8-16%, Tris-Glycine pre-cast mini gels (# XP08165BOX, Invitrogen), along with 5 μL of Precision Plus Protein Dual Color Standards (#161-0374, Bio-Rad), and run in Novex^TM^ Tris-Glycine SDS Running Buffer (#LC2675, Invitrogen) at 180 V for ∼45-60 min.

Following the electrophoresis, gels were rinsed with ddH2O and 1x Trans-Blot Turbo Transfer Buffer (#10026938, Bio-Rad) before semi-dry protein transfer to a Low-Fluorescence PVDF transfer membrane (#22860, ThermoFisher Scientific), using the Mixed Molecular Weight (1.3 A, 25 V, 7 min) pre-programmed settings on a Trans-Blot Turbo transfer machine (Bio-Rad). Membranes were blocked with iBind^TM^ Flex blocking solution (#SLF1020, ThermoFisher Scientific) for 1 h at RT and probed with relevant primary antibodies, diluted in iBind^TM^ Flex (see Table 6), overnight at 4°C. After three washes with 1x PBS supplemented with 0.1% Tween® 20 (PBS-T, #P7949, Sigma-Aldrich) membranes were further stained with relevant, LI-COR fluorophore-conjugated secondary antibodies (see Table 5) diluted in iBind^TM^ Flex supplemented with 1:200 10% SDS (#75746, Sigma-Aldrich). Unbound antibody was removed with six 1x PBS-T washes. Blots were visualised using the Odyssey Sa Infrared Imaging System (LI-COR Biosciences). Image Studio Lite open-source software v5.2 was used for densitometric analysis.

#### Enzyme-linked immunosorbent assay (ELISA)

Levels of intracellular tau and tau released to conditioned medium were quantified using Invitrogen^TM^ Tau (Total) Human ELISA kit (# KHB0041, ThermoFisher Scientific). Cytokine secretion was quantified using Human Uncoated IL-1β and IL-6 ELISA Kits (# 88-7261- 88and #88-7066-88, ThermoFisher Scientific). Protocols were followed as per the manufacturer’s instructions. Total protein was extracted and quantified from cells following vehicle or tau treatment, as described above. Conditioned medium for tau or cytokine quantification was centrifuged at 400 g for 5 min at 4°C to remove any floating cells and cell debris, aliquoted to 96-well V-bottom plates, parafilmed, and stored at -80°C until required.

After thawing, sample dilution was optimised to 1:100 for tau quantification in cell lysates, 1:10 for tau quantification in supernatants, and 1:10 and 1:100 for cytokine quantification in supernatants. Absorbance (450 nm) was measured using a Spectramax M5e Multi- mode Microplate reader (Molecular Devices). Background reading was subtracted from all values. Tau or cytokine concentration in samples was calculated by interpolation from standard curve (sigmoidal, 4PL, X is concentration) in GraphPad Prism v9.0. Values above and below the range of the standard curve were excluded. If multiple dilutions of a sample were assessed, values presented are an average of the dilutions.

#### EV purification

To enhance the release of EVs, iMGL cells were treated with 5 mM ATP for 15 min, added directly to the culture media. The conditioned media was then collected and cell lysates processed as described under the section: “Protein quantification”. The conditioned media underwent a series of differential centrifugation steps following[69] with minor alterations to purify mixed-size EV populations.

In short, the first centrifugation at 500 g for 5 min pellet any floating cells, the second at 2000 g for 20 min pellet dead cells and the third at 10,000 g for 30 min pellets cell debris. At centrifugation steps 1-3, the supernatant was retained and the pellet discarded, and to finally pellet the EVs, the conditioned media was ultracentrifuged at 100,000 g for 90 min in either re-usable 3.5 mL polycarbonate tubes (#349622, Beckman Coulter), or single-use 13.2 mL ultra-clear polypropylene tubes (#344059, Beckman Coulter), depending on the starting volume. Small volumes were processed in a Beckman Optima MAX-XP tabletop centrifuge using a TLA 100.3 fixed-angle rotor, and larger volumes in a Beckman Optima XPN-80 using a SW41Ti swinging bucket rotor. Samples were often frozen down at -70°C after the second spin and the protocol continued at a later date. After the last ultracentrifugation step, the supernatant was poured off and the tubes kept upside down for ∼1 min to let leftover supernatant run off and concentrate the EVs. For experiments where high concentrations of EVs were required (see section: “Cryo-electron tomography”), the pellet was re-suspended without further addition of PBS, otherwise, the EV-containing pellet was re-suspended in 50-100 µL of PBS. To enhance re-suspension, care was taken to wash around the bottom of the tube during re-suspension, and tubes were kept on a rocker overnight at 4°C before being transferred to protein LoBind^TM^ tubes (#10316752, Eppendorf) and stored at -70°C until further experimentation.

#### Nanoparticle tracking analysis (NTA)

The size distribution of EVs was measured using a NanoSight LM10 (Malvern Instruments) equipped with a 405 nm laser.

Samples of purified EVs were diluted in filtered PBS to achieve an appropriate concentration for NTA, 1:500 gave a particle concentration of ∼10^9^ particles/mL. Approximately 1 mL of the diluted suspension was injected into the sample chamber via a syringe from which it was introduced into the flow cell. Particle tracking commenced with the following settings: infusion-rate = 50, temperature = 26°C, framerate = 25 FPS, slider shutter = 1300, slider gain = 512. Each measurement was captured over a 60-sec duration and repeated five times prior to averaging. The size distribution was calculated using the Stokes-Einstein equation, which relates the diffusion coefficient to particle size.

#### Viability assays

##### Resazurin assay

Resazurin (#199303-5G, Sigma-Aldrich) was used to determine the cytotoxicity effect of drugs, rTauM and rTauF on iMac/iMGL. At the end of the pharmacological or tau treatment, cells were incubated with 10 µg/mL resazurin for 2 h at 37°C, 5% CO2. Viable cells with intact mitochondria would reduce the non-fluorescent blue resazurin to red fluorescent resorufin (Ex: 530-560 nm, Em: 590 nm), which was detected on the SpectraMax M5 microplate reader using the SoftMaxPro software v5. The amount of resorufin produced is proportional to the number of viable cells.

##### LDH assay

The cytotoxicity of the human AD brain tau seed preparation was assessed using a lactate dehydrogenase (LDH) detection kit (#426401, Biolegend), following the manual for “Assay using cell supernatant”. In short, macrophage precursors were plated at 32,000 cells/well in 96 well-plates and differentiated to iMGL. At DIV13 iMGL cells received a full media change in preparation for hTau treatment to ensure equal volume in each well. The hTau seeds were pre-mixed in 10 µL/well of iMGL media at 10x final concentrations to achieve 0.1, 0.25, 0.5, 0.75 a 1 µg/mL in test wells and added directly to the existing iMGL media in each well to avoid tau sticking to the plastic. When handling the hTau preparation, LoBind plasticware was used throughout. At 23.5 h post-treatment, wells designated for the “max LDH release” control condition were exposed to 10% triton-x to lyse the cells and incubated at 37°C for 30 min until harvest of the supernatant at 24 h. Vehicle-control wells contained untreated iMGL and background control wells contained iMGL media. Supernatants were centrifuged at 500 g for 5 min in a V-bottom plate to pellet cells and debris, then moved to a 96 well-plate for immediate assaying, following the kit instructions. Absorbance was read at 490 nm on a Spectramax M5e Multi-mode Microplate reader (Molecular Devices) and the averaged background subtracted from all values.

#### RNA-seq analysis

Raw sequencing reads were trimmed of adapter sequences using Trim-Galore before quality control with multiQC[70]. Transcripts were quantified using Kallisto (v0.44.0) with default options, with the additional command *“bootstrapping”* set to 50[71]. The paired-end sequencing reads for each sample were mapped to the human reference transcriptome (GRCh38.p14) which combined all cDNA sequences. The transcript abundances were imported and summarised to gene-level counts using R library “tximport”[72].

Data was prefiltered to include only genes with counts >10 in 3 or more samples. This reduced the dataset to 15,132 genes for the recombinant analysis, 15,090 for the brain tau analysis. Differentially expressed genes (DEGs) between the vehicle vs rTauM, vehicle vs rTauF, and hdTau vs hTau was performed using DESeq2 (v1.42.1) using the Wald test, LFC shrinkage was performed using the “apeglm” method. A gene was considered a DEG at FDR < 0.05 with a Log2FC > 0.5 (upregulated) or < -0.5 (downregulated)[73]. Gene ontology analysis was performed using GOSeq (v1.54.0)[74]. Gene lists for subtyping iMGL were taken from single cell Xenotransplanted samples from[39]. The list of AD GWAS hits was extracted from[26].

Code is deposited in GitHub: https://github.com/S-Washer/Karabova_2025_Tracking_tau_and_cellular_responses_in_microglia_RNAseq

#### Flow Cytometry

Cells differentiated on 24-well tissue culture plates (#3524, Corning) and treated according to experimental design were lifted by 10 min incubation with StemProTM AccutaseTM (# A1110501, Gibco) at 37°C, 5% CO2. Lifted cells in suspension were transferred to a 96-well V-bottom plate (#611V96 and #642000, ThermoFisher Scientific) and kept at 4°C or on ice throughout all steps, unless stated otherwise. All centrifugation was carried out at 400 g for 5 min.

For live-cell flow cytometry analyses of phagolysosomal proteolysis and tau internalisation, cells were spun, washed with 1x PBS, spun again, and resuspended in 40 μL/well of ice-cold Invitrogen^TM^ Live Cell Imaging Solution (LCIS) (# A14291DJ, ThermoFisher Scientific) in preparation for acquisition.

For cell surface marker staining, cells were lifted into ice-cold 1x PBS, counted, spun, and resuspended in ice-cold FACS buffer (1x PBS supplemented with 1% FBS (#F9665, Sigma-Aldrich), 10 μg/mL human-IgG (# I8640-100MG, Sigma-Aldrich), and 0.01% sodium azide) before being transferred into a 96-well V-bottom plate at 100,000 cells/well. Non-specific Fc-receptor interactions were blocked by 30 min incubation in FACS buffer. Cells were spun again and stained with primary antibodies diluted in FACS buffer (Table 5) directly without fixation for 1 h in the dark. If using fluorescent primary antibodies, isotype controls conjugated to the same fluorophore, obtained from the same company, were used at the same final antibody concentration. If using non-fluorescent primary antibodies, isotype control was achieved by staining with secondary antibody only.

Whenever secondary antibody incubation was necessary, cells were washed once with 1xFACS buffer following primary antibody incubation, then stained with secondary antibodies diluted in FACS buffer (Table 5) for 30 min in the dark. After staining, cells were washed once with FACS buffer and fixed by resuspension in 2% PFA (# J61899.AP, ThermoFisher Scientific) in FACS buffer. The protocol steps for total marker staining were identical to those used for surface marker staining, with the following exception: cell fixation was carried out with 2% PFA in FACS buffer for 10 min at RT immediately after transfer to a 96-well V-bottom plate. Cells were then washed once with FACS buffer, and permeabilised with 0.1% Triton-X100 (# T8787, Sigma-Aldrich) in 1x PBS for 10 min at RT, before blocking and staining.

Forward scatter (FSC-H), side-scatter (SSC-A), and fluorescence measurements were obtained by passing single-cell suspension through a Cytoflex LX (Beckman Coulter) flow cytometer. A minimum of 20,000 events were recorded per condition. Laser settings and gating strategies are described in detail in individual results sections. Acquired data was analysed using the FlowJo^TM^ v10 software.

#### Proteomics

See Supp. Methods for details of Sample Preparation, Proteomic Digestion, Phosphopeptide Enrichment, Manual Tip Enrichment for Purified Tau Inputs, LC-MS/MS of total proteome and phosphoproteome samples, EV and conditioned media samples, and Data Analysis.

#### Cryo-electron tomography (CryoET)

EVs were purified as described above from conditioned media harvested from 10 cm dish cultures of iMGL cells, either treated with vehicle or 1 µM rTauF overnight. Extra care was taken to reduce the final sample volume to ∼10 µL to concentrate the EVs on the grids.

2.5 µL of EV sample was applied to glow-discharged lacey-carbon grids (#AGS187-4, Agar scientific) three times using a Vitrobot (Mark IV, Thermo Fisher), using a blotforce of 6 for 3 sec before plunge-freezing in liquid ethane. No fiducials were added to avoid diluting the samples. The grids were then stored in liquid nitrogen until use.

##### Microscope setup

Data was collected using a Thermo Scientific 300 kV Titan Krios electron microscope with a Gatan K3 Bioquantum direct electron detector. The pixel size was1.4 Å and the energy filter slit width 20 eV. Tilt series from specified areas of interest were acquired using a dose-symmetric scheme with a tilt range of ±51° at 3° increments, making the total electron dose 3 e^-^/area^2^ * 35 tilts = 105 e^-^/area^2^. The defocus range was set between – 2.5 µm and –5.5 µm to optimise contrast.

##### Tomogram reconstruction

The fiducial-free tilt series were aligned using patch tracking in Aretomo version 1 from the IMOD software package, with simultaneous iterative reconstructive technique (SIRT) implemented. Tomograms were binned at either 4 or 8, making the reconstructed tomogram pixel sizes 5.6 Å (bin4) or 11.2 Å (bin8), and a lowpass filter of 0.035 (cutoff radius), 0.35 (sigma) was applied via the mtffilter function in IMOD.

##### Tomogram analysis and measurements

Tomograms were viewed and analysed using 3dmod from the IMOD software package. For generating Z-projections, tomograms with binning setting 8 were opened and rotated in the Slicer window and appropriate Z-projections made, typically of ∼ 5 slices, so ∼11 nm in thickness. EV diameters were measured manually using the Drawing Tool “Measure” and panning through each tomogram to find the maximum diameter of each EV within the field of view. Fibrils were drawn as open contours through the 3D volume of EVs while tomograms were open in the slicer window with the image thickness set to 5 to enhance contrast. Measurements were made from EVs containing obvious fibrils and within such an EV, only the fibrils with clear start and endpoints were measured, hence these quantifications do not claim to capture the entire population and probably provides an underestimate in terms of EVs containing fibrils. All fibril measurements were expressed as absolute length (in comparison with length of un-internalised rTauF), as well as matched up with the diameter of the EV from which the fibril originated and then expressed both as a function of EV diameter and as a percentage of EV diameter. Similarly, for fibril crossover distances, in a subset of EVs where this phenomenon was very obvious, the interval between nodes were drawn as open contours and averaged per fibril.

#### Thioflavin T assay

ThT assay was used to verify the presence of β-sheets in tau fibril preparations [69, 70]. 100 µL reactions were prepared in black, clear-bottom 96-well PhenoPlate^TM^ (#6055300, PerkinElmer) by mixing 28 μM tau monomer, 7 μM heparin, and 25 μM ThT (# T3516-5G, Sigma) in 1x PBS, pH 7.2. The sealed plate was incubated at 37°C with 250 rpm agitation for 11 days. ThT fluorescence readings (excitation wavelength 440 nm / emission wavelength 510 nm) were taken at regular intervals using a Spectramax M5e Multi-mode Microplate reader (Molecular Devices). Background fluorescence of 1x PBS buffer was subtracted from all values at corresponding time points.

#### 4R tau Real-Time Quaking-Induced Conversion (RT-QuIC) assay

A modified version of the 4R tau RT-QuIC seed amplification assay[75, 76] was used to determine the seeding capacity of soluble and insoluble intracellular tau and tau secreted to conditioned medium by iMGL.

Cells were differentiated in 6-well tissue culture plates and treated overnight with 1 μg/mL of rTauM, rTauF, hTau or vehicle. To remove uninternalised tau cells were incubated with 2.5% TryplE^TM^ for 1 min[67] followed by 1x PBS wash. Cells were left to process internalised tau in standard, tau-free, iMGL differentiation medium at 37°C, 5% CO2. After 24 h, conditioned medium and cell lysates were collected. The conditioned media was spun at 400 g for 5 min at 4°C to remove floating cells and at 2000 g for 20 min to remove cell debris, before being frozen on dry-ice and stored at -80°C until further processing. Cells were lysed with 100 μL of ice-cold Triton lysis buffer (1% Triton-X100 in 50 mM Tris (# T1503-500G, Sigma-Aldrich), 150 mM NaCl, pH 7.6) supplemented with EASYpack Protease Inhibitor Cocktail (#5892791001, Roche) and Pierce^TM^ Phosphatase Inhibitor Mini Tablets (#A32957, Thermo Fisher). Cell lysates were collected by scraping, then centrifuged at 21,000 g for 30 min at 4°C. Supernatants containing total cell protein including soluble tau (the Triton fraction) were transferred to Eppendorfs. Pellets containing insoluble tau were solubilised in SDS lysis buffer (1% SDS in 50 mM Tris, 150 mM NaCl, pH 7.6) supplemented with EASYpack Protease Inhibitor Cocktail and Pierce^TM^ Phosphatase Inhibitor Mini Tablets to form the SDS fraction[67]. The protein concentration of both fractions was quantified with Pierce^TM^ BCA Protein Assay Kit (#23227, ThermoFisher Scientific) and normalised to equal amounts by dilution with ddH2O. Samples were snap-frozen in liquid N2, then stored at -80°C. The amount of tau in supernatants, triton and SDS-fractions were quantified with Invitrogen^TM^ Tau (Total) ELISA as described above.

Medium and cell lysate aliquots were shipped on dry ice for RT-QuIC analysis by Alessia Santambrogio at the Yusuf Hamied Department of Chemistry, University of Cambridge. K11 tau (tau residues 244-394) was used as tau seeding substrate. One aliquot of lyophilized K11 purified from E. coli was dissolved in 1 mL of 8 M GuHCl (#G3272, Merck) prior to size-exclusion chromatography (SEC) separation on a Superdex 75 10/300 GL column (#17517401, Cytiva) equilibrated in 20 mM sodium phosphate, 200 mM NaCl, pH 7.4. 50 μL/well total volume reactions were prepared in a 384-well optical BTM Polybase Black plate (#242764, Thermo Scientific) by adding the individual samples (i.e. the Triton/SDS cell lysate fractions and the conditioned medium) in respective 1:10,000 and 1:100 final dilutions to the reaction buffer. The reaction buffer was comprised of 3 μM K11 tau monomer, 10 μM ThT, 500 mM Na2SO4, and 10 mM HEPES buffer at pH 7.4. Reactions were transferred to a 384-well Nunc microplate (#242764, non-treated polymer base) covered with aluminium sealing cover to prevent evaporation and subjected to rounds of 60 sec shaking (500 rpm, orbital) and 60 sec rest with periodic ThT readings every 15 min at 37 °C in a FLUOstar Omega lite microplate reader (BMG labtech).

#### Tau seeding assay in FRET Biosensor cell line

Tau RD P301S FRET Biosensor cell line (ATCC, CRL-3275) was used to evaluate the seeding competency of human recombinant 2N4R tau fibrils and the presence of seed-competent tau species in iMGL derived samples[47]. This biosensor consists of HEK293T cells that stably express tau RD harbouring the P301S mutation fused to either CFP or YFP. Internalisation of tau seeds induces the aggregation of the tau reporter proteins, producing a FRET signal that can be measured by flow cytometry. Cells were plated in 96-well plates at 25,000 cells per well. The following day, transfection complexes were prepared by combining the transfection reagent mix (1 µL Lipofectamine2000 + 9 µL OptiMEM) with 10 µL of CM or protein mix (50 nM rTauF, 1 µg of Triton/SDS lysates or 2 µg of EVs diluted in OptiMEM). Liposomes were incubated 20 min at RT before being added to the cells. At 48 h post-transfection, cells were lifted with StemPro Accutase and fixed with 2% PFA for 10 min at RT. BD Fortessa X20 was used to perform FRET flow cytometry. To measure CFP and FRET signals, cells were excited with a 405 nm laser, and fluorescence was detected with 405/50 nm and 525/50 nm band-pass filters, respectively. The FRET-positive gate was adjusted to cells that were transfected with lipofectamine alone. Each sample was tested in triplicates and 10,000 events per replicate were analysed using FlowJo^TM^ v10 software. Data is represented as integrated FRET density (percentage of FRET-positive cells multiplied by the median fluorescence intensity of FRET-positive population). Integrated FRET density from iMGL samples were normalised to total tau levels in each analysed fraction, which was quantified by ELISA. To visualize intracellular tau inclusions in the FRET-positive population, cells were imaged using Amnis ImageStreamX MkII and IDEAS 6.2 software.

#### Tau seeding assay in iNeurons

Human iPSCs were differentiated into cortical neurons by the forced expression of the neuronal transcription factor neurogenin 2 (Ngn2), as previously described[77], with slight modifications. iPSCs (125,000 cells/cm^2^) co-expressing the reverse tetracycline-controlled transactivator (rtTA) (pUbiq-rtTA, 19780, Addgene) and Ngn2 under the TetO promoter (pTetO-Ngn2-T2A-Puro, 52047, Addgene) constructs were plated in 96 well-plates precoated with 0.01% poly-L-ornithine and 10 µg/mL laminin. Cells were grown in Neurobasal plus medium (A3582901, Thermo Fisher) supplemented with 10 ng/mL human BDNF (450-02-10, Peprotech), 0.2 µM L-ascorbic acid (A0278, Sigma-Aldrich-Aldrich), 10 ng/mL human NT-3 (450-03-10, Peprotech), 3.3 µg/mL geltrex (A1413302, Thermo Fisher), GlutaMAX^TM^ (35050061, Thermo Fisher) and B-27 plus (A35828-01, Thermo Fisher) (Supp Table 3). Doxycycline (4 µg/mL, D9891, Sigma-Aldrich-Aldrich) was added on DIV0 to induce TetO-dependent Ngn2 expression and kept in the cultured until the end of the experiment. The media was also supplemented with 10 μM ROCK inhibitor (Y-27632, #1201029, Abcam) and kept in culture until DIV4. After 24 h puromycin (A1113803, Thermo Fisher) selection was performed on DIV2 and DIV3 at 2 µg/mL and 3 µg/mL, respectively.

Adeno-associated viral (AAV) particles expressing human 0N4R P301S Tau fused to Venus (AAV2.1/2.2-hSyn-TauP301S-Venus) were generated by the McEwan lab. Briefly, chimeric AAV1/2 capsid particles encoding P301S tau-venus under the hSyn promoter were generated by transfection of AAVPro HEK293T cells (632273, Takara) followed by two rounds of iodixanol (D1556, Sigma-Aldrich) gradient ultracentrifugation as previously described [78]. Viral titres and purity were assessed using qPCR for the AAV ITR and SDS-PAGE respectively. On DIV4, iNeurons were transduced with the AAV particles. On DIV5, a full media change was performed and 1 µM AraC (C1768, Sigma-Aldrich-Aldrich) was added to prevent the growth of undifferentiated cells. 50% media change was done every 3 days.

At DIV11, iNeurons were transfected using Lipofectamine200 with 2 µg of EVs as described above. At 65 h post-transfection, iPSC-neurons were fixed with 100% pre-chilled methanol at -20°C for 15 to remove any soluble tau[9], and immunostained with anti-MAP2 and anti-AT8 as described above. The integrated Venus or AT8 density was calculated as the product of Venus or AT8-positive cells and mean Venus or AT8 intensity, normalised to the number of MAP2-positive cells. The percentage of colocalization was calculated as the ratio of the number of cells with more than 50% overlap of Venus and AT8 signals to the number of MAP2-positive cells, times 100.

## Statistical information

Statistical analyses were performed in GraphPad Prism v9.0 (GraphPad Software Inc.) using one-way ANOVA, two-way ANOVA or Student’s t-test, with Tukey’s, Dunnett’s or Šídák’s multiple comparison tests, as appropriate. Specific methods are detailed in individual figures. Data is presented as mean ± standard error of the mean (SEM), unless stated otherwise. Significance was defined as *P ≤ 0.05, **P ≤ 0.01, ***P ≤ 0.001, ****P ≤ 0.0001. n.s.= not significant.

## Data availability

Raw RNA-seq .fastq, processed kallisto abundance.tsv and abundance.h5 files have been deposited in Gene Expression Omnibus (GSE291195). The mass spectrometry data have been deposited in ProteomeXchange (PRIDE database).

## Code availability

All code for RNAseq is available at https://github.com/S-Washer/Karabova_2025_Tracking_tau_and_cellular_responses_in_microglia_RNAseq/tree/main

## Acknowledgements and Funding

This work was supported by funding from The James Martin 21st Century Research Foundation (Partners in Tau). M.K.K. was supported by an Oxford E.P. Abraham Graduate Scholarship, with additional funding from Alzheimer’s Research UK Thames Valley Network Centre. A.H. was supported by a research grant to S.A.C. from Eli Lilly and Company. Human tissue samples were provided by the London Neurodegenerative Diseases Brain Bank at King’s College London. The brain bank is partly supported through the Brains for Dementia Research programme, jointly funded by Alzheimer’s Research UK and the Alzheimer’s Society.

We thank: Adam Harding for help with protein purification; Abul Tarafder and Tanmay Bharat for SENP2 protein; Amy Napier for lentivirus production; Robert Hedley and Vasiliki Tsioligka for help in the Don Mason Flow Cytometry Facility; Alan Wainman for help in the Dunn School Light Microscopy Facility; Richard Wade-Martins and Sarah Pearce for Opera Phenix access and assistance; Byron Caughey for advice on tau RT- QuIC; Lindsay Baker, Emma Silvester and Ellen Reed for assistance with tomogram reconstruction.

## Author contributions

Conceptualisation, M.K.K., A.D.S-B., A.H., K.S.K., W.S.J., S.A.C. Methodology, M.K.K., A.D.S-B., A.H., Z.B., E.J, C.E.M., D.P.O., I.V., R.F., S.K., K.A.X.C., W.A.M., W.S.J., S.A.C.

Software, S.J.W. Formal analysis, A.H., S.J.W., D.P.O. Investigation, M.K.K., A.D.S-B., A.H., Z.B., D.P.O, I.V., S.S.H., E.J., C.E.M., T.R.S.M-P., R.M., A.S., M.A.M. Data curation,

S.J.W., D.P.O. Writing - original draft, M.K.K., A.D.S-B., A.H., S.J.W., S.A.C. Writing - review and editing, M.K.K., A.D.S-B., A.H., S.J.W., Z.B., E.J, C.E.M., D.P.O., W.A.M.,

K.S.K., W.S.J., S.A.C. Visualisation, M.K.K., A.D.S-B., A.H., S.J.W., D.P.O. Supervision,

M.V., K.S.K. T.A.D., W.S.J., S.A.C. Project administration, S.A.C. Funding Acquisition, T.A.D., K.S.K., W.S.J., S.A.C.

## Materials & Correspondence

Sally A. Cowley, James and Lillian Martin Centre for Stem Cell Research, Sir William Dunn School of Pathology, University of Oxford, Oxford OX1 3RE, UK

