## Supplementary methods and figures for "Tracking tau and cellular responses in human iPSC-microglia from uptake to seedable secretion in extracellular vesicles"

### Supplementary information

#### Supplementary Methods

#### Human recombinant 2N4R tau protein expression and purification

SUMO_TauWT 2N4R_pETM11SUMO3_sense plasmid was generated in-house through subcloning Tau protein sequence from pNIC28-Bsa4_6xHis-tag_TauWT 2N4R vector, a gift from Donatella Di Rienzo, Oxford Drug Discovery Institute, into SUMO-GFP_TB004_pETM11SUMO3_sense plasmid, a gift from Tanmay Bharat, Sir William Dunn School of Pathology (Kreger Karabova, M. (2023). Investigating the role of human microglia in tau pathogenesis [PhD thesis]. University of Oxford https://ora.ox.ac.uk/objects/uuid:a9337b8a-d4bd-4bc1-8284-6b803c5e9f0a). A total of 15 ng of plasmid DNA with confirmed correct sequence were transformed into BL21(DE3) *E. coli* and glycerol stock made. Glycerol stocks were used to inoculate 40 mL LB broth with 50 µg/mL kanamycin. Starter cultures were grown overnight at 37°C with 180 rpm agitation. The overnight culture was further inoculated to 1 L of Terrific Broth (TB) broth supplemented with 50 µg/mL kanamycin and 1x TB salts and returned to the shaking incubator at 37 °C with 225 rpm agitation. At $OD_{600}$ = 0.5-0.7, the cells were induced with 0.5 mM isopropyl b-D-1-thiogalactopyranoside (IPTG) and left to grow for 1 h at 37°C and 225 rpm. Cells were then incubated overnight at 25°C with 200 rpm agitation. Cell pellets were harvested by centrifugation at 5,000 xg for 30 min at 4°C using the Beckman Coulter Avanti JXN-26 centrifuge (JLA-8.1000 rotor). Pellets were resuspended in 25 mL ice-cold lysis buffer per 1 L of pelleted culture and stored at -80°C until required.

On the day of the purification, bacterial cells were thawed, lysed using EmulsiFlex-C5 French Press homogeniser operating at 15 kpsi, and centrifuged at 4,500 xg for 1 h. An additional step of 10 min of direct heating in a 70°C water bath, followed by 4,500 xg centrifugation for 1 h at 4°C, was used for the large-scale purification of SUMO-tagged tau. Pellets were discarded and supernatant passed through a 0.45 μm filter. Samples were kept on ice throughout the entire purification procedure except during the cell heating step.

Purification steps were done using ÄKTA Pure 25M. As all steps were carried out at 4°C, buffer pH was adjusted at 4°C and following sterile-filtering and degassing, were kept at 4°C (see Table 10). The crude extract was applied to HisTrap column pre-equilibrated with 5 column volumes (CV) of buffer A1. The column was then washed with 50 CV buffer A2, which contains 0.1% Triton X-114[5] and it is a critical step for endotoxin removal from the preparation, followed by 20 CV buffer A1. Protein was eluted from the column with a linear gradient of buffer B. Overnight dialysis at 4°C in SnakeSkinT$^{M}$ Dialysis Tubing in the presence of Sentrin-specific protease 2 (SENP2) protease (1:100 protease:protein ratio), against 4 L of dialysis buffer was used to remove imidazole and cleave the 6xHis-SUMO tag.

SENP2 protease, purified in-house from *E. coli,* was a generous gift from Tanmay Bharat lab. The dialysed and cleaved product was re-applied to a HisTrap column pre-equilibrated with 5 CV of buffer C. Tag-less tau was eluted from the column with a linear gradient of buffer D to isolate the tag-less tau. Fractions containing cleaved tau were pooled and further purified using the CaptoS column pre-equilibrated with 5 CV of buffer E. The column was washed with 6 CV of buffer E and eluted with linear gradient of buffer F. Purest tau-containing fractions were concentrated to 2 mL using 10K MWCO Pierce Protein concentrator and filtered through a 0.22 μm cut-off filter. Residual endotoxin was removed from the final product using Pierce High-Capacity Endotoxin Removal Spin Column, as per manufacturer's instructions. Endotoxin levels were quantified with Pierce Limulus Amoebocyte Lysate (LAL) Chromogenic Endotoxin Quantitation Kit. The concentration of endotoxin-free 2N4R tau monomer was measured using Pierce BCA Protein Assay Kit - Reducing Agent Compatible due to the presence of β-mercaptoethanol in the elution buffer. Sample was aliquoted into Protein LoBind® tubes, and stored at -80°C.

If columns were re-used multiple times, a column cleaning procedure (1 h soak with 1 M NaOH followed by 10 CV ddH2O and 10 CV buffer A1 or buffer E washes) would precede sample application. Eluted protein fractions were analysed with SDS-PAGE gel electrophoresis. A total volume of 15 μL eluate was mixed with NuPAGE™ 4x LDS sample buffer and boiled at 70°C for 10 min. Samples were loaded on Novex™ WedgeWell™ 8-16%, Tris-Glycine pre-cast mini gels, and ran in Novex™ Tris-Glycine SDS Running Buffer at 200 V for 45 min. A 1 h RT gel incubation with SimplyBlue™ Safe Stain was used for protein detection.

All resources and buffer compositions are described in Supplementary Tables 7 and 8.

**Enrichment of tau PHFs from human AD brains**

Brain-derived tau (hTau) was enriched from Alzheimer's patient brains and QCed by E.M and Western blot and toxicity assessed (Supp. Fig 2A-C,E). The treatment dose of 0.25 µg/mL was chosen as this conferred no toxicity to iMGL after 24 h incubation.

Samples of brain material were obtained from the London Neurodegenerative Diseases Brain Bank at King's College London and processed at Eli Lilly and Company's facility in Indianapolis, IN, US. Seeding-competent tau was extracted as previously described[6] with modifications noted below from samples of frontal and temporal cortex, obtained from individuals diagnosed with AD (BNE/Braak stage 5-6), with an average age of 80.2 ± 2.3 years (range: 63-98 years). Tissues were dissected after an average postmortem delay of 30.6 ± 4.2 h (range: 10-66 h). Numerous brain tissue pieces from each AD patient tissue slab were preliminarily screened using a proprietary AT8 tau Alphascreen assay developed by Lilly (AT8 and DA9 antibodies obtained from Peter Davies). Each preparation utilized tissue pieces with high, medium, and low AT8 tau signal with a cut-off of > 0.25 ug/mL AT8 tau signal. Percentages of high, medium, and low AT8 tau signal tissue pieces were maintained across preparations to ensure consistency.

A total of 100 g of tissue per donor was processed by repeated homogenisation and filtration as follows: Frozen postmortem tissue samples were shipped on dry ice and stored at -80˚C until needed. Tissues were defrosted on ice and broken into smaller pieces before being added to cold homogenisation buffer (4 mL/1 g of tissue, sterile PBS (#14190-144, Gibco) containing Complete protease inhibitor cocktail (#11836145001, Roche)). Homogenisation was achieved using an Ultra Turrax (IKA T25, 20,000 rpm, 10min), after which the homogenate was centrifuged at 10,000 rpm for 10 min at 4°C. The supernatant was carefully removed and filtered through a 2-ply tissue into a beaker on ice, upon which the pellet was resuspended again in 400 mL homogenisation buffer and the homogenisation and filtration process repeated for a total of three times.

The resulting homogenate (~1000 mL) was then split between 500 mL centrifuge bottles (#431123, Corning) and appropriate volumes of 30% sarkosyl solution (N-Lauroylsarcosine sodium salt 30% aqueous solution, #61747, Sigma-Aldrich-Aldrich) were added for a final concentration of 1% sarkosyl. Following a 1 h incubation at room temperature on a flat rotating shaker at a medium speed, samples were centrifuged for 1 h at 45,000 rpm at 4°C (Optima XPN-80 ultracentrifuge (45Ti rotor), Beckman Coulter 65 mL ultracentrifuge tubes #355655). The extracted pellet was washed once by carefully adding and removing 1 mL homogenisation buffer, before being resuspended in 50 µL/initial gram of sample (5 mL/100 g tissue), of PBS containing 0.5x EDTA-free complete protease inhibitor cocktail (#11873580001, Roche) and transferred to a 15 mL falcon tube for sonication. Each sample was sonicated (Fisherbrand™ Model 120 Sonic Dismembrator with CL-18 probe) as follows until visible clumps were gone: 10 x 1 sec pulses (1 sec on, 1 sec off) at 40% amplitude ~four times, and then again ~twice after combining the samples. For this study, three separate preparations comprising a total of 17 AD patient tissue samples (12 females and 5 males at 63-90 years old) were pooled. The total-tau concentration of the pooled preparation was assessed by a proprietary Alphascreen assay developed by Lilly (DA9 and TG5 antibodies obtained from Peter Davies) in the presence of guanidine or PBS, using monomeric tau and PHF as standard curves and resulted in 78.7 µg/mL.

A rat cortical neuron seeding cell assay was performed to determine the optimal seeding concentration for the pooled tau seed preparation. Briefly, the pooled tau seed preparation was diluted to 2 µg/mL total tau in cell culture medium, sonicated (60 x 1 sec pulses at 20% amplitude), and filtered through a 0.22 µm filter into a deep-well plate. The resulting filtrate was serially diluted two-fold for a nine-point concentration-response curve. Enough media was removed from the rat cortical neuron cells at DIV8 (plated at 30K cells/well at DIV0) to leave 50 µL in the well, and 50 µL of cell medium or serially diluted pooled tau seed was added for a final concentration range of 0.0039-1 µg/mL total tau pooled tau seed. Half of the treated cell medium was removed and replaced with fresh medium on DIV16. On DIV22, all the medium was removed and the cells washed 3x with 100 µL PBS. After the final wash, the PBS was removed and 100 µL of 100% methanol (chilled at -20˚C) was added for 15 min at 4˚C to remove soluble proteins and fix the cells. Cells were washed 3x with 100 µL PBS and a final 100 µL PBS was added for storage at 4˚C prior to immunostaining and imaging.

To determine the amount of tau seeding, PBS was removed from the fixed cells and 100 µL of Odyssey blocking buffer with 0.1% Triton was added and incubated at room temperature for 60 min. The blocking buffer was removed and 100 µL of primary antibodies diluted in Odyssey blocking buffer with 0.1% Triton was added and incubated overnight at 4˚C while shaking at a slow speed. Cells were stained for rodent total tau using the T49 tau antibody at a 1:2000 dilution and co-stained with a NeuN antibody at a 1:1000 dilution to detect neurons. Primary antibodies were removed, and cells were washed 3x with 100 µL PBS, followed by incubation with 100 µL of appropriate secondary antibodies and nuclei-stain Hoechst 33342 at a 1:1000 dilution in Odyssey blocking buffer with 0.1% Triton for 1 h at room temperature. Secondary antibodies were removed and cells were washed 3x with 100 µL PBS and a final 100 µL PBS was added for storage in the dark at 4˚C until imaged using an Operetta instrument and data analysis software to determine the number of T49 tau inclusions per number of nuclei at each seed concentration. The resulting average EC80 seed concentration (0.1 µg/mL) determined from two separate tests was used as the optimal seeding concentration for additional seeding studies.

### Proteomics

#### *Sample Preparation*

iMGL were lysed in RIPA buffer supplemented with 5% SDS and protease and phosphatase inhibitors (Roche). Proteins secreted to the conditioned medium (CM) were precipitated by overnight incubation with 6% trichloroacetic acid (TCA) at 4°C in agitation. CM was then centrifuged at 10,000 xg for 10 min at 4°C, the supernatant was discarded, and the pellet was air-dried for 10 min and lysed in RIPA with 5% SDS. EVs were isolated as described above and lysed on ice for 10 min with 10x RIPA supplemented with 5% SDS and protease and phosphatase inhibitors.

### *Proteomic Digestion*

Approximately 300 µg of protein material was digested using an S-Trap™ sample processing procedure in a 96 well format according to the manufacturer's instruction[7]. Briefly, proteins were reduced with 20 mM DTT for 30 min at RT and alkylated with 40 mM of iodoacetamide, also for 30 min at RT and in the dark. Proteins were acidified to reach a final concentration of 1.2% phosphoric acid. Samples were then subjected to a six-fold dilution with 90% methanol/100 mM tetraethylammonium bromide (TEAB). Diluted protein material was then loaded and washed on an S-Trap™ plate. Each protein sample was digested with 6 µg of trypsin overnight at 37°C, achieving a final protein:enzyme ratio of 50:1. Tryptic peptides underwent sequential elution from the column, with a final elution buffer of 50% acetonitrile (ACN). EV and conditioned media samples were subjected to an identical procedure, except that in the starting volume of 50 µL, the concentration/amount of material was unknown.

### *Phosphopeptide Enrichment - Bravo Enrichment*

Approximately 10% of each tryptic peptide digest was kept aside to perform a total proteome analysis. The remaining sample was desalted using SOLA HRP cartridges and adapted with minor changes to manufacturer's protocol for phosphopeptide enrichment. Briefly, cartridges were first conditioned with solvent B (70% ACN, 0.1% trifluoroacetic acid (TFA)) and then washed with solvent A (0.1% TFA in H2O). Peptides were loaded in a 1:1 ratio (sample/solvent A). Bound peptides were then washed with solvent A, followed by elution with 1M glycolic acid in 50% ACN, 5% TFA. Phosphopeptide enrichment was performed using the pre-defined Agilent Bravo AssayMap liquid handler workflow for TiO2 cartridges (AssayMap 5 µL Titanium Dioxide TiO2 cartridges, Catalogue Number G5496-60016, Agilent). The prime and syringe wash solution consisted of 50% ACN, 5% NH3 in H2O and the equilibration and cartridge wash solution was 50% ACN, 2% TFA, 1 M glycolic acid in H20. Phosphopeptides were eluted in 15% ACN, 5% NH3. Samples were loaded onto the cartridges at 2.5 µL/min. Eluted phospho-enriched peptides were dried down and resuspended in 5% DMSO and 5% formic acid (FA).

### *Manual Tip Enrichment for Purified Tau Inputs*

Briefly, 250 µg of dried samples were re-suspended in 50 µL loading buffer (80% ACN, 5% TFA and 1M glycolic acid). TiO2 columns (Glycogen) were first spun down for 30 sec at 2,000 x g with the caps on to ensure all material was at the bottom of the tip. Columns were then washed once with 50 µL of elution buffer (5% ammonium hydroxide) at 1,500 x g for 2 min, followed by three washes of 65 µL loading buffer at 2,000 x g for 5 min. Following this, the LoBind® Eppendorf tubes were replaced with fresh ones. Samples were then loaded in 20 µL volumes and at 2,000 x g for 5 min until all material was bound to the column. Columns were washed twice with 65 µL loading buffer, once with 65 µL wash buffer 1 (80% ACN, 0.2% TFA), and then finally, two washes of 65 µL wash buffer 2 (20% ACN). All spins were performed at 2,000 x g for 5 min. After changing to fresh LoBind® Eppendorf tubes, 20 µL of 20% FA was added to each tube. Phosphopeptides were eluted by two washes of 20 µL elution buffer into the same tube. To recover residual eluate, a spin at 2,000 x g for 2 min was performed.

#### *LC-MS/MS of total proteome and phosphoproteome samples*

Peptides were chromatographically separated on a 15 cm x 150 µm x 1.5 µm analytical column (EV1137, Evosep) using the 30 samples per day (spd) gradient method of an Evosep One LC system. Eluted peptides were directed for mass spectrometry (MS) analysis on a timsTOF Pro mass spectrometer (Bruker). Phosphopeptide MS data were acquired in ddaPASEF mode. The ion mobility window was 0.60-1.60 Vs/cm2, with an accumulation and ramp time of 100 ms. The mass range of MS and MS/MS scans was *m/z* 100 – 1,700. MS/MS spectra were acquired in 10 PASEF ramps with a 4-frame overlap, giving a cycle time of 1.17 sec. Ions were selected for PASEF MS/MS if they met an intensity threshold of 2,500 and were sampled multiple times until a target intensity of 20,000 was reached. A polygon filter was used to exclude singly charged ions from MS/MS selection. Ions selected for fragmentation were isolated by the quadrupole and fragmented using an ion mobility-dependent collision energy that increased nonlinearly over the ion mobility range as follows: a collision energy (eV) of 20 at a 1/K0 of 0.60 Vs/cm2, 22 at 0.70, 25 at 0.75, 30 at 0.80, 35 at 0.85, 40 at 0.90, 45 at 0.95, 50 at 1.00, 55 at 1.10, 60 at 1.20, 65 at 1.30, 70 at 1.40, 75 at 1.50, and 80 at 1.60. A dynamic exclusion time of 24 sec was used. Total proteome MS data were acquired in diaPASEF mode using 16 diaPASEF scans per TIMS-MS scan with an accumulation and ramp time of 100 msec, for a total cycle time of 1.80 sec. The ion mobility range was set to 0.6-1.6 Vs/cm2. Each mass window isolated was *m/z* 26 wide, ranging from *m/z* 400-1200 with an ion mobility-dependent collision energy that increased linearly from 20 eV to 59 eV between 0.6-1.6 Vs/cm2.

#### *LC-MS/MS of EV and conditioned media samples*

Samples were analysed by LC-MS/MS using the Vanquish Neo UHPLC connected to Thermo Orbitrap Ascend mass spectrometer equipped with a high-field asymmetric ion mobility spectrometry (FAIMS) Pro Duo interface (all Thermo Fisher Scientific). The Vanquish Neo was operated in “Trap and Elute” mode using a PepMap Neo trap (5 µm, 300 µm x 5 mm; Thermo Fisher) and EASY-SPRAY PepMapNeo column (50 cm x 75 µm, 1500 bar; Thermo Fisher). Tryptic peptides were trapped and separated over a 75 min gradient, going from 2 to 18% B (0.1% FA in 100% ACN) in 40 min, to 35% in 20 min, up to 99% B in 1 min and then staying at 99% for 14 min. The flow rate was maintained at 300 nL/min throughout the gradient. The FAIMS Pro™ interface was operated in standard resolution with carrier gas flow rate of 3.8 L/min and compensation voltage set to -45. MS data were acquired in Data Independent Acquisition (DIA) mode, with minor changes from our previously described method[8-10]. Briefly, MS1 scans were collected in the Orbitrap mass analyser at a resolving power of 45K at *m/z* 200 over a range of *m/z* 350 – 1650. The MS1 normalised AGC was set at 125% (5e5 ions) with a maximum injection time of 91 msec and a RF lens at 30%. DIA MS2 scans were then acquired using the tMSn scan function at an Orbitrap resolution of 30K over 40 scan windows of variable width, with a normalised AGC target of 1000%, maximum injection time set to auto and a 30 % collision energy.

#### *Data Analysis*

MS raw files were searched against the Homo sapiens database downloaded from UniProtKB on November 7th 2023. Total proteome and phosphoproteome Data Dependant Acquisition (DDA) data were searched using Fragpipe (v22.0), while EV and conditioned media DIA data was searched using DIA-NN Software (v1.9) in library-free mode. Carbamidomethyl of Cys residues was set as a fixed modification, while oxidation of Met and N-term acetylation were set as variable modifications. Phosphorylation of Ser, Thr, and Tyr was set as a variable modification, where relevant. A maximum of 1 missed cleavage was allowed and the matching between runs function was turned off. Search results were analysed and visualised in Perseus (v. 2.0.10.0) by means of Principal Component Analysis (PCA), Volcano and Scatter plots. Dysregulated proteins and phosphopeptides across conditions were subjected to Gene Ontology and pathway enrichment analysis in STRINGdb (https://string-db.org).

### Supplementary Tables

**Supplementary Table 1: Cell lines used and donor information**

| ID # | Cell line | Donor age | Donor sex | Diagnosis | LRRK2 genotype | Symbol |
|---|---|---|---|---|---|---|
| 1 | SFC840-03-03 | 65-69 | F | Healthy control | WT/WT | circle |
| 2 | SFC841-03-01 | 35-39 | M | Healthy control | WT/WT | square |
| 3 | SFC856-03-04 | 75-79 | F | Healthy control | WT/WT | triangle |
| 4 | KOLF2.1S | 55-59 | M | Healthy control | WT/WT | diamond |
| 5 | SFC840-03-03 D10 | 65-69 | F | CRISPR/Cas9 edited | -/- | half black circle |
| 6 | SFC832-03-06 | 75-79 | F | G2019S PD | G2019S/WT | dotted square |
| 7 | SFC833-03-05 | 80-84 | M | G2019S PD | G2019S/WT | dotted diamond |
| 8 | SFC855-03-06 | 55-59 | M | G2019S PD | G2019S/WT | dotted hexagon |

**Supplementary Table 2: Overview of cell lines used in figures**

| Main figures | Panel | Cell line ID | Cell type |
|---|---|---|---|
| Fig. 1 | B | 1 | iMGL |
| | C | 1 | iMac |
| | D | 1 | iMac, iMGL |

| | | | |
|---|---|---|---|
| | E | 1 | iMac |
| | F | 1 | iMac, iMGL |
| Fig. 2 | A, B | 1, 2, 3, 4 | iMac, iMGL |
| | C | 1, 3, 4 | iMac |
| | D | 1, 3 | iMGL |
| Fig. 3 | B-F | 3 | iMGL |
| Fig. 4 | A-E | 3 | iMGL |
| Fig. 5 | B | 1, 2, 3 | iMGL |
| | C | 3 | iMGL |
| Fig. 6 | B-D | 1, 3, 4 | iMGL |
| Fig. 7 | B | 3 | iMGL |
| | C, D | 1, 3, 4 | iMGL |
| Fig. 8 | A | 3, 4 | iMGL |
| | B | 3, 4 | iMGL |
| | D-G | 3 | iMGL |
| Fig. 9 | B | 1, 3, 4 | iMGL |
| | C | 3 | iMGL |
| | D, E | 1, 3 | iMGL, iNeurons |
| **Supplementary figures** | **Panel** | **Cell line ID** | **Cell type** |
| Supp. Fig.2 | D | 1, 3, 4 | iMac, iMGL |
| | E | 4 | iMGL |
| Supp. Fig. 3 | A | 1, 2, 3 | iMac |
| | B | 1 | iMGL |
| | C | 1, 2, 3 | iMGL |
| | D | 1, 3 | iMGL |
| Supp. Fig. 4 | A | 1, 3, 5, 6, 7, 8 | iMac |
| | B | 1, 3, 4, 5, 6, 7, 8 | iMac |

| | | | |
|---|---|---|---|
| | C | 1, 4, 5, 6, 8 | iMac |
| Supp. Fig. 5 | B | 1, 2, 3 | iMGL |
| | C, D | 3 | iMGL |
| Supp. Fig. 6 | D | 3 | iMGL |
| Supp. Fig. 7 | B | 1, 2, 3, 4 | iMGL |
| | C | 2 | iMGL |
| Supp. Fig. 8 | A-D | 1, 3, 4 | iMGL |
| Supp. Fig. 9 | B | 3 | iMGL |
| | C | 1 | iMGL |
| | D | 1, 3 | iMGL |
| Supp. Fig. 10 | C | 4 | iMGL |
| | D | 1, 3, 4 | iMGL |

**Supplementary Table 3: iMac, iMGL and iNeuron differentiation media**

| Medium | Reagent | Supplier | Catalogue number | Final concentration |
|---|---|---|---|---|
| OXE8 | Advanced DMEM/F12 | Thermo Fisher | 12634010 | 1x |
| | GlutaMAX | Thermo Fisher | 35050-038 | 1x |
| | Heparin solution 0.2% | StemCell Technologies | 07980 | 100 ng/mL |
| | Ascorbic Acid | Sigma-Aldrich | A8960 | 0.22 mM |
| | HEPES | Thermo Fisher | 15630080 | 15 mM |
| | FGF-2 | R&D Systems | 4114-TC-01M | 100 ng/mL |
| | TFG-β | Peprotech | AF-100-21C | 2 ng/mL |
| Embryoid body (EB) medium | OXE8 | - | - | 1x |
| | BMP4 | Peprotech | PHC9534 | 50 ng/mL |
| | VEGF | Peprotech | PHC9394 | 50 ng/mL |
| | SCF | Miltenyi Biotec | 130-096-695 | 20 ng/mL |
| XVIVO precursor medium | X-VIVO 15 | Lonza | BE02-060F | 1x |
| | GlutaMAX | Thermo Fisher | 35050-038 | 1x |
| | 2-mercapto-ethanol | Gibco | 31350-010 | 1% |
| | IL-3 | Invitrogen | PHC0033 | 25 ng/mL |
| | M-CSF | Invitrogen | PHC9501 | 50 ng/mL |
| | Pen/Strep | Thermo Fisher | 15140-122 | 1% |

| | | | | |
|---|---|---|---|---|
| OXM macrophage medium | Advanced DMEM/F12 | Thermo Fisher | 12634010 | 1x |
| | GlutaMAX | Thermo Fisher | 35050-038 | 1x |
| | Human recombinant insulin solution | Sigma-Aldrich | 19278-5ML | 5 µg/mL |
| | HEPES 1 M pH 7.4 | Thermo Fisher | 15630080 | 15 mM |
| | M-CSF | Invitrogen | PHC9501 | 50 ng/mL |
| ITMG microglia medium | Advanced DMEM/F12 | Thermo Fisher | 12634010 | 1x |
| | GlutaMAX | Thermo Fisher | 35050-038 | 1x |
| | M-CSF | Invitrogen | PHC9501 | 25 ng/mL |
| | GM-CSF | Invitrogen | PHC2013 | 10 ng/mL |
| | IL-34 | Peprotech | 200-34 | 100 ng/mL |
| | TGFβ1 | Peprotech | 100-21C | 50 ng/mL |
| iNeuron medium | Neurobasal Plus | Thermo Fisher | A3582901 | 1x |
| | GlutaMAX | Thermo Fisher | 35050-038 | 1x |
| | BDNF | Peprotech | 450-02-10 | 10 ng/mL |
| | B27 Plus | Thermo Fisher | A35828-01 | 1x |
| | L-Ascorbic Acid | Sigma-Aldrich | A0278 | 0.2 µM |
| | NT3 | Peprotech | 450-03-10 | 10 ng/mL |
| | Geltrex | Invitrogen | A1413302 | 3.3 µL/mL |
| | Doxycycline | Sigma-Aldrich | D9891 | 2 µg/mL |

**Supplementary Table 4: iMac and iMGL pharmacological and LentiCRISPR treatments**

| **Reagent** | **Supplier** | **Catalogue Number** | **Final concentration** |
|---|---|---|---|
| IFNγ | Gibco | PHC0044 | 100 ng/mL |
| LPS | Invivogen | Tlrl-eklps | 100 ng/mL |
| sRAP | Ximbio | 153996 | 5-500 nM |
| Heparin | Sigma-Aldrich | H3393 | 10 µg/mL |
| FKN | PeproTech | 300-31 | 5 ng/mL |
| Leupeptin | Enzo Life Sciences | ALX-260-009-M005 | 50 µM |
| Pepstatin A | Enzo Life Sciences | ALX-260-085-M005 | 50 µM |
| E64d | Abcam | ab144048 | 50 µM |
| MG-132 | Sigma-Aldrich | 474791 | 25 µM |
| **Target** | **gRNA #** | **Sequence** | |
| LRP1 | 1 | GTCTCGATGCGGTCGTAGA | |
| LRP1 | 2 | TCTGTACTGGACGGACGAT | |

| | | |
|---|---|---|
| LRP1 | 3 | GCCGAGACCGCTCAATACG |
| INTG | 1 | TTGGGCAGAATGTCTGCCC |
| INTG | 2 | AAGCTCCTCACCATGCCCA |
| INTG | 3 | CAGTTGCCTAACAGGAGCA |
| INTG | 4 | ACCAAGGGTTACCAAGAAG |
| INTG | 5 | CACCTCCCAGTGTCTTGAA |
| INTG | 6 | CAGAGGTCAACCTTGACCC |
| INTG | 7 | CTTATTAGGGATCAAGGGT |
| INTG | 8 | CTAGATCTAGGGTGTGTTG |

**Supplementary Table 5: List of antibodies**

| Antibody | Supplier | Catalogue Number | FACS | ICC | WB | ID |
|---|---|---|---|---|---|---|
| LRP1 | Abcam | ab92544 | 1:100 | 1:200 | | |
| LRP1 | Cell Signaling | 64099 | | | 1:500 | |
| Tau12 | Sigma-Aldrich | MAB2241 | | 1:200 | 1:250 | |
| MAP2 | Cell Signalling | 8707 | | 1:200 | | |
| AT8 | Thermo Fisher | MN1020 | | 1:500 | | |
| DA9 | Peter Davies | N/A | | | 1:1000 | |
| T49 | Millipore | MABN827 | | 1:2000 | | |
| PHF-1 | Peter Davies | N/A | | | 1:1000 | |
| Vinculin | Bio-Rad | MCA465GA | | | 1:2000 | |
| Flotilin-I | Cell Signaling | 18634 | | | 1:500 | |
| Annexin-V | Cell Signaling | 8555 | | | 1:500 | |
| CD9 | Cell Signaling | 13174 | | | 1:500 | |
| CD9 | Abcam | ab263019 | | | 1:500 | |
| Calnexin | Abcam | ab133615 | | | 1:2000 | |
| NeuN | Millipore | ABN91 | | 1:1000 | | |
| HT7 | Invitrogen | MN1000 | | | | 50 µg |
| Mouse IgG1 | Santa Cruz | sc-3877 | | | | 50 µg |
| Donkey anti-Mouse A647 | Invitrogen | A31571 | | 1:500 | | |
| Donkey anti-Rabbit A568 | Invitrogen | A10042 | | 1:500 | | |
| Goat anti-Mouse IgG1 647 | Invitrogen | A21240 | | 1:500 | | |
| Goat anti-Chicken 488 | Invitrogen | A32931TR | | 1:500 | | |
| IRDye®800CW Goat anti-Rabbit | LI-COR | 926-32211 | | | 1:2000 | |

| | | | | | |
|---|---|---|---|---|---|
| IRDye®680RD Goat anti-Mouse | LI-COR | 926-68072 | | | 1:2000 |

**Supplementary Table 6: List of reagents**

| Reagent | Supplier | Catalogue Number |
|---|---|---|
| mTESR1 | StemCell Technologies | 85850 |
| Geltrex | Invitrogen | A1413302 |
| EDTA | Invitrogen | 15575-020 |
| TrypLE | Gibco | 12604013 |
| Aggrewell 800 plates | StemCell Technologies | 34815 |
| ROCK inhibitor Y-27632 | Abcam | 1201029 |
| Poly-L-Ornithine | Sigma-Aldrich | P4957 |
| Laminin | Sigma-Aldrich | L2020 |
| Puromycin | Thermo Fisher | A1113803 |
| AraC | Sigma-Aldrich | C1768 |
| DMEM | Sigma-Aldrich | D6429 |
| FBS | Sigma-Aldrich | F9665 |
| Human IgG | Sigma-Aldrich | I8640-100MG |
| Sodium Azide | Sigma-Aldrich | 08591 |
| OptiMEM | Gibco | 31985-062 |
| Lipofectamine2000 | Invitrogen | 11668027 |
| StemPro Accutase | Gibco | A1110501 |
| PBS | Gibco | 14190-144 |
| Complete protease inhibitor cocktail | Roche | 11873580001 |
| 30% sarkosyl solution | Sigma-Aldrich | 61747 |
| Dynabeads® Antibody Coupling kit | Life Technologies | 14311D |
| 4% PFA | Thermo Fisher | J61899.AP |
| Triton-X100 | Sigma-Aldrich | X100 |
| Donkey serum | Bio-Rad | C06SB |
| DAPI | Abcam | ab228954 |
| Hoechst 33342 | Invitrogen | H3570 |
| HEPES | Sigma-Aldrich | H0887 |
| 300-mesh carbon-coated copper grids | TAAB Laboratories | C267 |
| 3nm lacey carbon grids | Agar Scientific | AGS178-4 |
| 200-mesh carbon-coated copper grids | TAAB Laboratories | C101 |
| Uranyl acetate | Agar Scientific | R1260A |
| Glutaraldehyde | Agar Scientific | R1020 |
| PIPES buffer | Sigma-Aldrich | P6757 |

| | | |
|---|---|---|
| Glycine | Sigma-Aldrich | G7126 |
| Osmium tetroxide | TAAB Laboratories | O001/1 |
| Potassium ferrocyanide | Acros Organic | 223111000 |
| Agar 100-Hard epoxy resin | Agar Scientific | AGR1140 |
| BEEM capsules | Agar Scientific | AGG360-1 |
| Hexamethyldisilazane | Sigma-Aldrich | 440191 |
| RIPA buffer | Thermo Fisher | 89901 |
| Complete protease inhibitor cocktail | Roche | 5892791001 |
| Pierce Phosphatase inhibitor mini tablets | Thermo Fisher | A32957 |
| 10x RIPA buffer | Cell Signaling | 9806 |
| Pierce™ BCA Protein Assay Kit | Thermo Fisher | 23227 |
| NuPAGE™ 4x LDS sample buffer | Invitrogen | NP0007 |
| NuPAGE™ 10x sample reducing agent | Invitrogen | NP0009 |
| DTT | Sigma-Aldrich | 43816 |
| Novex™ WedgeWellTM 8-16%, Tris-Glycine pre-cast mini gels | Invitrogen | XP08165BOX |
| Precision Plus Protein Dual Color Standards | Bio-Rad | 161-0374 |
| NovexTM Tris-Glycine SDS Running Buffer | Invitrogen | LC2675 |
| Low-Fluorescence PVDF transfer membrane | Thermo Fisher | 22860 |
| Trans-Blot Turbo Transfer Buffer | Bio-Rad | 10026938 |
| iBind™ Flex blocking solution | Thermo Fisher | SLF1020 |
| Tween® 20 | Sigma-Aldrich | P7949 |
| SDS | Sigma-Aldrich | 75746 |
| Invitrogen™ Tau (Total) Human ELISA kit | Thermo Fisher | KHB0041 |
| IL-1β ELISA kit | Thermo Fisher | 88-7261-88 |
| Human Uncoated IL-6 ELISA kit | Thermo Fisher | 88-7066-88 |
| 3.5 mL polycarbonate tubes | Beckman Coulter | 349622 |
| 13.2 mL ultra-clear polypropylene tubes | Beckman Coulter | 344059 |
| Resazurin | Sigma-Aldrich | 199303 |
| LDH detection kit | Biolegend | 426401 |
| ThT | Sigma-Aldrich | T3516 |
| GuHCl | Merck | G3272 |
| Superdex 75 10/300 column | Cytiva | 17517401 |
| Silver stain kit for Mass Spectrometry | Thermo Fisher | 24600 |
| Ponceau S stain | Sigma-Aldrich | P3504 |
| Trizma® base | Sigma-Aldrich | T1503 |

**Supplementary Table 7: List of reagents for recombinant tau expression and purification**

| Reagent or resource | Supplier | Catalogue number |
|---|---|---|
| **PLASMIDS** | | |
| pNIC28-Bsa4_6xHis-tag_TauWT 2N4R | Donatella Di Rienzo | |
| SUMO-GFP_TB004_pETM11SUMO3 | Tanmay Bharat | |
| pSUMO_TauWT 2N4R_pETM11SUMO3 | In house | |
| **BACTERIAL STRAIN** | | |
| BL21(DE3) *E. coli* | Thermo Fisher | EC0114 |
| **ENZYMES** | | |
| SUMO-Specific Protease 2 (SENP2) | Tanmay Bharat | |
| SUMO-Specific Protease 2 (SENP2) | Leadgene | LDG0015RG |
| **CHEMICALS/REAGENTS** | | |
| LB broth | | |
| Kanamycin | Sigma | K4000 |
| Potassium phosphate monobasic | Sigma | P5379-500G |
| Potassium phosphate dibasic | Sigma | P8281-500G |
| Glycerol | Sigma | G6279 |
| Sodium chloride | Sigma | S7653-1KG |
| Yeast extract | Sigma | Y1625-1KG |
| Tryptone | Sigma | T7293-1KG |
| IPTG | Sigma | I5502-5G |
| HEPES | Sigma | H3375-500G |
| Triton X-114 | Sigma | X114 |
| Imidazole | Millipore | 288-32-4 |
| β-mercaptoethanol | MP Biomedicals | 190242 |
| SimplyBlue$^{TM}$ SafeStain | Thermo Fisher | 46-5044 |
| Bond-breaker$^{TM}$ TCEP Solution | Thermo Fisher | 77720 |
| DTT | Sigma Aldrich | 43816 |
| DNase | Invitrogen | 18047-019 |
| Heparin | Sigma Aldrich | H3393 |
| WF1 for cell culture | Gibco | A12873-01 |
| Pierce Protease Inhibitor XL Capsules, EDTA-free | Thermo Fisher | A37989 |
| **OTHERS** | | |
| Novex WedgeWell 8-16% Tris-Glycine Gel 1mmx15 wells | Invitrogen | XP08165BOX |
| NuPAGE$^{TM}$ 4x LDS sample buffer | Invitrogen | NP0007 |

| | | |
|---|---|---|
| Precision Plus Protein Dual Colour Standards | Bio-Rad | 1610374 |
| Tris-Glycine SDS Running buffer | Thermo Fisher | LC2675 |
| 0.45 µm filters | Starlab | E4780-1456 |
| 0.22 µm filters | Starlab | E4780-1226 |
| HisTrap™ HP | Cytiva | 17524802 |
| HiTrap™ Capto™S | Cytiva | 17544123 |
| SnakeSkin™ Dyalisis Tubing | Thermo Fisher | 68035 |
| Pierce Protein Concentrator PES, 10K MWCO, 5-20 ml | Thermo Fisher | 88528 |
| Pierce High Capacity Endotoxin Removal Spin Column, 0.5 ml | Thermo Fisher | 88274 |
| Pierce LAL Chromogenic Endotoxin Quant kit | Thermo Fisher | A39553 |
| Pierce BCA Protein Assay Kit - Reducing Agent Compatible | Thermo Fisher | 23250 |
| **PLASTICWARE** | | |
| Cuvettes | VWR | 634-0676 |
| 1 L Polypropylene Bottle Assembly | Beckman Coulter | C31597 |
| 75mm bottle top filter - 500 ml | Nalgene filtration products | 291-4520 |
| 15 mL Protein LoBind Tube | Eppendorf | 0030122208 |
| 50 mL Protein LoBind Tube | Eppendorf | 0030122240 |
| 0.5 mL Protein LoBind Tube | Eppendorf | 022431064 |
| 1.5 mL Protein LoBind Tube | Eppendorf | 022431081 |
| **EQUIPMENT** | | |
| Innova®44 Incubator Shaker | New Brunswick Scientific | |
| SpectraMax M5 microplate reader | Molecular Devices | |
| Avanti JXN-26 | Beckman Coulter | |
| JLA-8.1000 Fixed-Angle Aluminium rotor | Beckman Coulter | 363688 |
| EmulsiFlex-C5 French Press homogeniser | Avestin | |
| AKTA Pure 25M | Cytiva | |

**Supplementary Table 8: Buffer composition for recombinant tau expression and purification**

| Buffer | Component | Final concentration |
|---|---|---|
| Terrific broth (TB) | Tryptone | 12 g/L |

| | | |
|---|---|---|
| | Yeast extract | 24 g/L |
| | Glycerol | 0.4% v/v |
| 10x TB salt | Potassium phosphate monobasic | 0.17 M |
| | Potassium phosphate dibasic | 0.72 M |
| Lysis (pH 7.6) | HEPES | 25 mM |
| | NaCl | 150 mM |
| | Imidazole | 10 mM |
| | Bond Breaker ™ TCEP | 1 mM |
| | DNase I | 1.68 U/mL |
| | Pierce™ Protease Inhibitor XL Capsules, EDTA-free | 1x |
| HisTrap A1 (pH 7.6) | HEPES | 25 mM |
| | NaCl | 150 mM |
| | Imidazole | 10 mM |
| | Bond Breaker ™ TCEP | 1 mM |
| | Pierce™ Protease Inhibitor XL Capsules, EDTA-free | 1:500 |
| HisTrap A2 (pH 7.6) | HEPES | 25 mM |
| | NaCl | 150 mM |
| | Imidazole | 10 mM |
| | Bond Breaker ™ TCEP | 1 mM |
| | Triton™X-114 | 0.1% |
| HisTrap B (pH 7.6) | HEPES | 25 mM |
| | NaCl | 150 mM |
| | Imidazole | 500 mM |
| | Bond Breaker ™ TCEP | 1 mM |
| | Pierce™ Protease Inhibitor XL Capsules, EDTA-free | 1:500 |
| Dialysis (pH 7.6) | HEPES | 25 mM |
| | NaCl | 100 mM |
| | β-mercaptoethanol | 5 mM |
| HisTrap Rebind C (pH 7.1) | HEPES | 25 mM |
| | NaCl | 100 mM |
| | Imidazole | 10 mM |
| | β-mercaptoethanol | 5 mM |
| HisTrap Rebind D (pH 7.1) | HEPES | 25 mM |
| | NaCl | 100 mM |
| | Imidazole | 500 mM |
| | β-mercaptoethanol | 5 mM |
| CIEX E (pH 7.1) | HEPES | 25 mM |
| | NaCl | 100 mM |
| | β-mercaptoethanol | 5 mM |
| CIEX F | HEPES | 25 mM |
| | NaCl | 1 M |

| | | |
|---|---|---|
| (pH 7.1) | β-mercaptoethanol | 5 mM |

**Supplementary Table 9: Endotoxin removal from recombinant tau in published studies**

| Publication | Recombinant tau source | Endotoxin removal methods used | Endotoxin detection test | Test results provided |
|---|---|---|---|---|
| Asai et al., 2015 | rPeptide (T-1001-1), *E. coli* purification | N | N | N |
| Zhu et al., 2022 | AnaSpec (AS-55556), *E. coli* purification | N | N | N |
| Udeochu et al., 2023 | non-disclosed | N | N | N |
| Perea et al., 2022 | ClearColi purification | Y | Y | N/A |
| Funk et al., 2015 | *E. coli* purification | N | N | N |
| Ising et al., 2019 | *E. coli* purification | N | Y | N |
| Stancu et al., 2019 | *E. coli* purification | N | N | N |
| Pampuscenko et al., 2020 | *E. coli* purification | N | Y | Y (0.001-0.007% w/w LPS/tau) |
| Zilkova et al., 2020 | *E. coli* purification | Y | N | N |
| Das et al., 2020 | *E. coli* purification | N | N | N |
| Pampuscenko et al., 2021 | *E. coli* purification | N | Y | Y (0.001-0.007% w/w LPS/tau) |
| Jin et al., 2021 | *E. coli* purification | N | N | N |
| Wang et al., 2022 | *E. coli* purification | N | Y | Y (<1 EU/mL in working concentration) |
| Chinnathambi and Das, 2023 | *E. coli* purification | N | N | N |
| Falkon et al., 2024 | StressMarq Biosciences (SPR-327, SPR-329) E. coli BL21 | Y | Y | Y (<0.5 EU/mL at 2 mg/mL) |
| Bolos et al., 2017 | *E. coli* purification | N | N | N |

| | | | | |
|---|---|---|---|---|
| Chidambaram et al., 2024 (bioRxiv) | *E. coli* BL21 purification | N | N | N |
| Crotti et al., 2019 | *E. coli* BL21 Star (DE3) pLysS purification | N | N | N |
| Funk, 2024 | *E. coli* purification | N | N | N |
| Kovac et al., 2011 | *E. coli* purification | N | Y | Y (<0.001 EU/mL) |

**Supplementary Data 1.** RNA-seq Normalised Counts.

**Supplementary Data 2.** RNA-seq DESeq2_DEGs.

**Supplementary Data 3.** RNA-seq GOSeq_Enrichment.

**Supplementary Data 4.** List of proteins and DEPs found in cell lysates of iMGL treated with vehicle, rTauM, rTauF or hTau.

**Supplementary Data 5.** Phosphopeptides and DEPs found in cell lysates of vehicle-, rTauM-, rTauF- of hTau-treated iMGL.

**Supplementary Data 6.** Phosphopeptides found in *E.coli*-derived recombinant tau fibrils and pooled human tau.

**Supplementary Data 7.** List of proteins and DEPs found in the conditioned medium from iMGL treated with vehicle, rTauM, rTauF or hTau.

**Supplementary Data 8.** List of proteins and DEPs found in extracellular vesicles released by iMGL treated with vehicle, rTauM, rTauF or hTau.

### Supplementary figures and legends

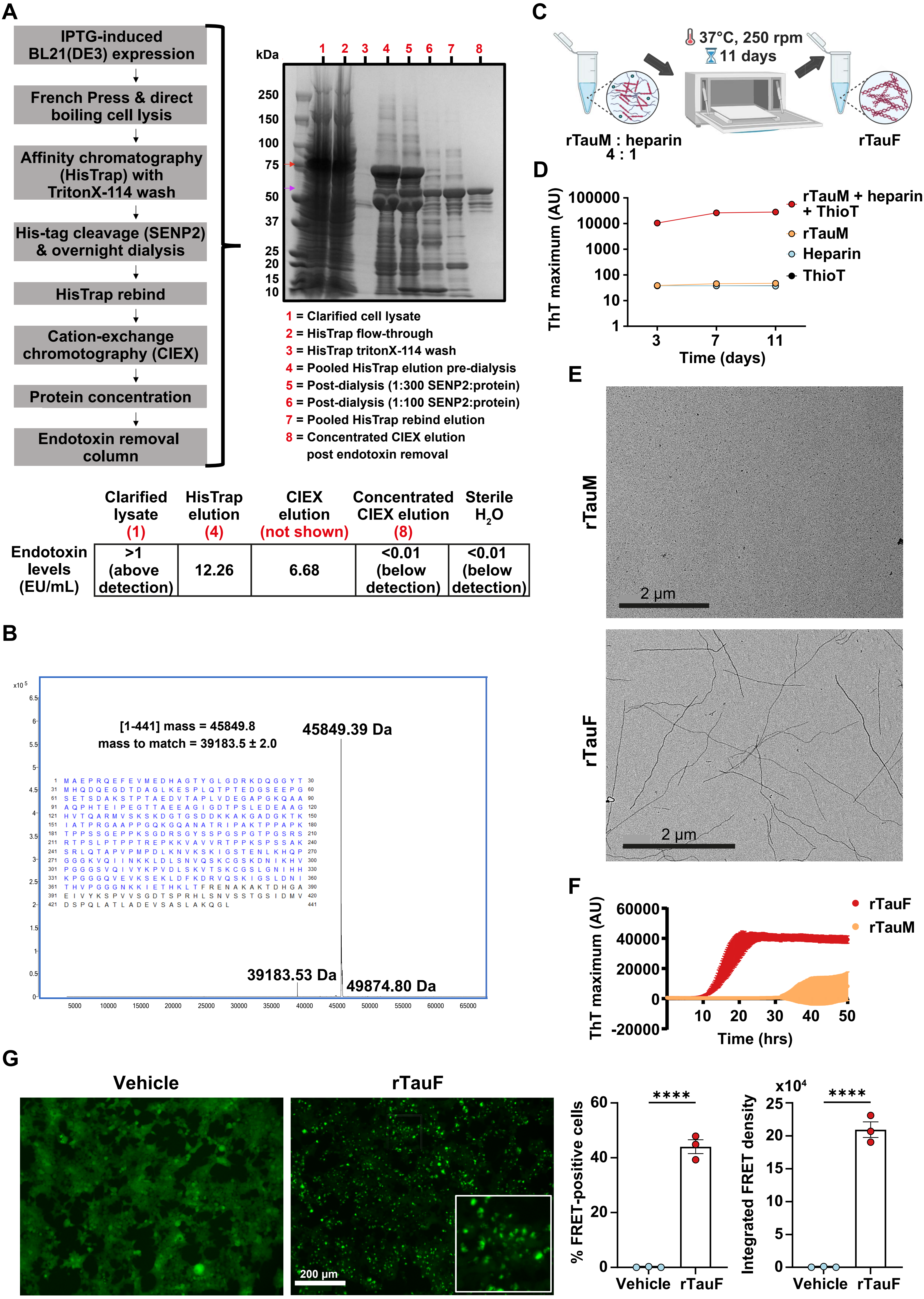

| | Clarified lysate (1) | HisTrap elution (4) | CIEX elution (not shown) | Concentrated CIEX elution (8) | Sterile $H_2O$ |
|---|---|---|---|---|---|
| Endotoxin levels (EU/mL) | >1 (above detection) | 12.26 | 6.68 | <0.01 (below detection) | <0.01 (below detection) |

**Supplementary Figure 1. Endotoxin-free production of recombinant 2N4R tau.**

**A** Purification workflow and SimplyBlue$^{TM}$-stained gel showing eluted protein fractions. Red arrow points to 6xHis_SUMO3 tagged tau product, the purple arrow points to the tag-less tau product post SENP2 proteolytic cleavage. A 6L bacterial culture yielded a total of 9.1 mg of recombinant tau. Endotoxin levels (EU/mL) in individual purified tau fractions were determined by LAL assay. Numbers in red correspond to fractions shown on the blot. Protein levels were normalised to 1 mg/mL in sterile dd$H_2O$. **B** Results of intact protein characterisation by LC/MS. **C** Workflow diagram of heparin-induced *in vitro* rTauM aggregation. **D** ThT assay confirming presence of β-sheet structures in rTauF preparation. **E** Negative stain TEM images of rTauM and rTauF. **F** Results of 4R tau RT-QuIC assay investigating seeding of truncated tau substrate (K11, tau aa residues 244-394) in the presence of rTauM and rTauF. A total of 4 technical replicates were run per condition. Data is presented as mean ± SD. **G** Representative images of tau inclusions in FRET Biosensor HEK cells treated with vehicle or transfected with 50 nM rTauF, and quantification of FRET-positive cells and integrated FRET density. Data is represented as mean ± SEM n = 3 technical replicates. Two-tailed unpaired Student t-test was used. ****$P < 0.0001$.

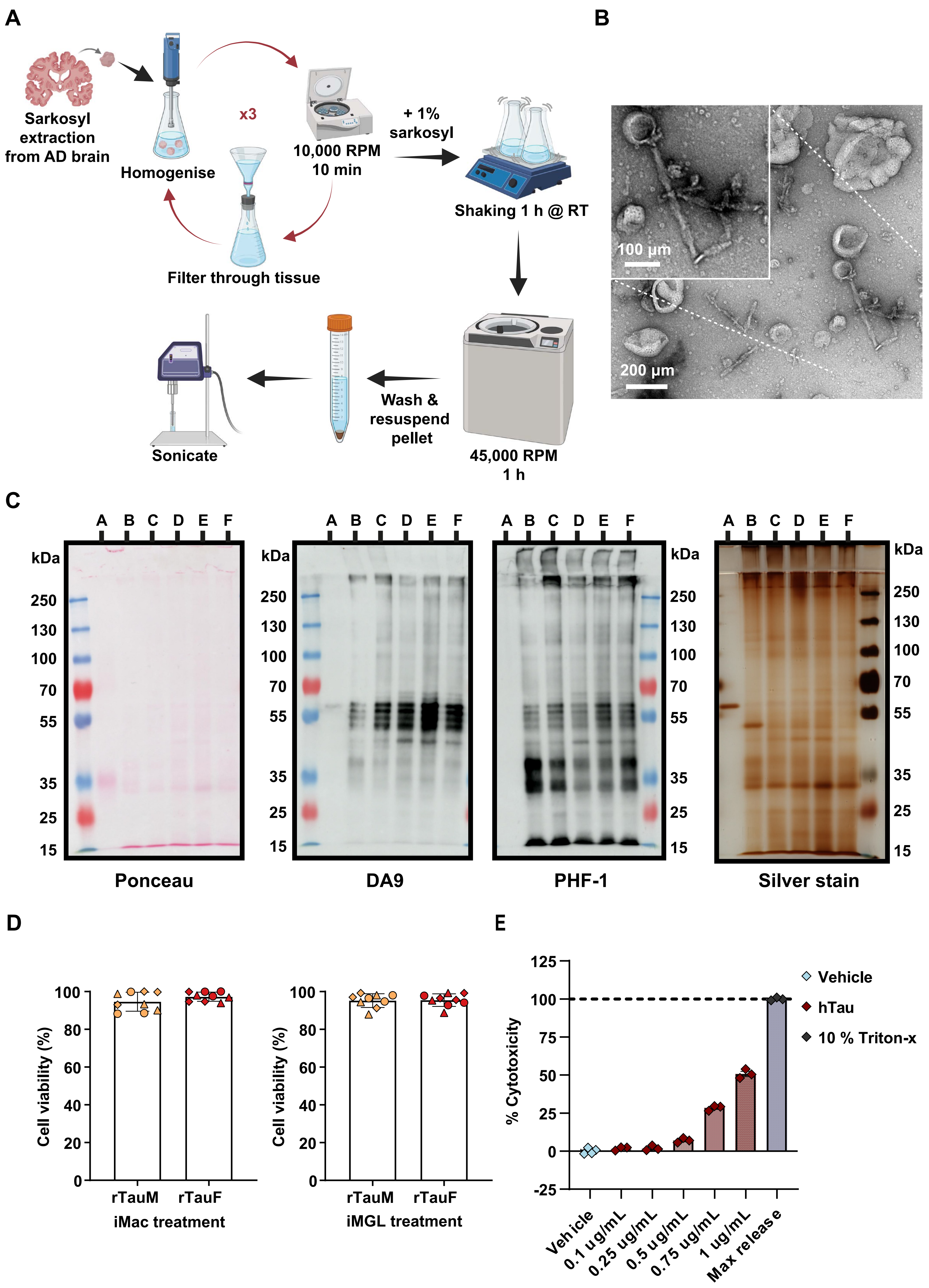

A
Sarkosyl extraction from AD brain
Homogenise
x3
10,000 RPM 10 min
Filter through tissue
+ 1% sarkosyl
Shaking 1 h @ RT
45,000 RPM 1 h
Wash & resuspend pellet
Sonicate
B
100 µm
200 µm
C
A B C D E F
kDa
250
130
100
70
55
35
25
15
Ponceau
DA9
PHF-1
Silver stain
D
Cell viability (%)
rTauM
rTauF
iMac treatment
iMGL treatment
E
% Cytotoxicity
Vehicle
hTau
10 % Triton-x
Vehicle
0.1 ug/mL
0.25 ug/mL
0.5 ug/mL
0.75 ug/mL
1 ug/mL
Max release

**Supplementary Figure 2. Purification of hTau from AD brain samples.**

**A** Workflow diagram of sarkosyl-insoluble tau PHFs (hTau) extraction from human AD brain. **B** Negative stain TEM image of extracted hTau preparation with blow-up of a representative fibril with visible helical twists. **C** Quality control western blots for hTau preparations. Markers (left to right): Ponceau (total protein), DA9 (total tau, aa102–140), PHF-1 (pTau S396/S404), silver stain (aggregated tau). Samples (same order for all blots): A: 2N4R/Tau 441 (control), B: Tau seed prep #1, C: Tau seed prep #1 flocculent, D: Tau seed prep #2, E: Tau seed prep #3, F: Pooled tau seed (from preparation B, D and E). **D** Resazurin cell viability assay in iMac/iMGL after 24 h incubation with 4 µg/mL rTauM or rTauF relative to unstimulated control.Data is represented as mean ±SEM of n = 3 in 3 control cell lines. **E** LDH cytotoxicity assay conducted on iMGL supernatants after 24 h incubation with increasing amounts of hTau. Percentage cytotoxicity calculated in relation to the maximum LDH measure from cells treated with 10% Triton-x, vehicle represents untreated iMGL. Data is represented as mean ±SEM of n = 3-4 technical replicates from one control cell line.

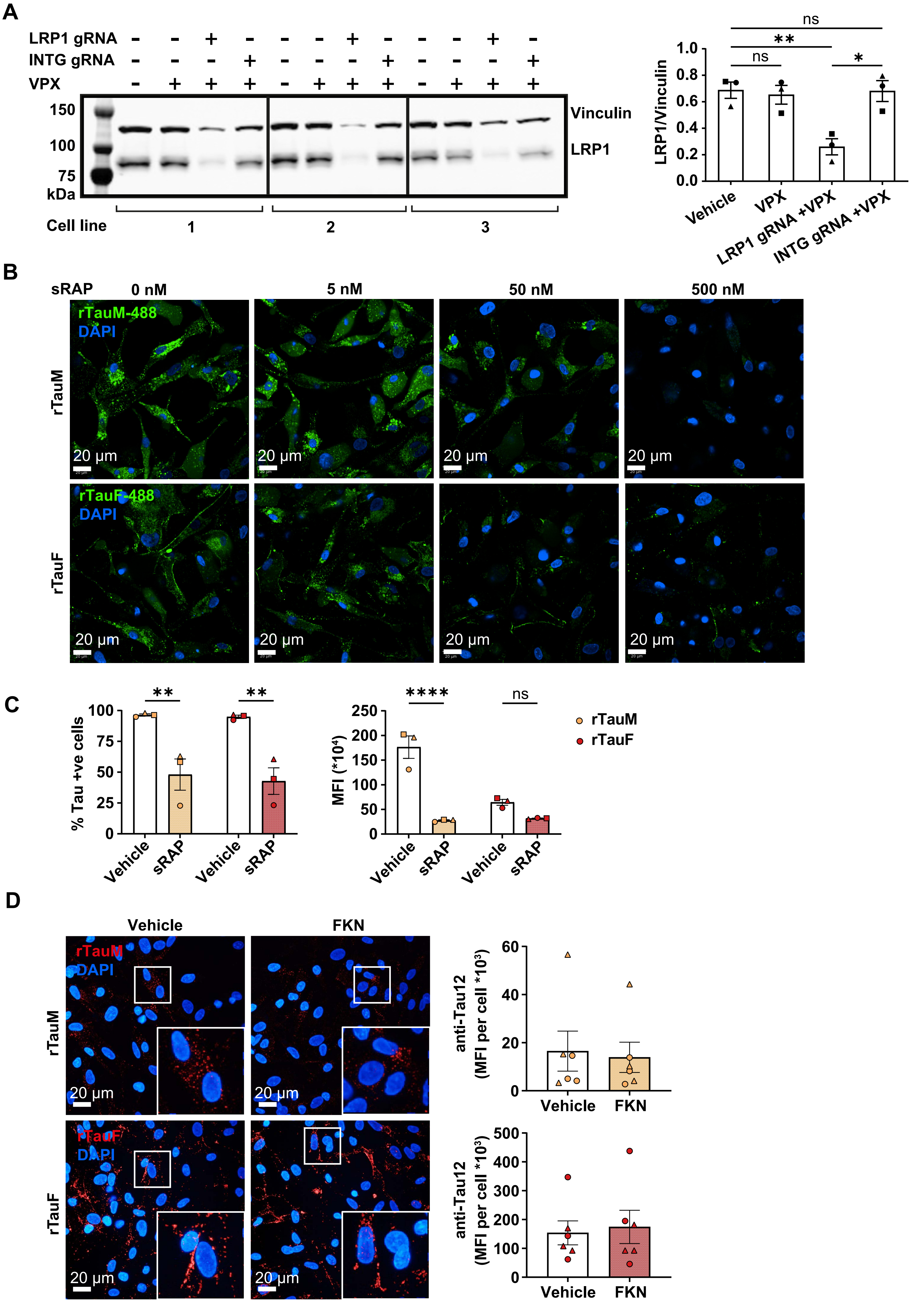

A
LRP1 gRNA
INTG gRNA
VPX
150
100
75
kDa
Vinculin
LRP1
Cell line
1
2
3
ns
**
ns
*
LRP1/Vinculin
Vehicle
VPX
LRP1 gRNA +VPX
INTG gRNA +VPX
B
sRAP
0 nM
5 nM
50 nM
500 nM
rTauM-488
DAPI
rTauM
rTauF-488
DAPI
rTauF
20 µm
C
% Tau +ve cells
MFI (*10⁴)
****
ns
rTauM
rTauF
Vehicle
sRAP
D
Vehicle
FKN
rTauM
DAPI
rTauF
DAPI
anti-Tau12
(MFI per cell *10³)

**Supplementary Figure 3. Tau entry in iMGL is LRP1- but not CX3CR1-dependent.**

**A** Western blot confirmation of CRISPR-Cas9 mediated LRP1 KD in iMac. LRP1-targeting versus Intergenic region-targeting (INTG) gRNAs, and transduction enhancer VPX (accessory protein delivered via virus-like particles to facilitate the degradation of SAMHD1, resulting in increasing the availability of dNTPs for viral reverse transcription). Data is displayed as mean ± SEM of n = 1 in 3 control cell lines. One-way ANOVA with Tukey's multiple comparison test was used. $^{*}P < 0.05$, $^{**}P < 0.01$. **B** A 2 h incubation of iMGL with sRAP and DyLight 488-conjugated rTauM or rTauF reduces tau uptake in a dose-dependent manner. **C** Flow cytometry analysis of 2 h DyLight 488-conjugated rTauM and rTauF uptake in the presence of 500 nM sRAP. Data is shown as mean ± SEM of n = 1 in 3 control cell lines. Two-way ANOVA with Bonferroni's multiple comparison test was used. $^{**}P < 0.01$, $^{****}P < 0.0001$. **D** Representative confocal microscopy images and quantification of Tau12 mean integrated density in iMGL treated with 5 ng/mL FKN for 1 h and challenged with 1 µg/mL rTauM or rTauF for 2 h. Data is represented as mean ± SEM of n = 3 in 2 control cell lines. Two-tailed unpaired Student t test.

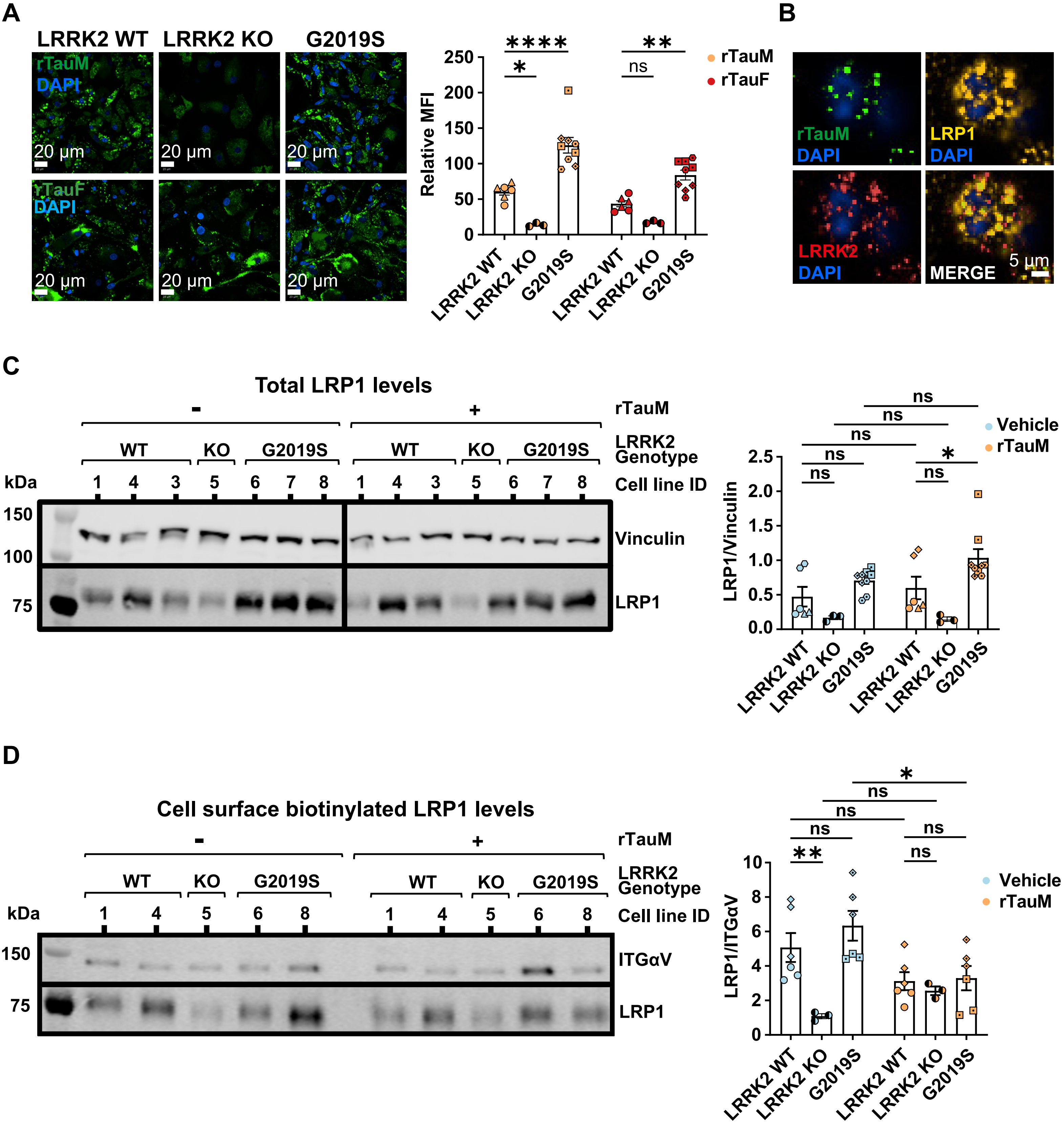
A
LRRK2 WT
LRRK2 KO
G2019S
rTauM
DAPI
rTauF
DAPI
20 μm
Relative MFI
250
200
150
100
50
0
****
*
**
ns
rTauM
rTauF
LRRK2 WT
LRRK2 KO
G2019S
B
rTauM
DAPI
LRP1
DAPI
LRRK2
DAPI
MERGE
5 μm
C
Total LRP1 levels
-
+
rTauM
WT
KO
G2019S
LRRK2 Genotype
kDa
1 4 3 5 6 7 8 1 4 3 5 6 7 8
Cell line ID
150
100
75
Vinculin
LRP1
LRP1/Vinculin
2.5
2.0
1.5
1.0
0.5
0.0
ns
*
Vehicle
rTauM
D
Cell surface biotinylated LRP1 levels
1 4 5 6 8 1 4 5 6 8
ITGαV
LRP1
LRP1/ITGαV
10
8
6
4
2
0
**

**Supplementary Figure 4. LRRK2 genotype influences uptake of DyLight 488-conjugated rTauM and rTauF.**

**A** Representative confocal microscopy images (left) and flow cytometry analysis (right) after 6 h incubation with DyLight 488-conjugated rTauM or rTauF in iMac. MFI of internalised rTauM/rTauF is quantified relative to individual cell-line autofluorescence. Data is shown as mean ± SEM of n = 3 in 2 LRRK2 WT lines, 1 LRRK2 KO line, and 3 G2019S LRRK2 cell lines. Two-way ANOVA with Dunnett's multiple comparisons test $*P < 0.05$, $**P < 0.01$, $****P < 0.0001$. **B** Representative confocal microscopy images showing overlap of LRRK2 puncta with LRP1-rTauM complexes in iMac after 2 h of incubation. **C** Western blot analysis of total and **D** biotinylated surface LRP1 levels in iMac at baseline and after 2 h of rTauM incubation. Data is represented as mean ± SEM of n = 2 in 3 LRRK2 WT, 1 LRRK2 KO, 3 G2019S cell lines for the total LRP1 protein levels, and n = 3 in 2 LRRK2 WT, 1 LRRK2 KO, 2 G2019S cell lines for the surface LRP1 protein levels. Two-way ANOVA with Šídák's multiple comparisons test was used. $*P < 0.05$, $**P < 0.01$.

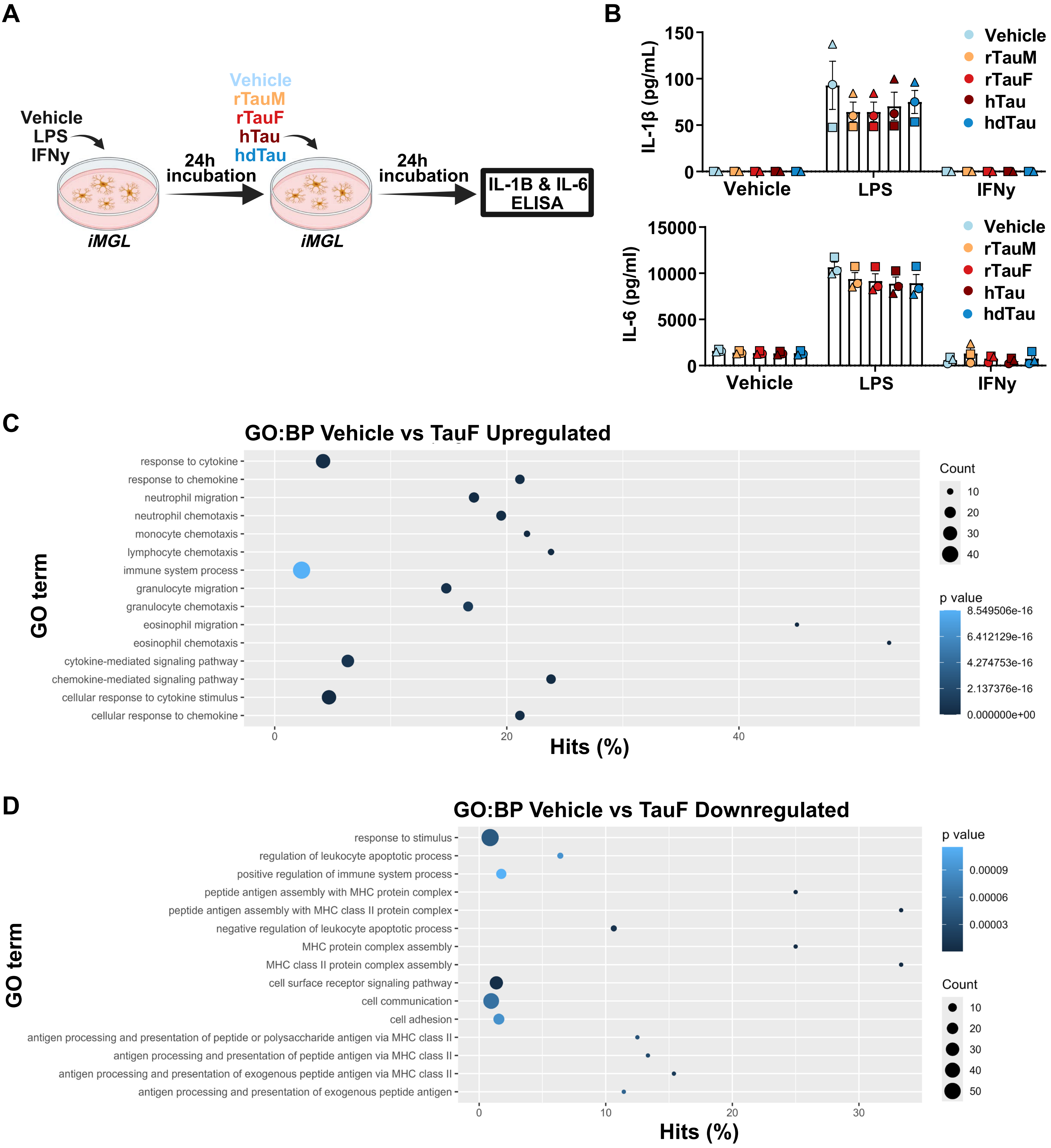
A
Vehicle
LPS
IFNy
iMGL
24h
incubation
Vehicle
rTauM
rTauF
hTau
hdTau
iMGL
24h
incubation
IL-1B & IL-6
ELISA
B
IL-1β (pg/mL)
150
100
50
0
Vehicle
LPS
IFNy
Vehicle
rTauM
rTauF
hTau
hdTau
IL-6 (pg/ml)
15000
10000
5000
0
Vehicle
LPS
IFNy
Vehicle
rTauM
rTauF
hTau
hdTau
C
GO:BP Vehicle vs TauF Upregulated
response to cytokine
response to chemokine
neutrophil migration
neutrophil chemotaxis
monocyte chemotaxis
lymphocyte chemotaxis
immune system process
granulocyte migration
granulocyte chemotaxis
eosinophil migration
eosinophil chemotaxis
cytokine-mediated signaling pathway
chemokine-mediated signaling pathway
cellular response to cytokine stimulus
cellular response to chemokine
GO term
0
20
40
Hits (%)
Count
10
20
30
40
p value
8.549506e-16
6.412129e-16
4.274753e-16
2.137376e-16
0.000000e+00
D
GO:BP Vehicle vs TauF Downregulated
response to stimulus
regulation of leukocyte apoptotic process
positive regulation of immune system process
peptide antigen assembly with MHC protein complex
peptide antigen assembly with MHC class II protein complex
negative regulation of leukocyte apoptotic process
MHC protein complex assembly
MHC class II protein complex assembly
cell surface receptor signaling pathway
cell communication
cell adhesion
antigen processing and presentation of peptide or polysaccharide antigen via MHC class II
antigen processing and presentation of peptide antigen via MHC class II
antigen processing and presentation of exogenous peptide antigen via MHC class II
antigen processing and presentation of exogenous peptide antigen
GO term
0
10
20
30
Hits (%)
p value
0.00009
0.00006
0.00003
Count
10
20
30
40
50

**Supplementary Figure 5. Cytokine and Supplementary RNA-seq analyses.**

**A** Schematic of experimental workflow. iMGL were pre-incubated for 24 h with inflammatory stimulants lipopolysaccharide (LPS) or interferon gamma (IFNy) and subsequently treated with rTau or hTau for 24 h, after which supernatants were collected. **B** IL-1β and IL-6 detected in iMGL supernatants by ELISA, n = 1 in 3 control cell lines. **C** Up-regulated Gene Ontology terms for vehicle vs rTauF. **D** Down-regulated Gene Ontology terms for vehicle vs rTauF.

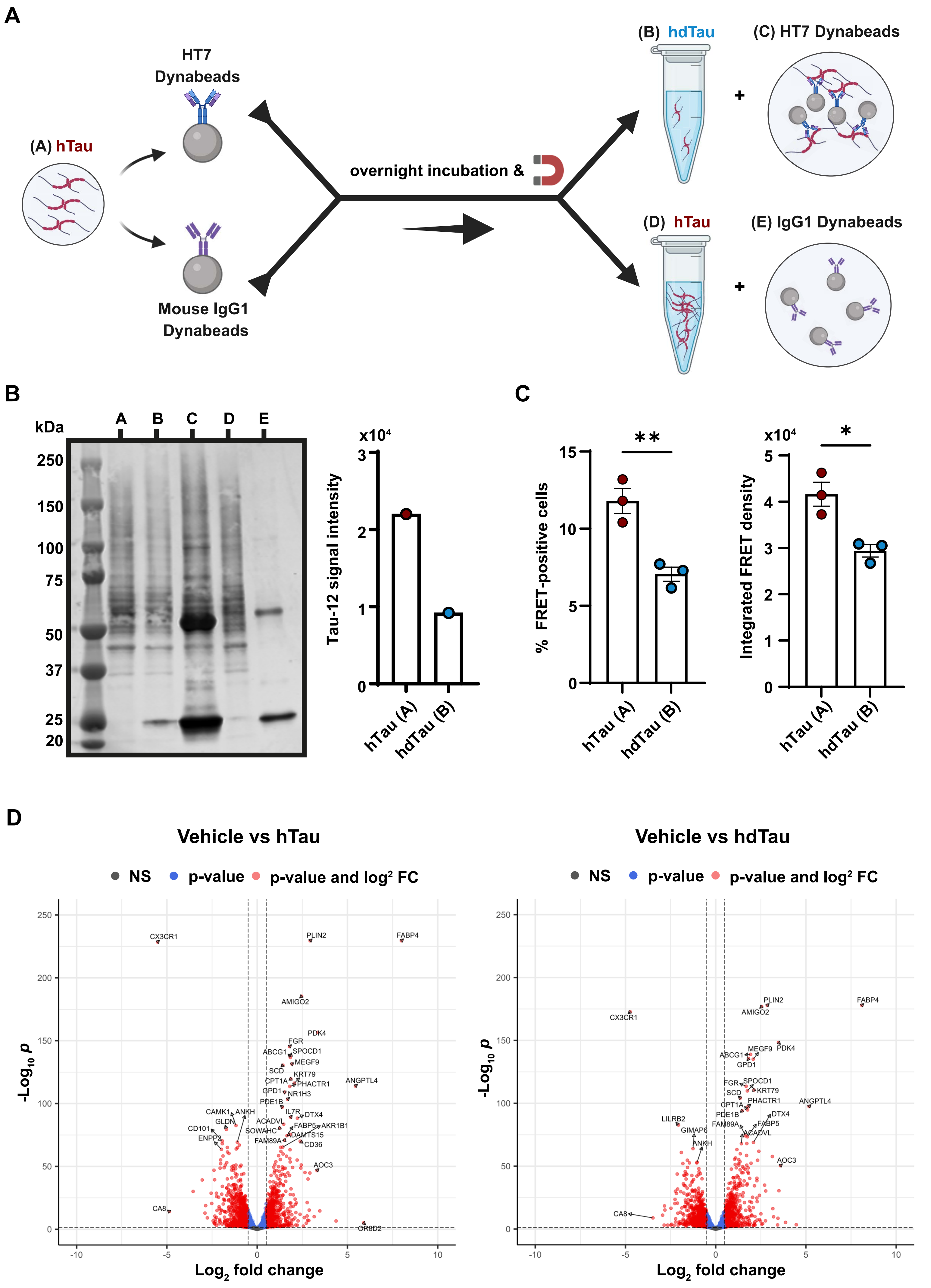
A
(A) hTau
HT7
Dynabeads
Mouse IgG1
Dynabeads
overnight incubation &
(B) hdTau
(C) HT7 Dynabeads
(D) hTau
(E) IgG1 Dynabeads
B
kDa
A
B
C
D
E
250
150
100
75
50
37
25
20
x10⁴
Tau-12 signal intensity
hTau (A)
hdTau (B)
C
% FRET-positive cells
Integrated FRET density
**
*
D
Vehicle vs hTau
Vehicle vs hdTau
NS
p-value
p-value and log² FC
-Log10 p
Log2 fold change
CX3CR1
PLIN2
FABP4
AMIGO2
PDK4
FGR
ABCG1
SPOCD1
MEGF9
SCD
KRT79
CPT1A
PHACTR1
ANGPTL4
GPD1
NR1H3
PDE1B
CAMK1
ANKH
IL7R
DTX4
GLDN
ACADVL
CD101
SOWAHC
FABP5
AKR1B1
ENPP2
ADAMTS15
FAM89A
CD36
AOC3
CA8
OR8D2
LILRB2
GIMAP6

**Supplementary Figure 6. hTau immunodepletion.**

**A** Schematic diagram of tau and control IgG immunodepletion from hTau preparation. **B** Western blot confirmation of tau reduction in hdTau preparation (left). Individual lane labels correspond to the schematic labels from **A**. Quantification of total tau signal reduction from lanes A and B (right). **C** Percentage of FRET-positive cells and integrated FRET density in Tau RD P301S FRET Biosensor HEK cells transfected with 1 ng of hTau or hdTau. Data is represented as mean ± SEM, n = 3 technical replicates. **P* < 0.05, ***P* < 0.01. Two-tailed unpaired Student t-test was used. **D** Volcano plots for vehicle vs hTau (left) and vehicle vs tau-depleted brain sample hdTau (right).

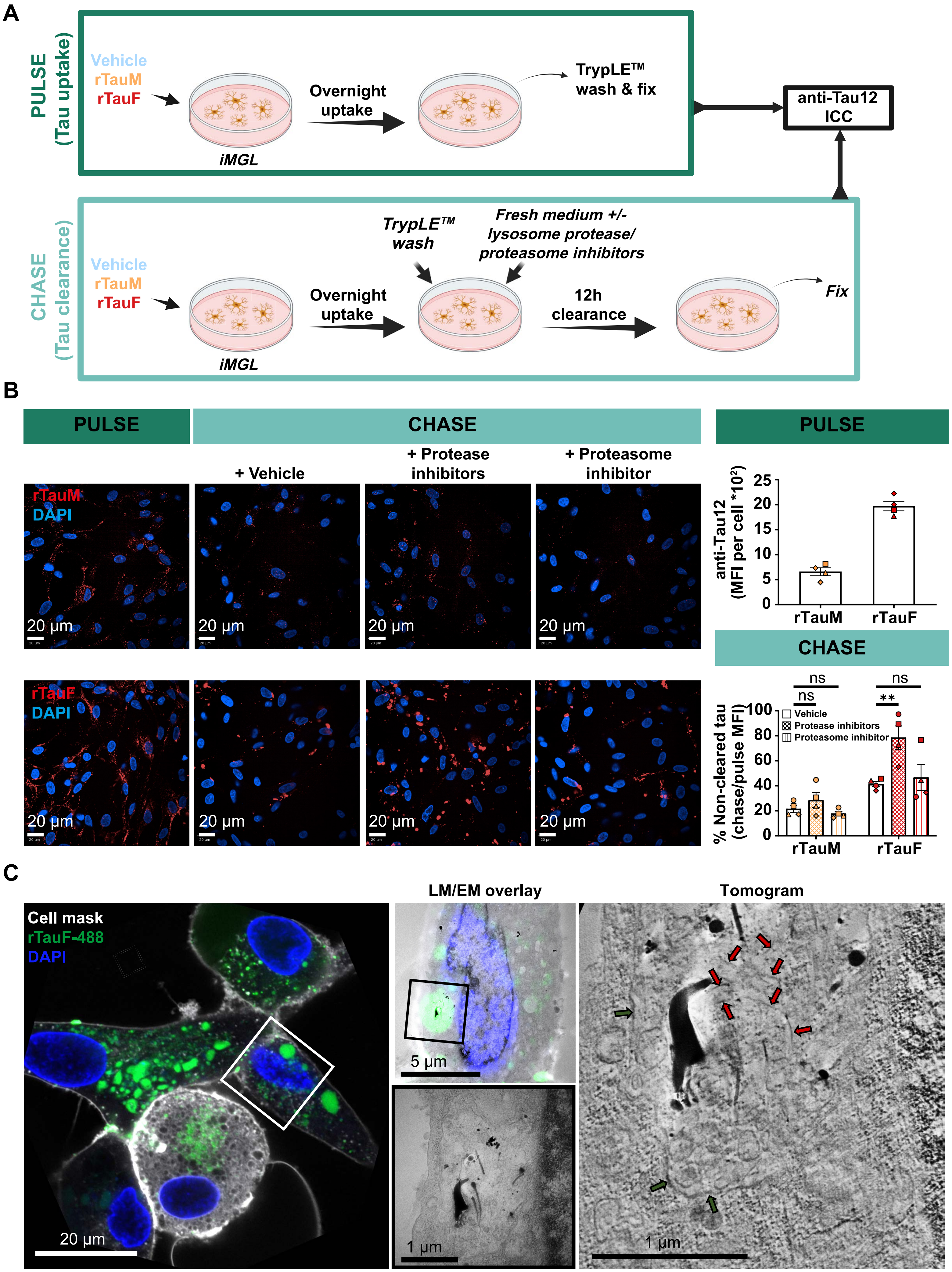
A
PULSE (Tau uptake)
Vehicle
rTauM
rTauF
Overnight uptake
iMGL
TrypLE™ wash & fix
anti-Tau12 ICC
CHASE (Tau clearance)
Vehicle
rTauM
rTauF
Overnight uptake
iMGL
TrypLE™ wash
Fresh medium +/- lysosome protease/ proteasome inhibitors
12h clearance
Fix
B
PULSE
CHASE
+ Vehicle
+ Protease inhibitors
+ Proteasome inhibitor
rTauM
DAPI
rTauF
DAPI
20 µm
PULSE
anti-Tau12 (MFI per cell *10²)
25
20
15
10
5
0
rTauM
rTauF
CHASE
ns
ns
ns
**
% Non-cleared tau (chase/pulse MFI)
100
80
60
40
20
0
Vehicle
Protease inhibitors
Proteasome inhibitor
rTauM
rTauF
C
Cell mask
rTauF-488
DAPI
20 µm
LM/EM overlay
5 µm
1 µm
Tomogram
1 µm

**Supplementary Figure 7. Tau degradation pathways and further correlative light and electron microscopy of iMGL treated with Dylight-488-labelled rTauF.**

**A** Treatment schematic. Pulse phase involved only overnight incubation with rTauM/rTauF and enzymatic wash to assess baseline uptake of tau, whereas for the chase conditions, iMGL were incubated with either protease or proteasome inhibitors during the 12 h clearance phase. **B** Representative confocal microscopy images (left) and quantification of Tau12 mean integrated density (right) showing the effect of lysosomal protease inhibition or proteasome inhibition on rTauM and rTauF clearance in iMGL. Data is represented as mean ± SEM of n = 1-2 in 3 control cell lines (pulse) or n = 1 in 4 control cell lines (chase). Two-way ANOVA with Dunnett's multiple comparisons test $^{**}P < 0.01$. **C** Correlative light and electron microscopy (CLEM) images overlay (left) of iMGL incubated overnight (16 h) with Dylight 488-labelled rTauF. Tomogram z-projection (average intensity, right) of the same region containing rTauF (red arrows) within a partial enclosing membrane (green arrows).

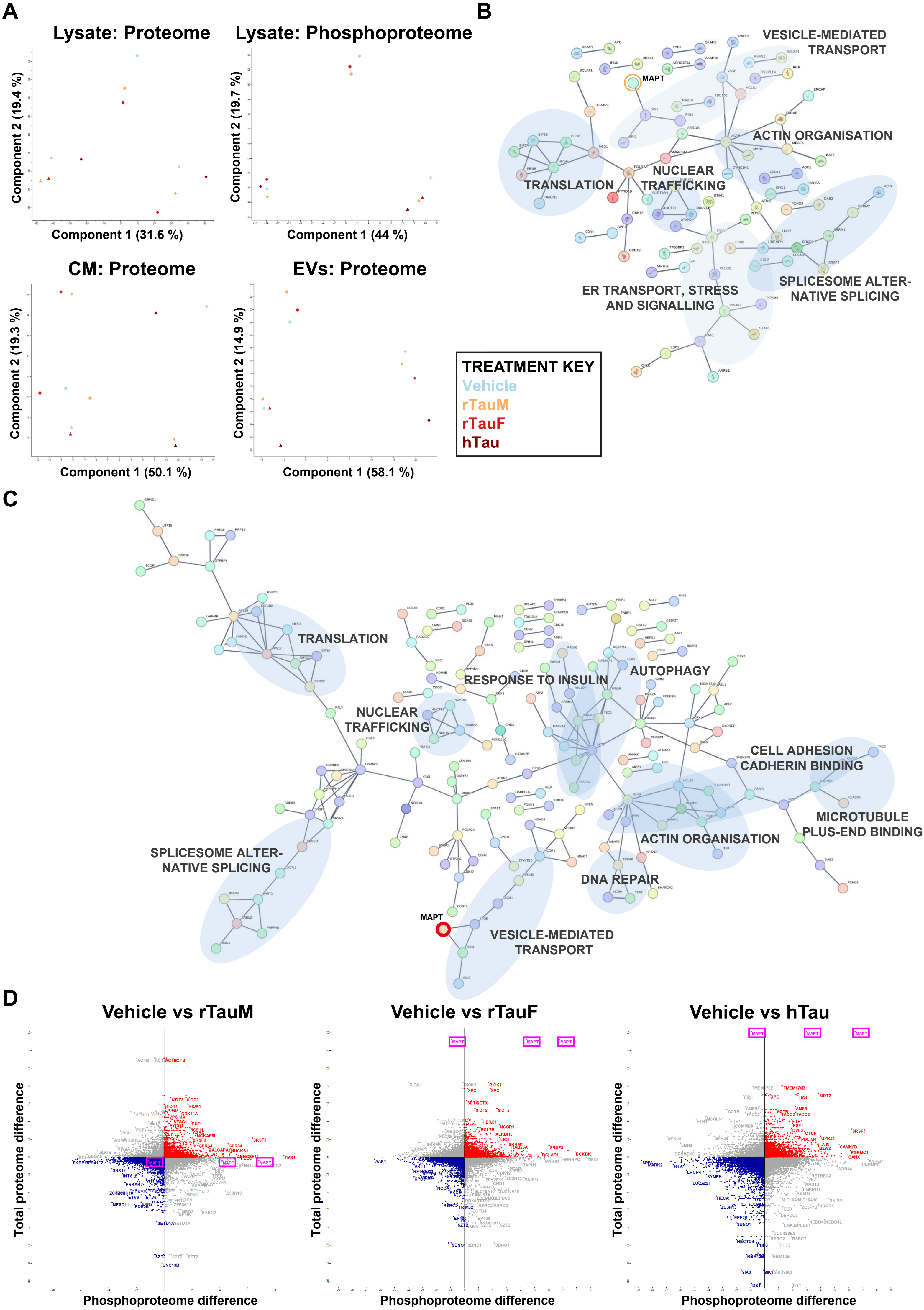
A
Lysate: Proteome
Component 2 (19.4 %)
Component 1 (31.6 %)
Lysate: Phosphoproteome
Component 2 (19.7 %)
Component 1 (44 %)
CM: Proteome
Component 2 (19.3 %)
Component 1 (50.1 %)
EVs: Proteome
Component 2 (14.9 %)
Component 1 (58.1 %)
B
VESICLE-MEDIATED TRANSPORT
MAPT
ACTIN ORGANISATION
TRANSLATION
NUCLEAR TRAFFICKING
SPLICESOME ALTER-NATIVE SPLICING
ER TRANSPORT, STRESS AND SIGNALLING
TREATMENT KEY
Vehicle
rTauM
rTauF
hTau
C
TRANSLATION
RESPONSE TO INSULIN
AUTOPHAGY
NUCLEAR TRAFFICKING
CELL ADHESION CADHERIN BINDING
MICROTUBULE PLUS-END BINDING
ACTIN ORGANISATION
DNA REPAIR
SPLICESOME ALTER-NATIVE SPLICING
MAPT
VESICLE-MEDIATED TRANSPORT
D
Vehicle vs rTauM
Vehicle vs rTauF
Vehicle vs hTau
Total proteome difference
Phosphoproteome difference
MAPT

**Supplementary Figure 8. Proteomics and phosphoproteomics of iMGL.**

**A** Principal Component Analysis plots of all iMGL samples analysed by proteomics (cell lysate, CM and EVs) and phosphoproteomics (cell lysates). **B-C** STRING analysis of differentially expressed phosphopeptides in iMGL treated with rTauM **(B)** or hTau **(C)**. MAPT highlighted with orange (rTauM, B) or red circle (hTau, C). Only high-confidence interactions (interaction score > 0.9) are displayed. **D** Volcano plots of phosphopeptides versus total proteins identified in iMGL treated with rTauM, rTauF, or hTau compared to vehicle. MAPT peptides highlighted with pink boxes.

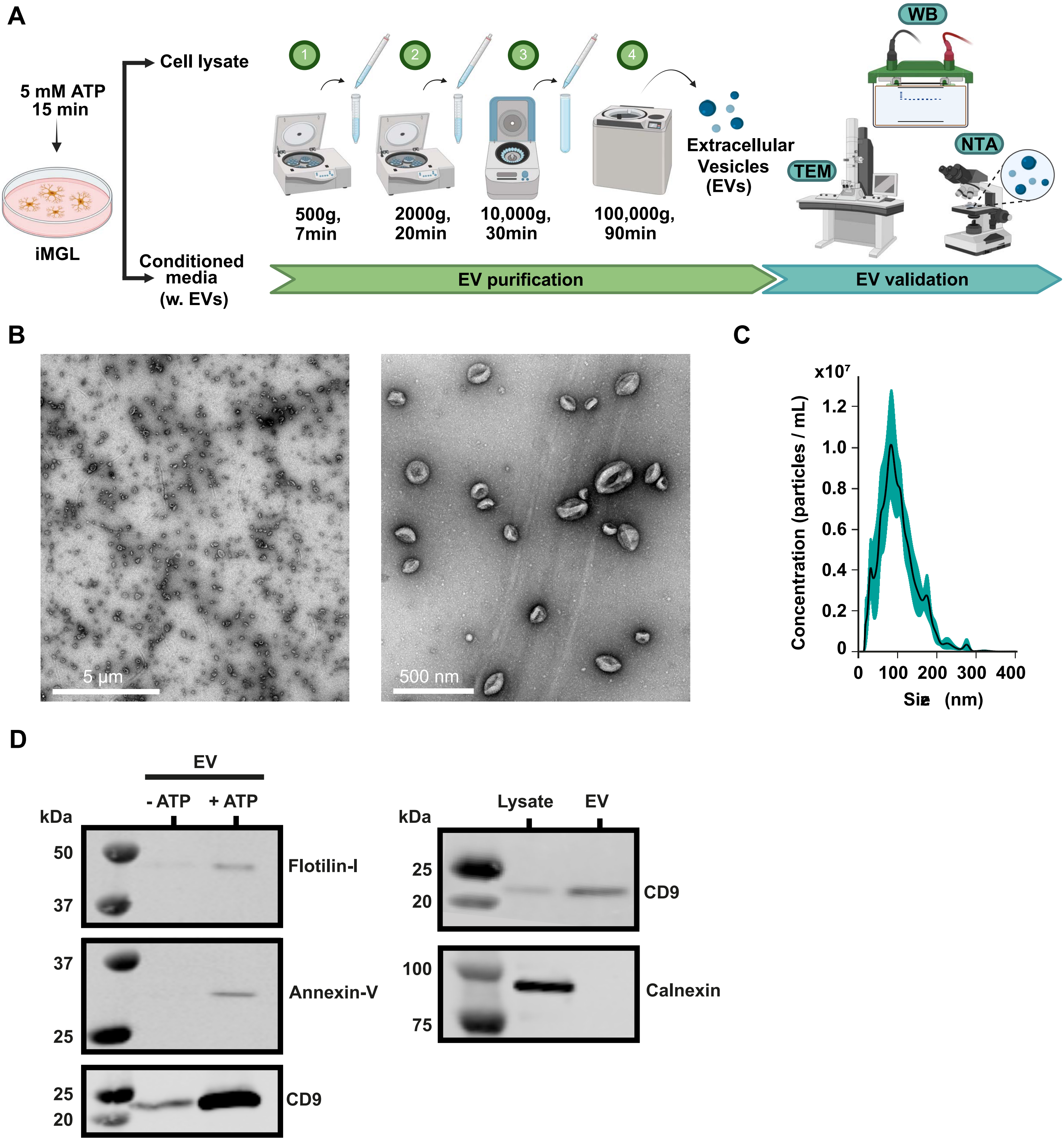
A
5 mM ATP
15 min
iMGL
Cell lysate
Conditioned media (w. EVs)
1
2
3
4
500g, 7min
2000g, 20min
10,000g, 30min
100,000g, 90min
Extracellular Vesicles (EVs)
WB
TEM
NTA
EV purification
EV validation
B
5 µm
500 nm
C
x10⁷
Concentration (particles / mL)
1.2
1.0
0.8
0.6
0.4
0.2
0
0
100
200
300
400
Size (nm)
D
EV
- ATP
+ ATP
kDa
50
37
Flotilin-I
37
25
Annexin-V
25
20
CD9
Lysate
EV
kDa
25
20
CD9
100
75
Calnexin

**Supplementary Figure 9. Validation of the EV purification process.**

**A** Experimental workflow illustrating the purification of EVs from iMGL cells treated with 15 mM ATP (15 min) by a series of centrifugation steps of increasing speed and validated with both transmission electron microscopy (TEM), nanoparticle tracking analysis (NTA) and western blotting (WB). **B** Representative TEM images at different magnifications illustrating shape and sizes of purified EVs. **C** Quantification of particle sizes via NTA. **D** Enriched expression of EV markers Flotilin-I, Annexin-V and CD9 as a result of ATP stimulation (left). Validation of EVs with positive control marker CD9 and negative control marker calnexin, only expressed in cell lysate (right).

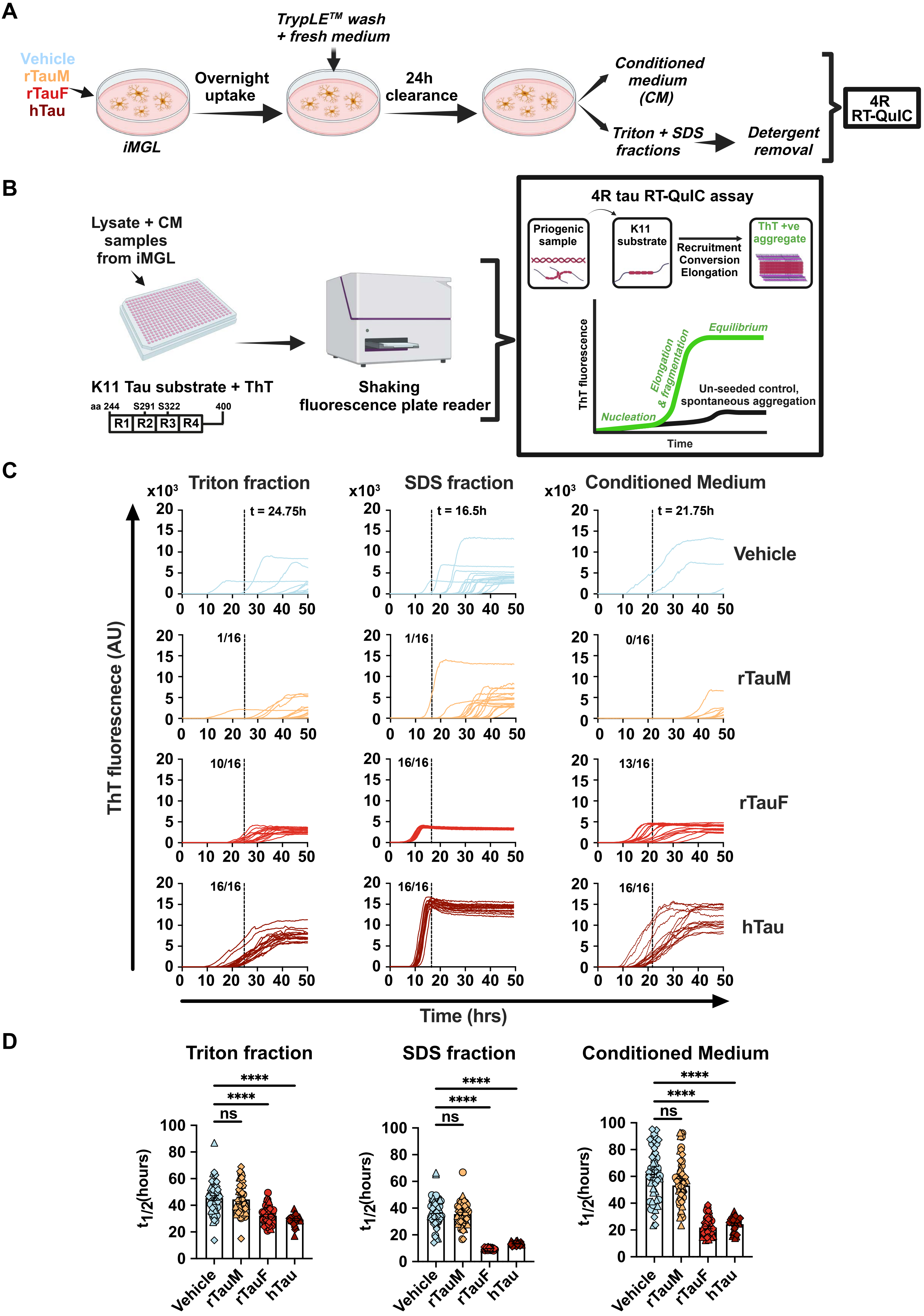
A
Vehicle
rTauM
rTauF
hTau
iMGL
Overnight uptake
TrypLE™ wash + fresh medium
24h clearance
Conditioned medium (CM)
Triton + SDS fractions
Detergent removal
4R RT-QuIC
B
Lysate + CM samples from iMGL
K11 Tau substrate + ThT
aa 244
S291 S322
400
R1 R2 R3 R4
Shaking fluorescence plate reader
4R tau RT-QuIC assay
Priogenic sample
K11 substrate
Recruitment Conversion Elongation
ThT +ve aggregate
ThT fluorescence
Equilibrium
Elongation & fragmentation
Nucleation
Un-seeded control, spontaneous aggregation
Time
C
Triton fraction
SDS fraction
Conditioned Medium
x10³
t = 24.75h
t = 16.5h
t = 21.75h
Vehicle
rTauM
rTauF
hTau
1/16
1/16
0/16
10/16
16/16
13/16
16/16
16/16
16/16
ThT fluorescnece (AU)
Time (hrs)
D
Triton fraction
SDS fraction
Conditioned Medium
****
****
ns
t1/2(hours)
Vehicle
rTauM
rTauF
hTau

**Supplementary Figure 10. Seeding capacity of tau taken up by iMGL, assessed with RT-QuIC.**

**A** Experimental workflow. **B** 4R tau RT-QuIC workflow and illustration of expected curves from the assay and their interpretation. **C** Representative seeding activity of samples collected from a control iMGL cell line. Total protein levels in cell lysate fractions and media were normalised to 1 mg/mL. Triton and SDS lysate fractions were applied to the reaction buffer at 1:10,000 dilution, conditioned media at 1:100, to account for tau concentration differences. Dashed line is set at a kinetic cut-off between templated substrate seeding vs spontaneous aggregation. A reaction was considered seeded upon reaching ThT fluorescence threshold > 100 * SD baseline fluorescence[1-4]. Numbers refer to the fraction of the 16 replicates for a given sample type that reaches the kinetic cut-off. **D** t1/2 calculated from raw RT-QuIC ThT reactions as sigmoidal dose-response, variable slope logEC50. Data is represented as mean ± SEM of n = 1 in 3 control cell lines and 16 technical replicates per sample. Two-way ANOVA with Tukey's multiple comparisons test. *$P < 0.05$, **$P < 0.01$, ****$P < 0.0001$.

**Supplementary Video 1.** 3D reconstruction (tomogram) of internalised Dylight 488-labelled rTauF shown in Fig. 5C.

**Supplementary Video 2.** 3D reconstruction (tomogram) of internalised Dylight 488-labelled rTauF shown in Fig. 5C with segmentation model overlay. rTauF (red) and membranous structures (yellow) within a partial enclosing membrane (green) are shown.
